# Simulations reveal challenges to artificial community selection and possible strategies for success

**DOI:** 10.1101/264689

**Authors:** Li Xie, Alex E. Yuan, Wenying Shou

## Abstract

Multi-species microbial communities often display “community functions” arising from interactions of member species. Interactions are often difficult to decipher, making it challenging to design communities with desired functions. Alternatively, similar to artificial selection for individuals in agriculture and industry, one could repeatedly choose communities with the highest community functions to reproduce by randomly partitioning each into multiple “Newborn” communities for the next cycle. However, previous efforts in selecting complex communities have generated mixed outcomes that are difficult to interpret. To understand how to effectively enact community selection, we simulated community selection to improve a community function that requires two species and imposes a fitness cost on one or both species. Our simulations predict that improvement could be easily stalled unless various aspects of selection, including promoting species coexistence, suppressing non-contributors, adopting a “bet-hedging” strategy when choosing communities to reproduce, and reducing stochastic fluctuations in species biomass of Newborn communities, were carefully considered. When these considerations were addressed in experimentally feasible manners, community selection could overcome natural selection to improve community function, and may even force species to evolve growth restraint to achieve species coexistence. Our conclusions hold under various alternative model assumptions, and are thus applicable to a variety of communities.

## Introduction

Multi-species microbial communities often display important *community functions*, defined as biochemical activities not achievable by member species in isolation. For example, a six-species microbial community, but not any member species alone, cleared relapsing *Clostridium difficile* infections in mice [1]. Community functions arise from *interactions* where an individual alters the physiology of another individual. Thus, to improve community functions, one could identify and modify interactions [2, 3]. In reality, this is no trivial task: each species can release tens or more compounds, many of which may influence the partner species in diverse fashions [4, 5, 6, 7]. From this myriad of interactions, one would then need to identify those critical for community function, and modify them by altering species genotypes or the abiotic environment. One could also artificially assemble different combinations of species or genotypes at various ratios to screen for high community function (e.g. [8, 9]). However, some species may not be culturable in isolation, and the number of combinations becomes very large even for a moderate number of species and genotypes especially if various ratios were to be tested.

In an alternative approach, artificial selection of whole communities could be carried out over cycles to improve community function [10, 11, 12, 13, 14] (reviewed in [15, 16, 17]; Figure 1A). A selection cycle starts with a collection of low-density “Newborn” communities with artificially-imposed boundaries (e.g. inside culture tubes). These low-density communities are incubated for a period of time (“maturation”) to form “Adult” communities. During maturation, community members multiply and interact with each other and possibly mutate, and the community function of interest (purple shade) develops. At the end of maturation, desired Adult communities (e.g. darkest purple shade) are chosen to “reproduce” where each is randomly partitioned into multiple Newborn communities to start the next cycle. Superficially, this process may seem straightforward since “one gets what one selects for”. After all, artificial selection on individuals has been successfully implemented to obtain, for example, proteins of enhanced activities ([18, 19, 20]; Figure S1). However, compared to artificial selection of individuals or mono-species groups, artificial selection of multi-species communities is more challenging (see detailed explanation in Figure S1). For example, member species critical for community function may get lost during selection cycles.

**Figure 1:**
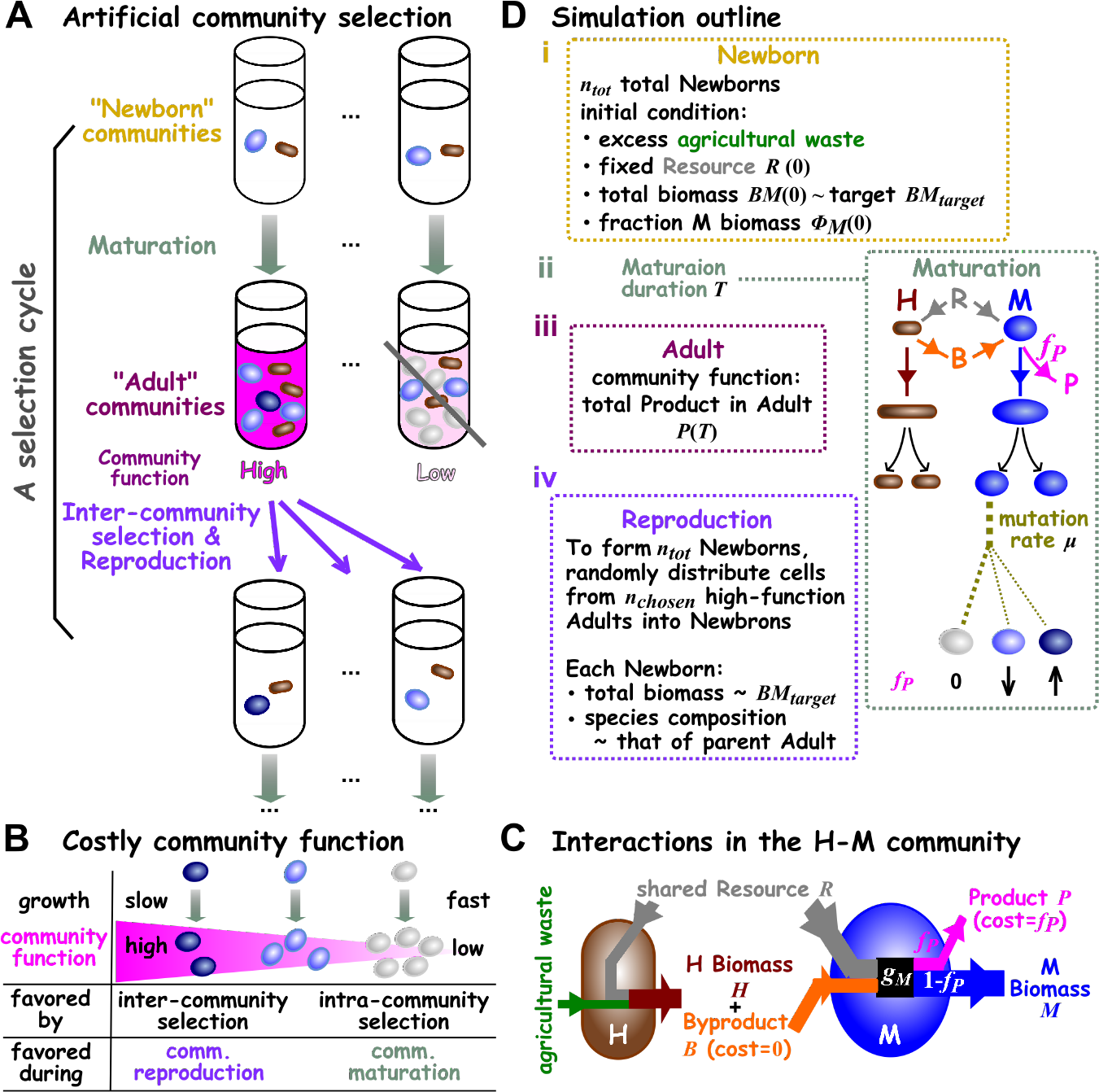
Community selection. (**A**) **Schematic of artificial community selection**. (**B**) **Costly community function**. Darker blue cells contribute more to community function per cell and thus divide more slowly than light cells. High-contributors are disfavored by intra-community selection during community maturation. However, communities dominated by high-contributors are favored by inter-community selection and have a higher chance to reproduce. (**C**) **A Helper-Manufacturer community that converts substrates into a product.** Helper H consumes agricultural waste (present in excess) and Resource to grow biomass, and concomitantly releases Byproduct B at no fitness cost to itself. H’s Byproduct B is required by Manufacturer M. M consumes Resource and H’s Byproduct, and invests a fraction *f*_*P*_ of its potential growth *g*_*M*_ to make Product P while channeling the remaining to biomass growth. When biomass growth ceases, Byproduct and Product are no longer made. The five state variables (italicized) *H*, *M*, *R*, *B*, and *P* correspond to the amount of H biomass, M biomass, Resource, Byproduct, and Product in a community, respectively. (**D**) **Simulating artificial selection of H-M communities**. i. In our simulations, cycles of selection were performed on a total of *n*_*tot*_ = 100 communities with the indicated initial conditions. At the beginning of the first cycle, each Newborn had a total biomass of the target biomass (*BM*_*target*_=100; 60 M and 40 H each of biomass 1). In subsequent cycles, each Newborn’s species ratio would be approximately that of its parent Adult. The amount of Resource in each Newborn was fixed at a value that could support a total biomass of 10^4^ (unless otherwise stated). ii. The maturation time *T* was chosen so that for an average community, Resource was not depleted by time *T* (in experimental terms, this would avoid complications of the stationary phase). During maturation, Resource *R*, Byproduct *B*, Product *P*, and each cell’s biomass were calculated from differential equations (Methods, Section 6). Once a cell’s biomass had grown from 1 to 2, it divided into two identical daughter cells. Death occurred stochastically to individual cells (not depicted). After division, mutations (different shades of oval) occurred stochastically to change a cell’s phenotypes (e.g. M’s *f*_*P*_). iii. At the end of a cycle, community functions (total Product *P*(*T*)) were ranked. iv. During community reproduction, high-functioning Adults were chosen and diluted into Newborns so that on average, each Newborn had a total biomass of approximately the target biomass *BM*_*target*_. A total of *n*_*tot*_ = 100 Newborns were generated for the next selection cycle.

The few attempts at community selection have generated interesting results. One theoretical study simulated artificial selection on multi-species communities based on the presence or absence of a member species [21]. Communities responded to selection, but only under certain conditions. In another theoretical study, multi-species communities responded to artificial selection based on their ability to modify their abiotic environment in user-defined fashions [12]. In both cases, the response to selection quickly leveled off, and could be generated without mutations. Thus, community selection acted entirely on species types instead of new genotypes [21, 12]. In experiments, complex microbial communities were selected for various traits [10, 11, 13, 14]. For example, microbial communities selected to promote early or late flowering in plants were dominated by distinct species types [13]. However in other cases, a community trait may fail to improve despite selection, and may improve even without selection [10, 11].

Because communities used in these selection attempts were complex, much remains unknown. First, was the trait under selection a community function or achievable by a single species? If the latter, then community selection may not be needed, and the simpler task of selecting individuals or mono-species groups could be performed instead (Figure S1). Second, did selection act solely on species types or also on newly-arising genotypes? If selection acted solely on species types ([21, 12, 13]), then without immigration of new species to generate new variations, community function may quickly plateau [21, 12]. If selection acted on genotypes, then community function could continue to improve as new genotypes evolve. Finally, why might a community trait sometimes fail to improve despite selection [10, 11]?

In this study, we simulated artificial selection on communities with two defined species whose phenotypes can be modified by random mutations. Our goal is to improve a “costly” community function. A community function is costly if any community member’s fitness is reduced by contributing to that community function (Figure 1B). Costly community functions are particularly challenging to improve: since contributors to community function grow slower than non-contributors, non-contributors will take over during community maturation. If all Adult communities are dominated by non-contributors, then community selection will fail. To improve a costly community function, inter-community selection during community reproduction (which occurs infrequently once every cycle) must overcome intra-community selection throughout community maturation (Figure 1B).

By simulating a simplified two-species community, we could compare the efficacy of different selection regimens with relative ease, and begin to mechanistically understand evolutionary dynamics during community selection. We also designed our simulations to mimic real lab experiments so that our conclusions could guide future experiments. For example, our simulations incorporated not only chemical mechanisms of species interactions (as advocated by [22, 23]), but also experimental procedures (e.g. pipetting cultures during community reproduction). Model parameters, including species phenotypes, mutation rate, and distribution of mutation effects, were based on a wide variety of published experiments. Note that most previous models focused on binary genotypes (e.g. contributing or not contributing to community function), and therefore could not model community function improvement driven by the evolution of quantitative phenotypes. We show that artificial community selection can improve a costly community function, but only after circumventing a multitude of failure traps.

## Results

We will first introduce the subject of our community selection simulation: a commensal two-species community that converts substrates to a valued product. We will then define community function, and describe how we simulate artificial community selection. Using simulation results, we will demonstrate critical measures that make community selection effective, including promoting species coexistence, suppressing non-contributors, adopting a “bet-hedging” strategy when choosing Adult communities to reproduce, and being mindful about how routine experimental procedures can impede selection. Finally, we show that our conclusions are robust under alternative model assumptions, applicable to mutualistic communities and communities whose member species may not coexist. To avoid confusion, we will use “community selection” or “selection” to describe the entire process of artificial community selection (community formation, growth, selection, and reproduction), and use “choose” or “inter-community selection” to refer to the selection step where the experimentalist decides which communities will reproduce.

### A Helper-Manufacturer community that converts substrates into a product

Motivated by previous successes in engineering two-species microbial communities that convert substrates into useful products [24, 25, 26], we numerically simulated selection of such communities.

In our community, Manufacturer M can manufacture Product P of value to us (e.g. a bio-fuel or a drug) at a fitness cost to self, but only if assisted by Helper H (Figure 1C). Specifically, Helper but not Manufacturer can digest an agricultural waste (e.g. cellulose), and as Helper grows biomass, Helper releases Byproduct B at no fitness cost to itself. Manufacturer requires H’s Byproduct (e.g. carbon source) to grow. In addition, Manufacturer invests *f*_*P*_ (0 ≤ *f*_*P*_ ≤ 1) fraction of its potential growth to make Product P while using the rest (1 − *f*_*P*_) for its biomass growth. Both species also require a shared Resource R (e.g. nitrogen). Thus, the two species together, but not any species alone, can convert substrates (agricultural waste and Resource) into Product.

We define community function as the total amount of Product accumulated as a low-density Newborn community grows into an Adult community over maturation time *T*, i.e. *P*(*T*). In Discussions, we explain problems associated with an alternative definition of community function (e.g. per capita production; Methods Section 7; Figure S2). We will initially focus on the scenario where community function is not costly to Helpers, but incurs a fitness cost of *f*_*P*_ to M. Later, we will show that our conclusions also hold when community function is costly to both H and M. Below, we will describe how we simulated community selection, followed by how we chose parameters of species phenotypes and parameters of selection regimen.

### Simulating community selection

We simulated four stages of community selection (Figure 1D): (i) forming Newborn communities; (ii) Newborn communities maturing into Adult communities; (iii) choosing high-functioning Adult communities, and (iv) reproducing the chosen Adult communities by splitting each into multiple Newborn communities of the next cycle. Our simulation was individual-based. That is, it tracked phenotypes and biomass of individual H and M cells in each community as cells grew, divided, mutated, or died. Our simulations also tracked dynamics of chemicals (including Product) in each community, and accounted for actual experimental steps such as pipetting cultures during community reproduction. Below is a brief summary of our simulations, with more details in Methods (Section 6).

Each simulation started with *n*_*tot*_ number of Newborn communities. Each Newborn community always started with a fixed amount of Resource *R*(0) and a total biomass close to a target value *BM*_*target*_ (see Discussions for problems associated with not having a biomass target, such as diluting an Adult by a fixed fold into Newborns). Agricultural waste was always supplied in excess and thus did not enter our equations. Note that except for the first cycle, the relative abundance of species in a Newborn community was approximately that of its parent Adult community.

During community maturation, biomass of individual cells grew. The biomass growth rate of an H cell depended on Resource concentration (Monod equation; Figure S3A; Eq. 23). As H grew, it consumed Resource and simultaneously released Byproduct (Eqs. 21 and 22). The potential growth rate of an M cell depended on the concentrations of Resource and H’s Byproduct (Mankad-Bungay dual-nutrient equation [27]; Figure S3B; see experimental support in Figure S4). M cell’s actual biomass growth rate was (1 − *f*_*P*_) fraction of M’s potential growth rate (Eq. 24). As M grew, it consumed Resource and Byproduct (Eqs. 21 and 22), and released Product at a rate proportional to *f*_*P*_ and M’s potential growth rate (Eqs. 8). Once an H or M cell’s biomass grew from 1 to 2, it divided into two cells of equal biomass with identical phenotypes, thus capturing experimental observations of continuous biomass increase (Figure S5) and discrete cell division events [28]. Meanwhile, H and M cells died stochastically at a constant death rate. Although mutations can occur during any stage of the cell cycle, we assigned mutations immediately after cell division, where each phenotype of both cells mutated independently.

Mutable phenotypes included H and M’s maximal growth rates and affinities for nutrients (“growth parameters”), and M’s *f*_*P*_ (the fraction of potential growth diverted for making Product), since these phenotypes have been observed to rapidly change during evolution ([29, 30, 31, 32]). Mutated phenotypes could range between 0 and their respective evolutionary upper bounds. Among mutations that alter phenotypes (denoted “mutations”), on average, half abolished the function (e.g. zero growth rate, zero affinity, or *f*_*P*_ = 0) based on experiments on GFP, viruses, and yeast [33, 34, 35]. Effects of the other 50% mutations were bilateral-exponentially distributed, enhancing or diminishing a phenotype by a few percent, based on our re-analysis of published yeast data sets [36] (Figure S6). We held death rates constant, since death rates were much smaller than growth rates and thus mutations in death rates would be inconsequential. We also held release and consumption coefficients constant. This is because, for example, the amount of Byproduct released per H biomass generated is constrained by biochemical stoichiometry.

At the end of community maturation time *T*, we compared community function *P*(*T*) (the total amount of Product accumulated in the community by time *T*) for each Adult community, and chose high-functioning Adults to reproduce. Each chosen Adult was randomly partitioned into Newborns with target total biomass *BM*_*target*_. For example, if the chosen Adult had a total biomass of 60*BM*_*target*_, then each cell would be assigned a random integer from 1 to 60, and those cells with the same random integer would be allocated to the same Newborn. Experimentally, this is equivalent to volumetric dilution using a pipette. Thus, for each Newborn, the total biomass and species ratio fluctuated around their expected values in a fashion associated with pipetting (Methods Section 9). In the “top-dog” strategy, we always chose the highest-functioning Adult available to us: after the highest-functioning Adult was used up for making Newborns, we then reproduced the next highest-functioning Adult in the same way and randomly chose enough Newborns so that a total of *n*_*tot*_ Newborns were generated for the next selection cycle.

### Choosing species: enhancing species coexistence

In order to improve community function through community selection, species need to coexist throughout selection cycles. That is, all species must grow at a similar average growth rate within each cycle. Furthermore, species ratio should not be extreme because otherwise, the low-abundance species could be lost by chance during Newborn formation. Species coexistence at a moderate ratio has been experimentally realized in engineered communities [24, 25, 37, 38].

To achieve species coexistence at a moderate ratio in the H-M community, three considerations need to be made. First, the fraction of growth M diverted for making Product (*f*_*P*_) must not be too large, otherwise M would always grow slower than H and thus eventually go extinct (Figure 2A, top). Second, H and M’s growth parameters (maximal growth rates in excess nutrients; affinities for nutrients) must be balanced. This is because upon Newborn formation, H can immediately start to grow on agricultural waste and Resource, while M cannot grow until H’s Byproduct has accumulated to a sufficiently high level. Thus to achieve coexistence, M must grow faster than H at some point during community maturation. Third, to achieve a moderate steady-state species ratio, metabolite release and consumption need to be balanced [37]. Otherwise, the ratio between metabolite releaser and consumer can be extreme.

**Figure 2:**
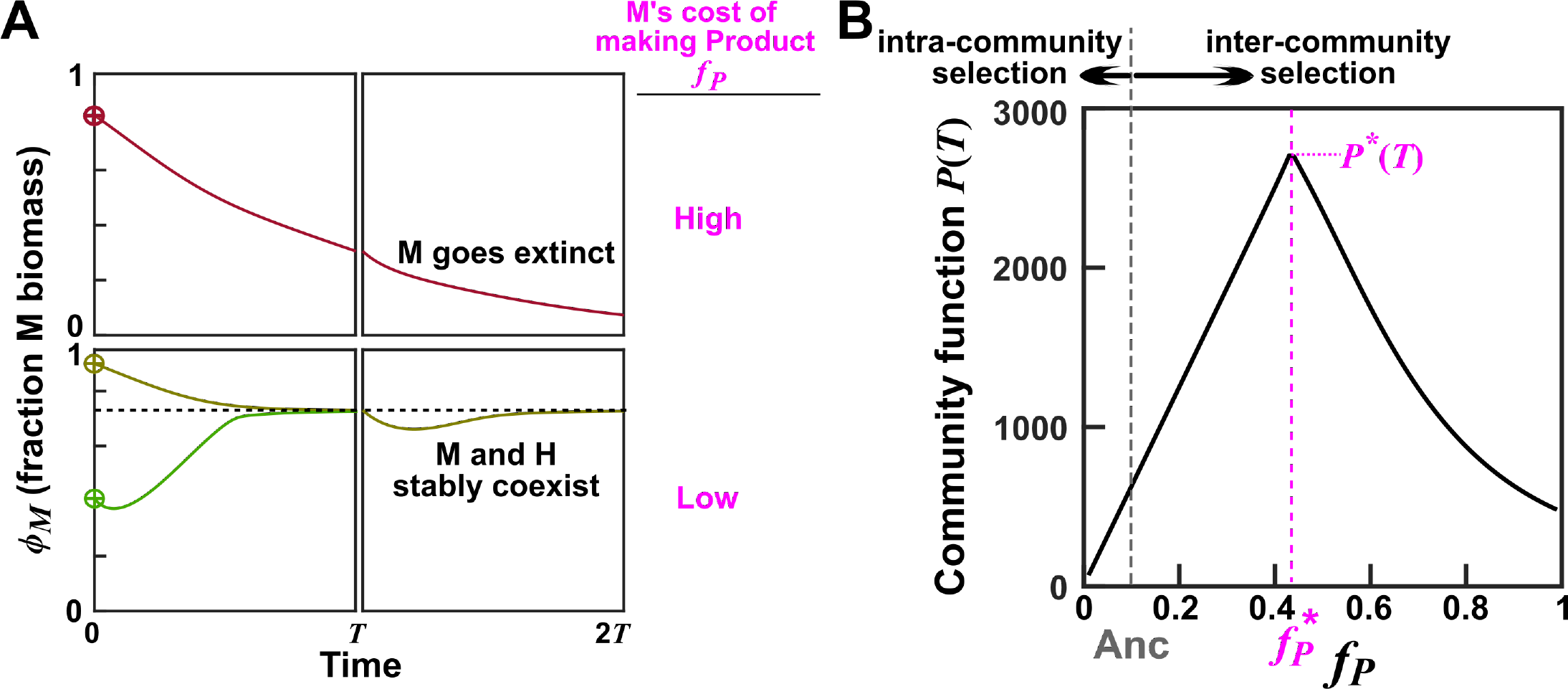
Optimal community function is achieved at an intermediate cost. Calculations were based on equations 6-10 with H and M’s growth parameters fixed at their respective evolutionary upper bounds (Table 1, last column). (**A**) **H and M can stably coexist at low** *f*_*P*_. **Top**: When *f*_*P*_, the fraction of potential growth Manufacturer diverts for making Product, is high (e.g. *f*_*P*_ = 0.8), M will eventually go extinct (i.e. fraction of M < 1/total population). **Bottom**: At low *f*_*P*_ (e.g. *f*_*P*_ = 0.1), H and M can stably coexist. That is, different initial species ratios will converge to a steady state value. At the end of the first cycle (time *T* = 17), Byproduct and Resource were re-set to the initial conditions at time zero (0 and 10^4^, respectively), and total biomass was reduced to the target value *BM*_*target*_ (=100) while the fraction of M biomass *ϕ*_*M*_ remained the same as that of the parent community. See main text for how values of maturation time and Resource were chosen. (**B**) **Optimal community function occurs at an intermediate** *f*_*P*_. Community functions at various combinations of *f*_*P*_ and fraction of M biomass (out of *BM*_*target*_ = 100 total biomass) were computed by integrating Eqs 6-10. Maximal community function *P*(*T*) is achieved at an intermediate 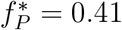 (magenta dashed line) when Newborn species composition is also optimal (46 H and 54 M cells). Note that at zero *f*_*P*_, no Product would be made; at *f*_*P*_ = 1, M would go extinct. The maximal *P*\*(*T*) could not be further improved even if we allowed all growth parameters and *f*_*P*_ to mutate (Figure S10). Thus, *P*\*(*T*) is locally maximal in the sense that small deviation will always reduce *P*(*T*). Ancestral *f*_*P*_ (grey) is lower than 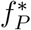. The central question is: can community selection improve *f*_*P*_ to 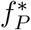 despite natural selection’s favoring lower *f*_*P*_?

Based on these considerations and published yeast and *E. coli* measurements, we chose H and M’s ancestral growth parameters and their evolutionary upper bounds, as well as release, consumption, and death parameters (Table 1, Methods Section 2). This ensured that throughout evolution, different species ratios would converge toward a moderate steady state value during community maturation (Figure 2A, bottom). Note that if species were not chosen properly, selection might fail due to insufficient species coexistence (Figure 6A), although we will demonstrate that under effective community selection, requirements on species coexistence could be relaxed (Figure 6B and C).

**Table 1:** Parameters for ancestral and evolved (growth- and mono-adapted) H and M. Parameters in the “Evolved” column are used for most simulations and figures unless otherwise specified. For maximal growth rates, * represents evolutionary upper bound. For *K*_*SpeciesM etabolite*_, * represents evolutionary lower bound, which corresponds to evolutionary upper bound for Species’s affinity for Metabolite (1/*K*_*SpeciesMetabolite*_). # is from Figure S25B. In Methods Section 2, we explained our parameter choices (including why we hold some parameters constant during evolution).

|  | Definition | Ancestral | Evolved |
| --- | --- | --- | --- |
| $\tilde{r}_B$ | amount of $\hat{B}$ released per H biomass grown | scaling factor, 1 | no change |
| $\tilde{r}_P$ | amount of $\hat{P}$ released at the cost of one M biomass | scaling factor, 1 | no change |
| $\tilde{R}(0)$ | initial amount of Resource in Newborn | scaling factor, 1 | |
| $f_P$ | fraction of M growth diverted to producing P | 0.10 | 0.13# |
| $K_{MR}$ | fold of $\tilde{R}(0)$ at which $g_{Mmax}/2$ is achieved in excess B | 1 | 1/3* |
| $K_{MB}$ | amount of $\hat{B}$ at which $g_{Mmax}/2$ is achieved in excess R, scaled against $\tilde{r}_B$ | $\frac{5}{3} \times 10^2$ | $\frac{1}{3} \times 10^2*$ |
| $K_{HR}$ | fold of $\tilde{R}(0)$ at which $g_{Hmax}/2$ is achieved | 1 | 1/5* |
| $g_{Mmax}$ | maximal biomass growth rate of M | 0.58/unit time | 0.7/unit time* |
| $g_{Hmax}$ | maximal biomass growth rate of H | 0.25/unit time | 0.3/unit time* |
| $\delta_M$ | death rate of M | $3.5 \times 10^{-3}$/unit time | no change |
| $\delta_H$ | death rate of H | $1.5 \times 10^{-3}$/unit time | no change |
| $c_{RM}$ | fraction of $\tilde{R}(0)$ consumed per M biomass grown | $10^{-4}$ | no change |
| $c_{RH}$ | fraction of $\tilde{R}(0)$ consumed per H biomass grown | $10^{-4}$ | no change |
| $c_{BM}$ | amount of $\hat{B}$ consumed per M biomass grown, scaled against $\tilde{r}_B$ | $\frac{1}{3}$ | no change |
| $P_{mut}$ | mutation probability per cell division for each mutable phenotype | $2 \times 10^{-5} \sim 2 \times 10^{-3}$ | |

### Choosing selection regimen parameters: avoiding known failure modes

After choosing member species with appropriate phenotypes, we need to consider the parameters of our selection regimen (Figure 1D). These parameters include the total number of communities under selection (*n*_*tot*_), the number of Adult communities chosen to reproduce (*n*_*chosen*_), Newborn target total biomass (*BM*_*target*_) which indicates the “bottleneck size” when splitting an Adult community into Newborn communities, the amount of Resource added to each Newborn (*R*(0)), the amount of mutagenesis which controls the rate of phenotype-altering mutations (*μ*), and maturation time (*T*). Compared to the well-studied problem of group selection where the unit of selection is a mono-species group [39, 40, 41, 42, 43, 44, 45, 46, 47, 48, 49, 50, 51, 52, 53], community selection is more challenging (Discussions; Figure S1). However, the two types of selections do share some common aspects (Discussions; Figure S1). Thus, we can apply group selection theory, together with other practical considerations, to better design community selection regimen.

If the total number of communities *n*_*tot*_ is very large, then the chosen community will likely display a higher community function than if *n*_*tot*_ is small, but experimentally achieving a large *n*_*tot*_ is more challenging. We chose a total of 100 communities (*n*_*tot*_=100).

*n*_*chosen*_, the number of Adults chosen by the experimentalist to reproduce, reflects selection strength. Since the top-functioning Adult is presumably the most desirable, we reproduced it into as many Newborns as possible, and then reproduced the second best etc until we obtained *n*_*tot*_ Newborn communities for the next cycle (the “top-dog” strategy). Later, we will compare the “top-dog” strategy with other strategies employing weaker selection strengths.

If the mutation rate is very low, then community function cannot rapidly improve. If the mutation rate is very high, then non-contributors will be generated at a high rate, and as the fast-growing non-contributors take over during community maturation, community function will likely collapse. Here, we chose *μ*, the rate of phenotype-altering mutations, to be biologically realistic (0.002 per cell per generation per phenotype, which is lower than the highest values observed experimentally; Methods Section 4).

If Newborn target total biomass *BM*_*target*_ is very large, or if the number of doublings within maturation time *T* is very large, then non-contributors will take over in all communities during maturation (Figure S7, compare B-D with A), as predicted by group selection theory. On the other hand, if both *BM*_*target*_ and the number of generations within *T* are very small, then mutations will be rare within each cycle, and many cycles will be required to improve community function. Finally, if *BM*_*target*_ is very small, then a member species might get lost by chance during Newborn formation. In our simulations, we chose Newborn’s target total biomass *BM*_*target*_ = 100 biomass (50~100 cells). Unless otherwise stated, we fixed the input Resource *R*(0) to support a maximal total biomass of 10^4^, and chose maturation time *T* so that even if H and M had evolved to grow as fast as possible, total biomass would undergo ~6 doublings (increasing from ~100 to ~7000). Thus, by the end of *T*, ≤70% Resource would be consumed by an average community. This meant that when implemented experimentally, we could avoid complications of Resource depletion and stationary phase, while not wasting too much Resource.

### Community selection may not be effective under conditions reflecting common lab practices

We initially simulated community selection, only allowing M’s *f*_*P*_ to be modified by mutations while fixing H and M’s growth parameters (maximal growth rates in excess metabolites; affinities for metabolites) to their evolutionary upper bounds. Such a simplification is justified with our particular parameter choices (Table 1) for the following reasons. First, during community selection growth parameters improved to their evolutionary upper bounds anyways (Figure S8C and F). Second, we obtained qualitatively similar conclusions regardless of whether we fixed growth parameters or not (e.g. compare final community functions in Figure 3B and E versus Figure S8A and D). Later, we will show a case where growth parameters cannot be fixed to upper bounds (Figure 6). In the current case if Newborn’s total biomass is fixed to the target value, then with only *f*_*P*_ mutating, we can calculate the theoretical maximal community function *P*\*(*T*) and its associated optimal 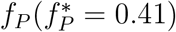 and optimal species ratio (Figure 2B). We started with ancestral *f*_*P*_ lower than 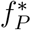. Could inter-community selection for high community function increase ancestral *f*_*P*_ to 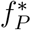, despite intra-community natural selection favoring lower *f*_*P*_?

**Figure 3:**
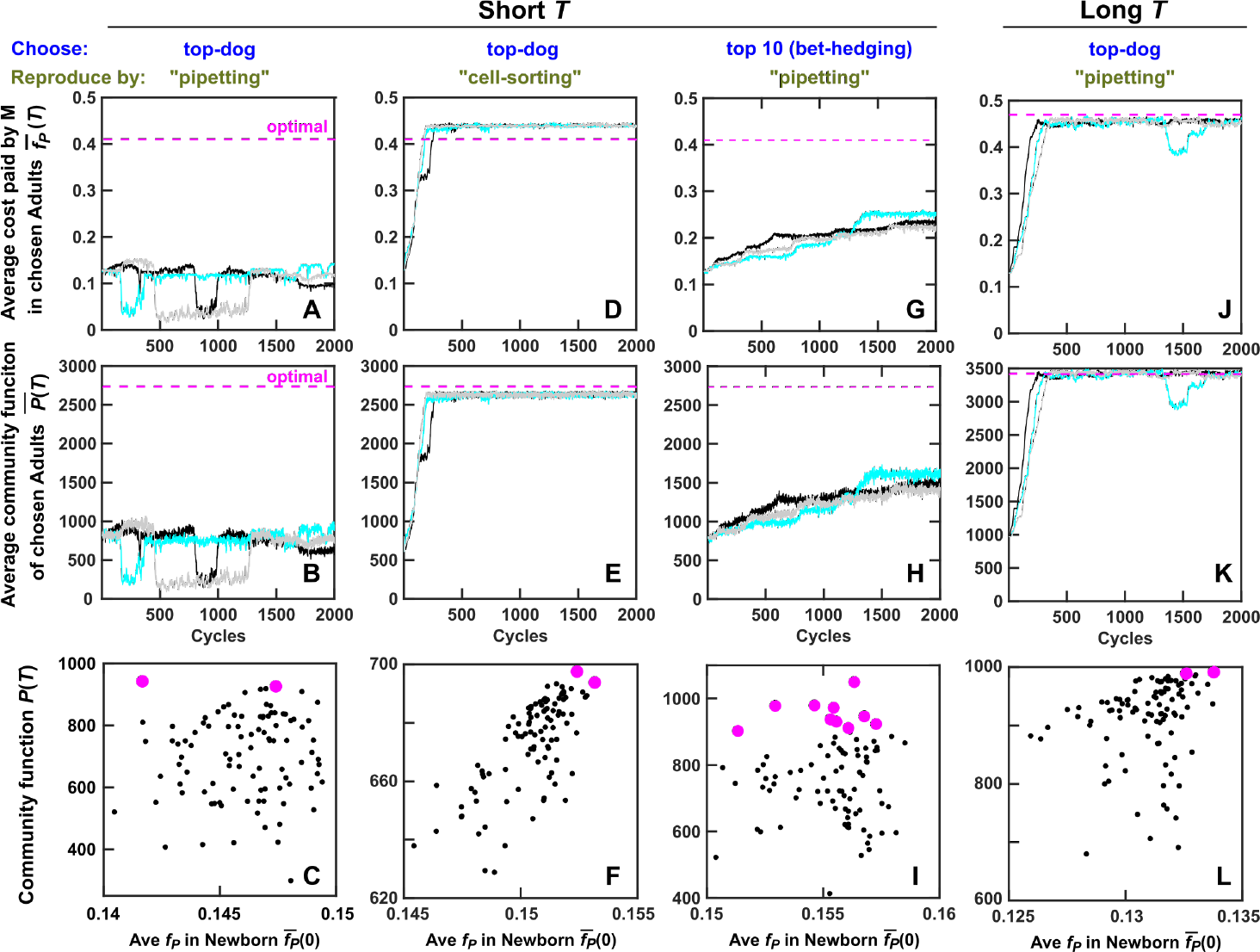
Community selection can be stalled by routine experimental procedures, and succeeds when community function correlates with its heritable determinant or under the “bet-hedging” strategy. (**A-I**) Evolution dynamics when the maturation time *T* was sufficiently short to avoid Resource depletion and stationary phase (*T* = 17). (**A-C**) Adults were chosen using the “top-dog” strategy, and diluted into progeny Newborns through pipetting (i.e. H and M biomass fluctuated around their expected values). Community selection was not effective. Average *f*_*P*_ and community function failed to improve to their theoretical optima, and community function poorly correlated with its heritable determinant 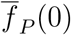. Black and magenta dots: un-chosen and chosen communities from one selection cycle, respectively. (**D-F**) Adults were chosen using the “top-dog” strategy, and a fixed H biomass and M biomass from the chosen Adults were sorted into Newborns. Community selection was successful. Community function also correlated with its heritable determinant 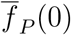. Here, Newborn total biomass *BM*(0) and fraction of M biomass *ϕ*_*M*_(0) were respectively fixed to *BM*_*target*_ = 100 and *ϕ*_*M*_(*T*) of the chosen Adult of the previous cycle. (**G-I**) When we chose the top ten Adults and let each reproduce 10 Newborns via pipetting, community function improved despite poor correlation between community function and its heritable determinant 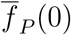. For selection dynamics over many cycles, see Figure S14. (**J-L**) Evolution dynamics when maturation time was long (*T* = 20) such that most Resource was consumed by the end of *T*. Adults were chosen using the “top-dog” strategy, and reproduced via “pipetting”. Community selection was successful due to high correlation between community function and its heritable determinant 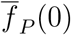, assuming that variable duration in stationary phase would not introduce non-heritable variations in community function. Black, cyan and gray curves are three independent simulation trials. 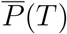 was the average of *P*(*T*) across all chosen Adults. 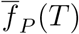 was obtained by first averaging among M within each chosen Adult and then averaging across all chosen Adults.

As expected, in control simulations where Adult communities were randomly chosen to reproduce, community function was driven to zero by natural selection as fast-growing non-producing M took over (Figure S9).

When we used the “top-dog” strategy and chose the top-functioning communities to reproduce, *f*_*P*_ and community function *P*(*T*) did not decline to zero, but they barely improved, and both were far below their theoretical optima (Figure 3A and B).

### Common lab practices can generate sufficiently large non-heritable variations in community function to interfere with selection

Why did community selection fail to improve *f*_*P*_ and community function? One possibility is that community function was not sufficiently heritable from one cycle to the next (Figure S1). We therefore investigated the heredity of community function by examining the heredity of community function determinants.

Community function *P*(*T*) was largely determined by phenotypes of cells in the Newborn community. This is because maturation time was sufficiently short (~6 doublings) that newly-arising genotypes could not rise to high frequencies within one cycle to significantly affect community function. Since all phenotypes except for *f*_*P*_ were fixed, community function had three independent determinants: Newborn’s total biomass *BM*(0), Newborn’s fraction of M biomass *ϕ*_*M*_(0), and the average *f*_*P*_ over all M cells in Newborn 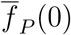 (Eq 6-10).

A community function determinant is considered heritable if it is correlated between Newborns of one cycle (Figure 4A, bottom row) and their respective progeny Newborns in the next cycle (Figure 4A, color-matched top row). Among the three determinants, 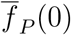 was heritable (Figure 4B): if a Newborn community had a high average *f*_*P*_, so would the mature Adult community and Newborn communities reproduced from it. On the other hand, Newborn total biomass *BM*(0) was not heritable (Figure 4C). This is because when an Adult community reproduced via pipette dilution, the dilution factor was adjusted so that the total biomass of a progeny Newborn community was on average the target biomass *BM*_*target*_. Newborn’s fraction of M biomass *ϕ*_*M*_(0), which fluctuated around that of its parent Adult, was not heritable either (Figure 4D). This is because regardless of the species composition of Newborns, Adults would have similar steady state species composition (Figure 2A bottom panel), and so would their offspring Newborns.

**Figure 4:**
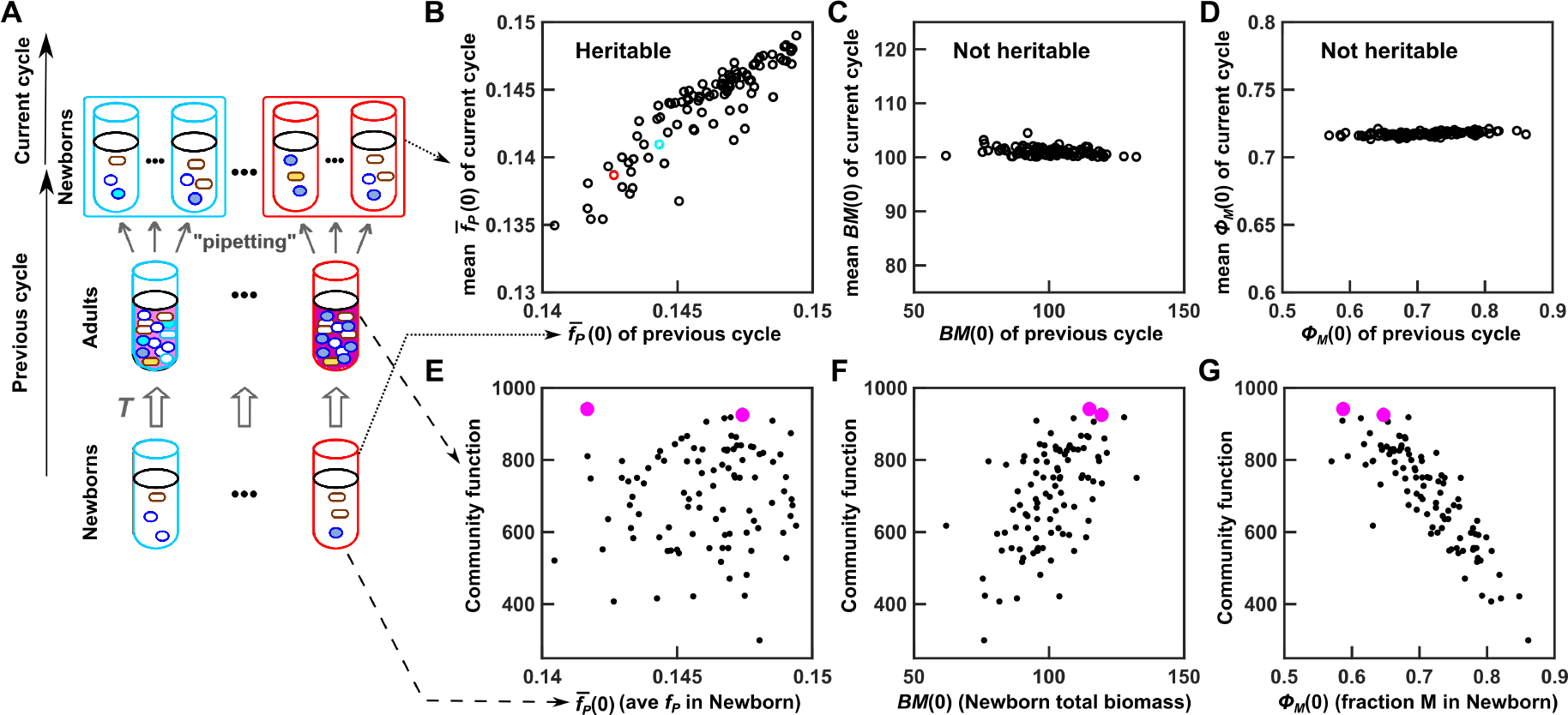
During ineffective community selection, community function correlates weakly with its heritable determinant and strongly with non-heritable determinants. (**A**) Schematic. Newborns and corresponding Adults from the “Previous cycle” were taken from the 180th cycle of the simulation displayed in black of Figure 3A-B. We then allowed each Adult to reproduce Newborns (“current cycle”), forming 100 lineages (tubes with the same color outline belong to the same lineage). (**B**-**D**) Among the three determinants of community function, 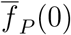 (*f*_*P*_ averaged among M cells in Newborn) is heritable, but *BM*(0) (total biomass of Newborn) and *ϕ*_*M*_(0) (fraction of M biomass in Newborn) are not. For each lineage, the community function determinant at the previous cycle was scatter plotted against the average value at the current cycle. (**E**-**G**) During ineffective community selection (Figure 3B), *P*(*T*) correlates weakly with heritable determinant, but strongly with non-heritable determinants. Each dot represents one community, and the two magenta dots represent the two “successful” Newborns that achieved the highest community function at adulthood.

In successful community selection, variations in community function should be mainly caused by variations in its heritable determinants. However, we found that community function *P*(*T*) weakly correlated with its heritable determinant 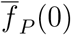, but strongly correlated with its non-heritable determinants (Figure 4E-G). For example, the Newborn that would achieve the highest function had a below-median 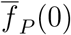 (left magenta dot in Figure 4E), but had high total biomass *BM*(0) and low fraction of M biomass *ϕ*_*M*_(0) (Figure 4F, G). In other words, variation in community function is largely non-heritable, as it largely arises from variation in non-heritable determinants.

The reason for strong correlations between *P*(*T*) and the two non-heritable determinants became clear by examining community dynamics. Recall that to avoid stationary phase, we had chosen maturation time so that Resource would be in excess by the end of maturation. Thus, a “lucky” Newborn community starting with a higher-than-average total biomass would convert more Resource to Product (dotted lines in top panels of Figure S11). Similarly, if a Newborn started with higher-than-average fraction of Helper H biomass, then H would produce higher-than-average Byproduct which meant that M would endure a shorter growth lag and make more Product (dotted lines in bottom panels of Figure S11).

To summarize, when community function significantly correlated with its non-heritable determinants (Figure 4F & G), community selection failed to improve community function (Figure 3B).

### Reducing non-heritable variations in an experimentally feasible manner promotes artificial community selection

Reducing non-heritable variations in community function should enable community selection to work. One possibility would be to reduce the stochastic fluctuations in non-heritable determinants. Indeed, when each Newborn received a fixed biomass of Helper H and Manufacturer M (“cell-sorting”; i.e. fixing *BM*(0) and *ϕ*_*M*_(0); Methods, Section 6), community function *P*(*T*) became strongly correlated with its heritable determinant 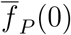 (Figure 3F). In this case, both 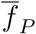 and community function *P*(*T*) improved under selection (Figure 3, D and E) to near the optimal. *P*(*T*) improvement was not seen if either Newborn total biomass or species fraction was allowed to fluctuate stochastically (Figure S12). *P*(*T*) also improved if fixed numbers of H and M cells (instead of biomass) were allocated into each Newborn, even though each cell’s biomass fluctuated between 1 and 2 (Figure S13C; Methods, Section 6). Allocating a fixed biomass or number of cells from each species to Newborn communities could be experimentally realized using a cell sorter if different species have different fluorescence ([54]).

Non-heritable variations in *P*(*T*) could also be curtailed by reducing the dependence of *P*(*T*) on non-heritable determinants. For example, we could extend the maturation time *T* to nearly deplete Resource. In this selection regimen, Newborns would still experience stochastic fluctuations in Newborn total biomass *BM*(0) and fraction of M biomass *ϕ*_*M*_(0). However, all communities would end up with similar *P*(*T*) since “unlucky” communities would have time to “catch up” as “lucky” communities wait in stationary phase. Indeed, with this extended *T*, community function became strongly correlated with its heritable determinant 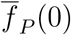 and community function improved without having to fix Newborn total biomass or species composition (Figure 3, J-L; Figure S13D). However in practice, non-heritable variations in community function could still arise from stochastic fluctuations in the duration of stationary phase (which could affect cell survival or recovery time in the next selection cycle).

### Bet-hedging can promote community selection when non-heritable variations in community function hinder selection

Since the highest community function may not correspond to the highest *f*_*P*_ (Figure 3C), we examined whether a “bet-hedging” strategy might outperform the “top-dog” strategy. In the “bet-hedging” strategy, we chose, for example, the top ten Adults, each reproducing ten Newborns. Although the “bet-hedging” strategy did not work as effectively as minimizing non-heritable variations in community function (compare Figure 3 D-F “cell-sorting” with G-I “bet-hedging”; Figure S14), “bet-hedging” under a wide range of selection strengths outperformed “top-dog”. Specifically, when we used pipetting to dilute Adults into Newborns, community function failed to improve under the “top-dog” strategy, but improved when top 5 to top 50 Adult communities were chosen to reproduce (Figure S15; Figure 3, compare G-I with A-C).

The superiority of “bet-hedging” over “top-dog” rests on giving “unlucky” Adults a chance to reproduce. We reached this conclusion by noting that if we minimized non-heritable variations in community function by fixing species biomass in Newborns (“cell-sorting”), then the “top-dog” strategy is superior to the “bet-hedging” strategy (Figure S16).

As expected, uncertainty in community function measurement - another source of non-heritable variation - interferes with community selection (compare Figure 3 A-I with Figure 5 A, C). In this case, “bet-hedging” and “cell-sorting” can synergize to increase the efficacy of community selection (Figure 5, compare B and C with D). Since it is difficult to suppress non-heritable variations in community function, the “bet-hedging” strategy could be useful for experimentalists.

**Figure 5:**
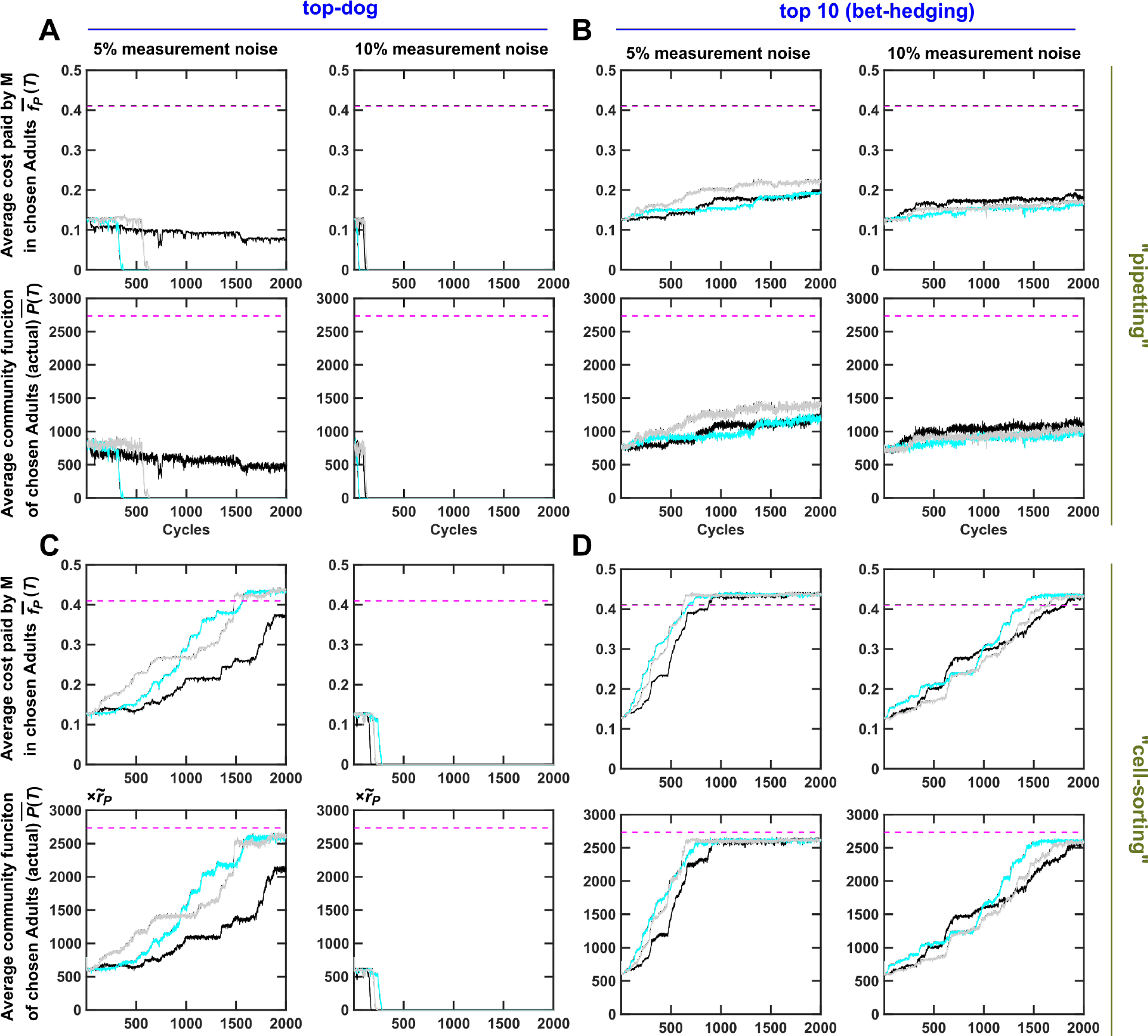
Community selection impeded by community function measurement noise can be rescued by “bet-hedging” acting in synergy with “cell-sorting”. Adult communities were chosen to reproduce based on “measured community function *P*(*T*)” - the sum of actual *P*(*T*) and an “uncertainty term” randomly drawn from a normal distribution with zero mean and standard deviations of 5% or 10% of the ancestral *P*(*T*). Dynamics of average *f*_*P*_ and average community function of the chosen Adult communities (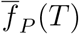 and 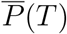) are plotted. When uncertainty in community function measurement is low (5%), cell-sorting largely rescues ineffective community selection (**A-D**, left panels). When uncertainty in community function measurement is high (10%), both cell-sorting and bet-hedging are required (**A-D**, right panels). Black, cyan and gray curves are three independent simulation trials. 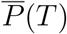 was averaged across the chosen Adults. 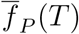 was obtained by first averaging among M within each chosen Adult and then averaging across the chosen Adults.

**Figure 6:**
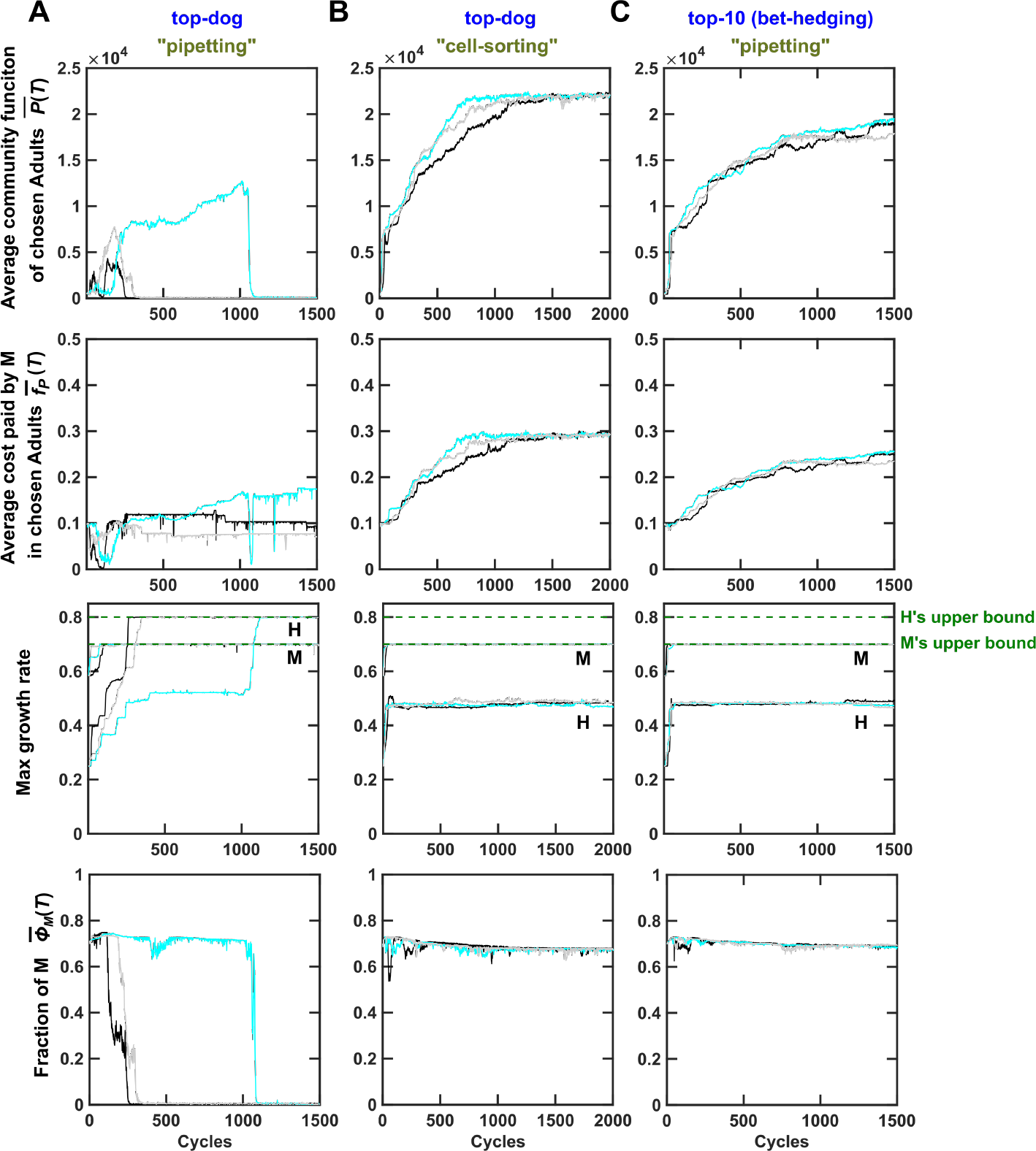
Effective community selection can encourage species coexistence. Here, the evolutionary upper bound for 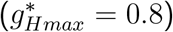 was larger than that for 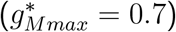, opposite to that in Figures 2-5 (**A**) When the “top-dog” strategy and “pipetting” were used to choose and reproduce Adult communities, M was almost outcompeted by H as H evolved to grow faster than M (third panel). Although M would ordinarily go extinct, community selection managed to maintain M at a very low level (bottom). This imbalanced species ratio resulted in very low community function (top). When community selection was effective, either using the “top-dog” strategy with “cell-sorting” (**B**), or the “bet-hedging” strategy with “pipetting” (**C**), community selection successfully improved community function and 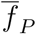. Strikingly in both **B** and **C**, H’s growth parameter did not increase to its evolutionary upper bound (Figure S17B and C), allowing a balanced species ratio (bottom) and high community function (top). Resource supplied to Newborn communities here supports 10^5^ total biomass to accommodate faster growth rates (and hence community function is larger than in other figures). Black, cyan and gray curves are three independent simulation trials. 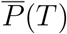 and 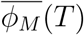 (fraction of M biomass in Adult communities) were averaged across the chosen Adults. 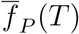 was obtained by first averaging among M within each chosen Adult and then averaging across all chosen Adults.

### Community selection can enforce species coexistence

In most communities, species coexistence may not be guaranteed due to competition for shared resources. Here, we show that properly executed community selection could also improve the functions of such communities, in part by enforcing species coexistence. Consider an H-M community where, unlike the H-M community we have considered so far, H had the evolutionary potential to grow much faster than M. In this case, high community function not only required M to pay a fitness cost of *f*_*P*_, but also required H to grow sufficiently slowly to not out-compete M.

We started community selection at ancestral growth parameters, and allowed them and *f*_*P*_ to mutate. When community selection was ineffective (“top-dog” with “pipetting”; Figure 6A), H’s maximal growth rate evolved to exceed M’s maximal growth rate (Figure 6A, Figure S17). This drove M to almost extinction, and community function was very low (Figure 6A). During effective community selection (“top-dog” with “cell-sorting” or “bet-hedging” with “pipetting”, Figure 6B-C), H’s maximal growth rate remained far below its evolutionary upper bound and far below M’s maximal growth rate (Figure 6B-C). In this case, H and M can coexist at a moderate ratio, and community function improved (Figure 6B-C).

### Robust conclusions under alternative model assumptions

We have demonstrated that when selecting for high H-M community function, seemingly innocuous experimental procedures (e.g. choosing the top-functioning Adults and pipetting portions of them to form Newborns) could be problematic. Instead, more precise procedures (e.g. “cell-sorting”) or reduced selection strength (e.g. “bed-hedging”) might be required. Our conclusions hold when we used a much lower mutation rate (2×10^−5^ instead of 2×10^−3^ mutation per cell per generation per phenotype, Figure S18), although lower mutation rate slowed down community function improvement. Our conclusions also hold when we used a different distribution of mutation effects (a non-null mutation increasing or decreasing *f*_*P*_ by on average 2% or eliminating null mutants; Figure S19), or incorporating epistasis (i.e. a non-null mutation would likely reduce *f*_*P*_ if the current *f*_*P*_ was high, and enhance *f*_*P*_ if the current *f*_*P*_ was low; Figure S20; Figure S21; Methods Section 5).

To further test the generality of our conclusions, we simulated community selection on a mutualistic H-M community. Specifically, we assumed that Byproduct was inhibitory to H. Thus, H benefited M by providing Byproduct, and M benefited H by removing the inhibitory Byproduct, similar to the syntrophic community of *Desulfovibrio vulgaris* and *Methanococcus maripaludis* [55]. We obtained similar conclusions in this mutualistic H-M community (Figure S22). We have also shown that similar conclusions hold for communities where member species may not coexist (Figure 6).

In summary, our conclusions seem general under a variety of model assumptions, and apply to a variety of communities.

## Discussions

How might we improve functions of multi-species microbial communities via artificial selection? A common and highly valuable approach is to identify appropriate combinations of species types. For example, by using cellulose as the main carbon source in a process called “enrichment”, Kato et al. obtained a community consisting of a few species that together degrade cellulose [56]. A more elaborate scheme is to perform artificial community selection to improve a desired community trait [10, 11, 14, 13, 17]. However, if we solely rely on species types, then without a constant influx of new species, community function will likely level off quickly [12, 21]. Here, we consider artificial selection of communities with defined member species so that improvement of community function requires new genotypes that contribute more toward the community function of interest at a cost to itself.

### Community selection can be challenging but is feasible

Artificial selection of whole communities to improve a costly community function requires careful considerations. We have considered species choice (Figures 2), mutation rate, the total number of communities under selection, Newborn target total biomass (bottleneck size; Figure S7), the number of generations during maturation (which in turn depends on the amount of Resource added to each Newborn and the maturation time; Figure S7), selection strength (Figure 3), and how we reproduce Adults (e.g. volumetric pipetting versus “cell-sorting”, Figure 3), and the uncertainty in community function measurements (Figure 5).

Many of these considerations face dilemmas. For example, a large Newborn size (*BM*_*target*_) would lead to reproducible take-over by non-contributors (Figure S7), but a small Newborn size would mean that large non-heritable variations in community function can readily arise and interfere with selection unless special measures are taken (Figure 3).

We can take obvious steps to mitigate non-heritable variations in community function. For example, we can repeatedly measure community function to increase measurement precision, thereby facilitating selection (Figure 5). We can also use the bet-hedging strategy so that lower-functioning communities harboring desirable genotypes can have a chance to reproduce (Figure 3). We can use a cell sorter to fix the cell number or biomass of member species in Newborns so that community function suffers less non-heritable variations (Figure 3).

The need to suppress non-heritable variations in community function can have practical implications that may initially seem non-intuitive. For example, when shared resource is non-limiting (to avoid stationary phase), we must dilute a chosen Adult community to a fixed target biomass instead of by a fixed-fold. This is because otherwise, selection would fail as we choose larger and larger Newborn size instead of higher and higher *f*_*P*_ (Methods, Section 7; Figure S23).

The definition of community function is also critical. If we had defined community function as Product per M biomass in the Adult community *P*(*T*)/*M*(*T*) (which is approximately proportional to 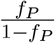: see Methods Section 7), then we will be selecting for higher and higher *f*_*P*_, and M can go extinct (Figure S2). Note that certain types of community function might be easier to improve if they are insensitive to fluctuations in Newborn species biomass. An example is the H:M ratio which converges to a fixed value during maturation (Figure 2A bottom panel).

### Intra-community selection versus inter-community selection

When improving a costly community function, intra-community selection and inter-community selection are both important. Intra-community selection occurs during community maturation, and favors fast growers. Inter-community selection occurs during community reproduction, and favors high community function.

For production cost *f*_*P*_, intra-community selection favors low *f*_*P*_, while inter-community selection favors 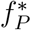, the *f*_*P*_ leading to the highest community function. Thus, when current 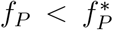, inter-community selection runs against intra-community selection. When current 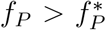, intra- and inter-community selections are aligned.

For growth parameters (maximal growth rates, affinities for metabolites), depending on their evolutionary upper bounds, the two selection forces may or may not be aligned. For example, using parameters in Table 1, improving growth parameters promoted community function (Figure S8A-C; Figure S24A). This is because with these choices of evolutionary upper bounds, H could not evolve to grow so fast to overwhelm M, Thus, with sufficient Resource and without the danger of species loss (Figure 2 bottom), faster H and M growth resulted in more Byproduct, larger M populations, and consequently higher Product level. If H could evolve to grow faster than M, then increasing growth parameters could decrease community function due to H dominance (Figure 6A; Figure S17A; Figure S24B), although properly executed community selection can improve community function while promoting species coexistence (Figure 6B, C).

### Contrasting selection at different levels

From a “gene-centric” perspective, selection of individuals bears resemblance to selection of communities, as the survival of an individual relies on synergistic interactions between different genes at different activity levels. To ensure sufficient heredity between an individual and its offspring, elaborate cellular mechanisms have evolved, and they include cell cycle checkpoints to ensure accurate DNA replication and segregation [57], small RNA-mediated silencing of transposons [58], and CRISPR-Cas degradation of foreign viral DNA [59]. In community selection, heredity-enhancing mechanisms such as stable species ratio (Figure 2A bottom panel) could already be in place or arise due to mutations that affect species interactions. If a mechanism such as endosymbiosis should evolve in response to community selection, then community selection could transition to individual selection.

Group selection, and in a related sense, kin selection [39, 40, 60, 41, 42, 43, 44, 45, 46, 47, 48, 49, 50, 51, 52], have been extensively examined to explain, for example, the evolution of traits that lower individual fitness but increase the success of a group (e.g. sterile ants helping the survival of an ant colony). Note that the term “group selection” has often been used to describe individual selection in spatially-structured populations without group births and deaths, although such usage has been criticized [61]. Interestingly, artificial group selection can sometimes be viewed as artificial individual selection. For example, when Newborn groups start with a single contributor, then artificial group selection is equivalent to artificial individual selection where the trait under selection is the founder’s ability to produce a product over time as it grows into a population. On the other hand, if group function relies on distinct genotypes interacting synergistically, then group selection is similar to community selection.

Group selection can be thought of as a special case of community selection, except that group function can arise as a single founder multiplies. Therefore, group selection and community selection are similar in some aspects, but differ in other aspects. First, in both group selection and community selection, Newborn size must not be too large [62, 63] and maturation time must not be too long. Otherwise, all entities (groups or communities) will accumulate non-contributors in a similar fashion, and this low inter-entity variation impedes selection (Price equation [53]; Figure S1B; Figure S7). Second, species interactions in a community could drive species composition to a value sub-optimal for community function ([64]). This could also occur during artificial group selection if the founder genotype gives rise to sub-populations of distinct phenotypes interacting synergistically to generate group function (e.g. the growth of cyanobacteria filaments relying on differentiating into nitrogen-fixing cells and photosynthetic cells [65]). Otherwise, the problem of sub-optimal composition does not exist for group selection. Finally, in group selection, when a Newborn group starts with a small number of individuals (e.g. one individual), a fraction of Newborn groups of the next cycle will be highly similar to the original Newborn group (Figure S1B, bottom panel). This facilitates group selection. In contrast, when a Newborn community starts with a small number of total individuals, large stochastic fluctuations in Newborn composition can interfere with community selection (Figure 3). In the extreme case, a member species may even be lost by chance. Even if a fixed biomass of each species is sorted into Newborns, heredity is much reduced during community selection due to random sampling of genotypes from member species. For example, if Newborn groups are initiated with a single contributor and if the highest-functioning Adult group has accumulated 50% non-contributors, then 50% Newborn groups of the next cycle will be initiated with a single contributor and thus display group function. In contrast, if a Newborn community starts with one contributor from each of the two species and 50% non-contributors have accumulated in each species, then only 50%×50%= 25% Newborn communities of the next cycle will be initiated with contributors from both species and display community function.

### Community function may not be maximized through pre-optimizing member species in monocultures

If we know how each member species contributes to community function, might we pre-optimize member species in monocultures before assembling them into high-functioning communities? This turned out to be challenging due to the difficulty of recapitulating community dynamics in monocultures. For example, when we tried to improve M’s *f*_*P*_ by artificial group selection, *f*_*P*_ failed to improve to 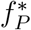 optimal for community function. Specifically, we started with *n*_*tot*_ of 100 Newborn M groups, each inoculated with one M cell (to facilitate group selection, Figure S1B bottom panel) [62]. We would supply each Newborn M group with the same amount of Resource as we would for H-M communities and excess Byproduct (since it is difficult to reproduce community Byproduct dynamics in M groups). After incubating these M groups for the same maturation time *T*, the group with the highest level of Product, *P*(*T*), would be chosen and reproduced into Newborn M groups for the next cycle. M’s growth parameters improved to evolutionary upper bounds (Figure S25A), since with Resource and Byproduct in excess, more M cells would lead to higher group function. When growth parameters were fixed to evolutionary upper bounds, optimal *f*_*P*_ for monoculture *P*(*T*) could be calculated to occur at an intermediate value (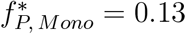; Figure S25B). Optimal group function was indeed realized during selection (Figure S25A). However, optimal *f*_*P*_ for group function is much lower than optimal *f*_*P*_ for community function (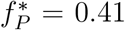; see Methods Section 8 for an explanation). Thus, optimizing monoculture activity does not necessarily lead to optimized community function.

### Implications of our work

In light of this study, we offer an alternative interpretation of previous work. In the work of [10], authors tested two selection regimens with Newborn sizes differing by 100-fold. The authors hypothesized that smaller Newborns would have a high level of variation which should facilitate selection. However, the hypothesis was not corroborated by experiments. As a possible explanation, the authors invoked the “butterfly effect” (the sensitivity of chaotic systems to initial conditions). Our results suggest that even for non-chaotic systems like the H-M community, selection could fail due to interference from non-heritable variations. This is because in Newborns with small sizes, fluctuations in community composition can be large, which compromises heredity of community trait.

A general implication of our work is that before launching a selection experiment, one should carefully design the selection regimen. Although some community functions are not sensitive to fluctuations in Newborn biomass compositions (e.g. steady state ratio or growth rate of mutualistic communities [66, 37]), many are. How might we check? The first method involves estimating the “signal to noise” ratio: one could initiate Newborn community replicates and measure community functions using the most precise method (e.g. cell-sorting during Newborn formation; many repeated measurements of community function). Despite this, some levels of non-heritable variations in community function are inevitable due to, for example, non-genetic phenotypic variations among cells [67] or stochasticity in cell birth and death. If “noises” (variations among community replicates) are small compared to “signals” (variations among communities with different genotypes and thus different community functions), then one can test and possibly adopt less precise procedures (e.g. cell culture pipetting during Newborn formation; fewer repeated measurements of community function). The second method involves estimating the heritability empirically if significant variations in community function naturally arise within the first few cycles. In this case, one could experimentally evaluate whether community functions of the previous cycle are correlated with community functions of the current cycle (across independent lineages (similar to Figure 4). Regardless, given the ubiquitous nature of non-heritable variations in community function, the bet-hedging strategy should be tested.

Microbes can co-evolve with each other and with their host in nature [68, 69, 70]. Some have proposed that complex microbial communities such as the gut microbiota could serve as a unit of selection [16]. Our work suggests that if selection for a costly microbial community function should occur in nature, then mechanisms for suppressing non-heritable variations in community function should be in place.

### Future directions

Our work touched upon only the tip of the iceberg of community selection. We expect that certain rules will be insensitive to details of a community. For example, reducing non-heritable variations in community function and judicious bet-hedging can facilitate community selection. Regardless, many fascinating questions remain. Here, we outline a few:

1. How might we best “hedge bets” when choosing Adult communities to reproduce? We have chosen top ten communities to contribute an equal number of Newborns, but alternative strategies (e.g. allowing higher-functioning Adults to contribute more Newborns) may work better. This aspect might be explored through applying population genetics theories, which has considered the balance between the strength of selection and variation among the individuals.
2. How might migration (community mixing) impact selection? Here, we did not consider migration. Excessive migration could deter community selection by allowing fast-growing non-contributors to spread. However, by combining the best genotypes of multiple member species, migration could speed up community selection, much like the effects of sexual recombination in the evolution of finite populations [71].
3. How might interaction structure affect selection efficacy? We have shown that our conclusions hold for two-species communities engaging in commensalism or mutualism. We have also shown that our conclusions hold regardless of whether the two species can evolve to coexist or not. The next step would be to test other types of interactions and complex interaction networks. For example, when species mutually inhibit each other, multistability could arise such that species dominance [72] and thus community function are sensitive to stochastic fluctuations in Newborn species density. How might multistability affect community selection? For a complex community, there might be multiple optima of community function, especially when the optimal community function requires multiple species with partially redundant activities. Consider the community function of waste degradation where one species degrades high-concentration waste incompletely while a complementary species degrades waste thoroughly but requires low starting waste concentration. If during Newborn formation, any one species is lost by chance, then community function would be stuck at local sub-optima. In this case, “bet-hedging” with community migration (mixing) could recover the lost species and help community selection reach a higher optimum.
4. Develop a general theory to understand how the rate of community function improvement depends on variations and heredity in community function, which are in turn affected by experimental parameters including Newborn size, the number of generations during maturation, mutation rate, the total number of communities under selection, selection strength, migration frequency, precision in community function measurement, and fluctuations during community reproduction.
5. Experimentally test model predictions. The assay for community function should be fast and precise. If community function is sensitive to species biomass in Newborns, then member species should ideally be distinguishable by flow cytometer (e.g. different fluorescence or different scattering patterns). Note that cell-sorting only needs to be performed on several high-functioning communities, and thus would not be cost prohibitive.
6. Discover interaction mechanisms important for community function. Once high-functioning communities are obtained through selection, we could compare metagenomes of evolved communities with those of ancestral communities. This would illuminate species interactions that are important for community function.

## Methods

### 1 Equations

*H*, the biomass of H, changes as a function of growth and death,

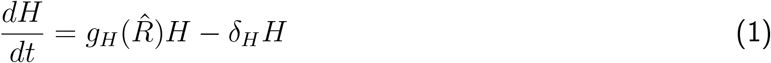

Grow rate *g*_*H*_ depends on the level of Resource 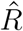 (hat 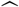 representing pre-scaled value) as described by the Monod growth model

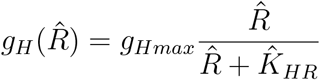

where 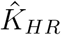 is the 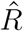 at which *g*_*Hmax*_/2 is achieved. *δ*_*H*_ is the death rate of H. Note that since agricultural waste is in excess, its level does not change and thus does not enter the equation.

*M*, the biomass of M, changes as a function of growth and death,

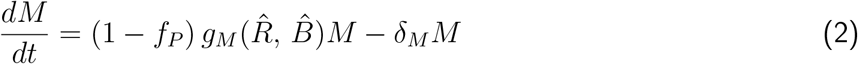

Total potential growth rate of M *g*_*M*_ depends on the levels of Resource and Byproduct (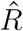 and 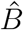) according to the Mankad-Bungay model [27] due to its experimental support:

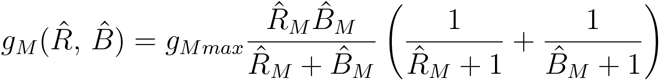

where 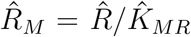 and 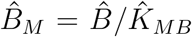 (Figure S3). 1 − *f*_*P*_ fraction of M growth is channeled to biomass increase. *f*_*P*_ fraction of M growth is channeled to making Product:

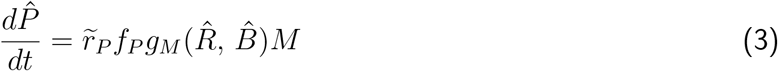

where 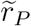 is the amount of Product made at the cost of one M biomass (tilde ~ representing scaling below and Table 1).

Resource 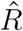 is consumed proportionally to the growth of M and H; Byproduct 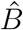 is released proportionally to H growth and consumed proportionally to M growth:

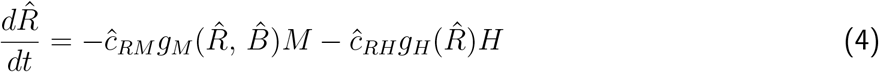

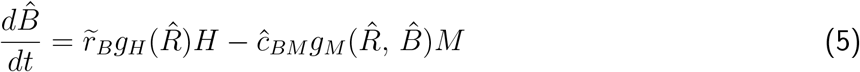

Here, 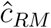 and 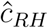 are the amounts of 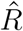 consumed per potential M biomass and H biomass, respectively. 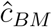 is the amount of 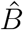 consumed per potential M biomass. 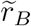 is the amount of 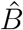 released per H biomass grown. Our model assumes that Byproduct or Product is generated proportionally to H or M biomass grown, which is reasonable given the stoichiometry of metabolic reactions and experimental support [73]. The volume of community is set to be 1, and thus cell or metabolite quantities (which are considered here) are numerically identical to cell or metabolite concentrations.

In equations above, scaling factors are marked by “~”, and will become 1 after scaling. Variables and parameters with hats will be scaled and lose their hats afterwards. Variables and parameters without hats will not be scaled. We scale Resource-related variable 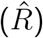 and parameters 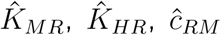 and 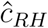) against 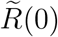 (Resource supplied to Newborn), Byproduct-related variable 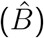 and parameters (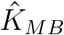 and 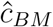) against 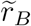 (amount of Byproduct released per H biomass grown), and Product-related variable 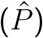 against 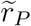 (amount of Product made at the cost of one M biomass). For biologists who usually think of quantities with units, the purpose of scaling (and getting rid of units) is to reduce the number of parameters. For example, H biomass growth rate can be re-written as:

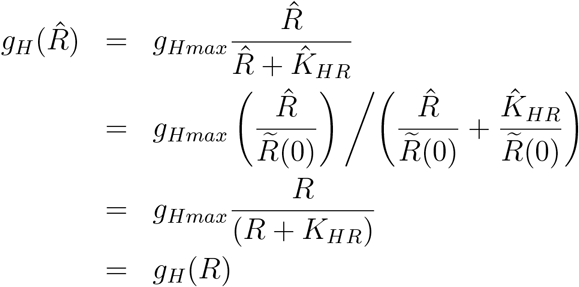

where 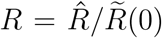 and 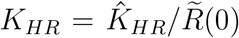. Thus, the unscaled 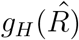 and the scaled *g*_*H*_(*R*) share identical forms (Figure S3). After scaling, the value of 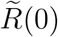 becomes irrelevant (1 with no unit). Similarly, since 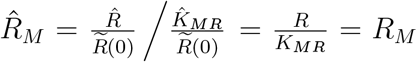 and 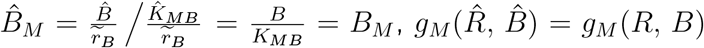 (Figure S4).

Thus, scaled equations are

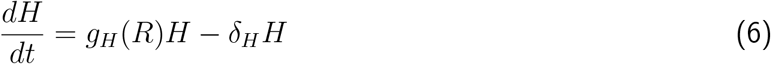

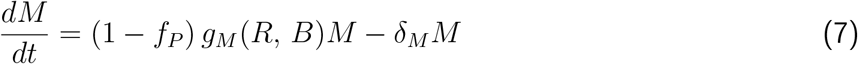

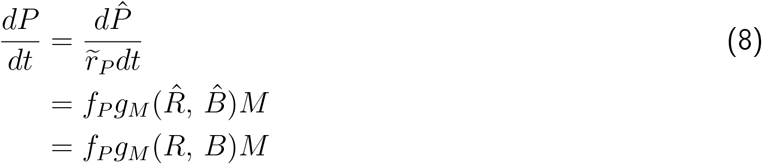

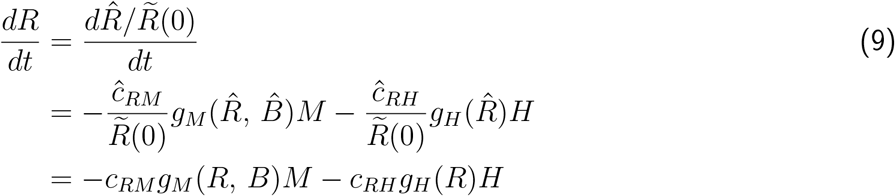

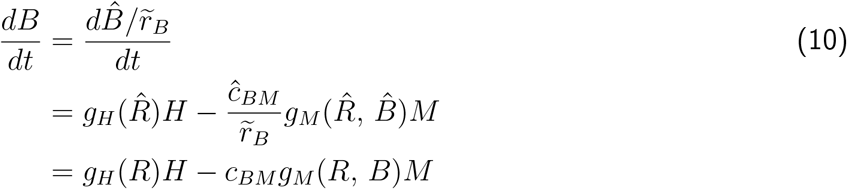

We have not scaled time here, although time can also be scaled by, for example, the community maturation time. Here, time has the unit of unit time (e.g. hr), and to avoid repetition, we often drop the time unit. After scaling, values of all parameters (including scaling factors) are in Table 1, and variables in our model and simulations are summarized in Table 2.

**Table 2:**
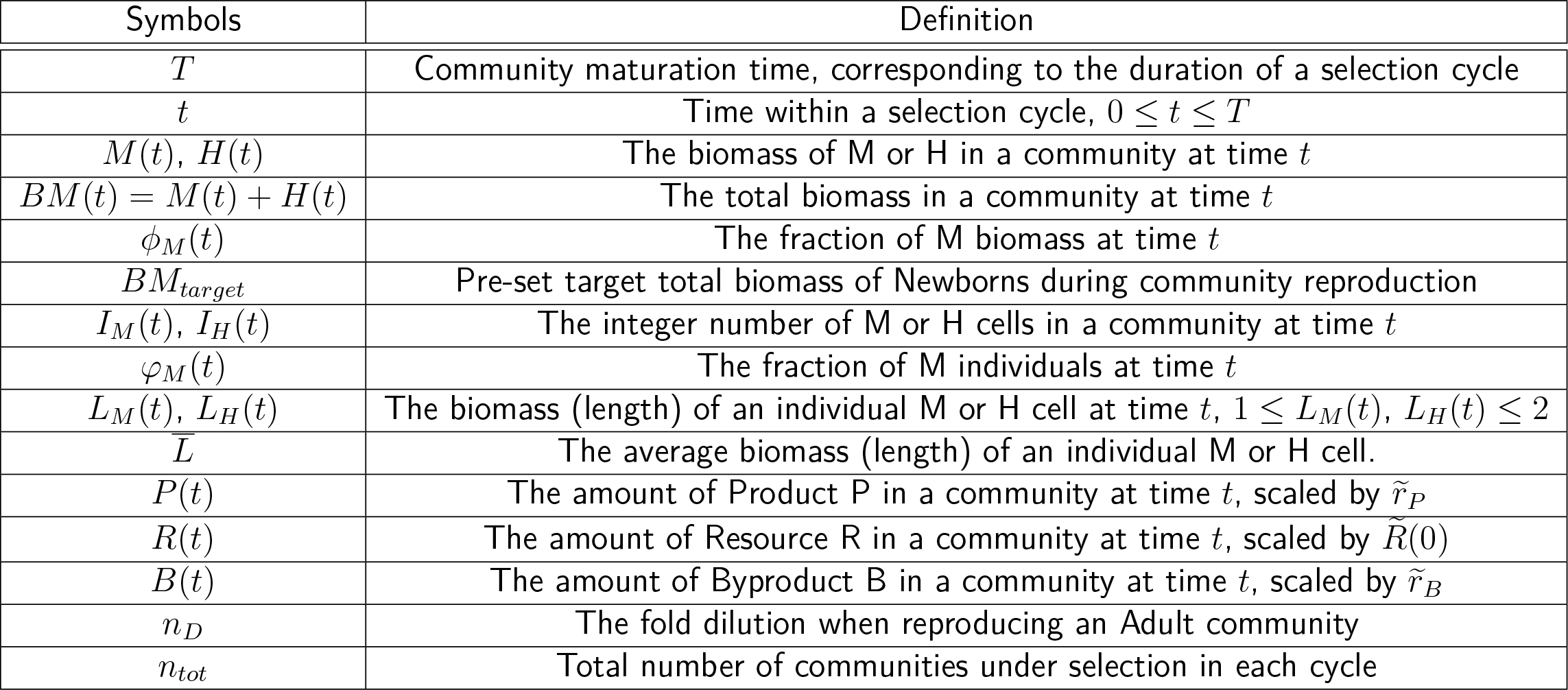
A summary of variables used in the simulation.

From Eq. 10:

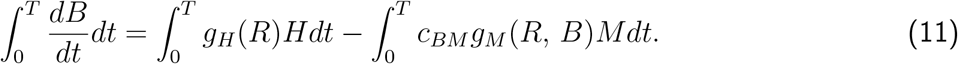

If we approximate Eq. 6-7 by ignoring the death rates so that 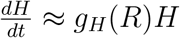 and 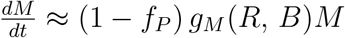, Eq. 11 becomes

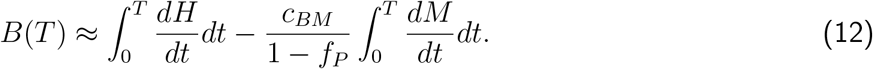

If B is the limiting factor for the growth of M so that B is mostly depleted, we can approximate *B* ≈ 0. If *T* is large enough so that both M and H has multiplied significantly and *H*(*T*) ≫ *H*(0) and *M*(*T*) ≫ *M*(0), Eq. 12 becomes

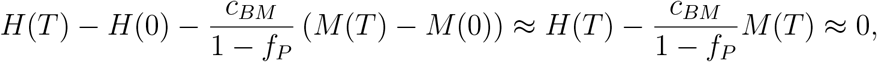

the M:H ratio at time *T* is

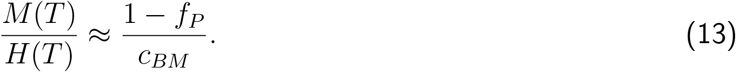

The steady state *ϕ*_*M*_, *ϕ*_*M*,*SS*_, is then

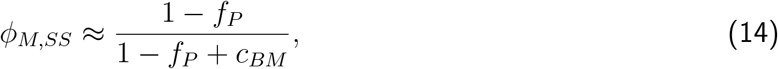

because if a community has *ϕ*_*M*_(0) = *ϕ*_*M*,*SS*_ at its Newborn stage, it has the same *ϕ*_*M*_(*T*) = *ϕ*_*M*,*SS*_ at its Adult stage.

In our simulations, because we supplied the H-M community with abundant R to avoid stationary phase, H grows almost at the maximal rate through *T* and releases B. If *f*_*P*_ is not too large (*f*_*P*_ < 0.4), which is satisfied in our simulations, M grows at a maximal rate allowed by B and keeps B at a low level. Thus, Eq. 14 is applicable and predicts the steady-state *ϕ*_*M*,*SS*_ well (see Figure S26). Note that significant deviation occurs when *f*_*P*_ > 0.4. This is because when *f*_*P*_ is large, M’s biomass does not grow fast enough to deplete B so that we cannot approximate *B*(*T*) ≈ 0 anymore.

### 2 Parameter choices

Our parameter choices are based on experimental measurements from a variety of organisms. Additionally, we chose growth parameters (maximal growth rates and affinities for metabolites) of ancestral and evolved H and M so that 1) the two species can coexist at a moderate ratio for a range of *f*_*P*_ over multiple selection cycles and 2) improving all growth parameters up to their evolutionary upper bounds generally improves community function (Methods Section 3). This way, we could simplify our simulation by fixing growth parameters at their respective evolutionary upper bounds. With only one mutable parameter (*f*_*P*_), we can identify the optimal 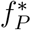 associated with maximal community function (Figure 2B).

For ancestral H, we set *g*_*Hmax*_ = 0.25 (equivalent to 2.8-hr doubling time if we choose hr as the time unit), *K*_*HR*_ = 1 and *c*_*RH*_ = 10^−4^ (both with unit of 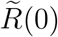) (Table 1). This way, ancestral H can grow by about 10-fold by the end of *T* = 17. These parameters are biologically realistic. For example, for a *lys-S. cerevisiae* strain with lysine as Resource, un-scaled Monod constant is 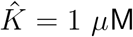, and consumption 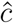 is 2 fmole/cell (Ref. [38]’s Figure 2 and Source Data 1, bioRxiv). Thus, if we choose 10 *μ*L as the community volume 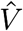 and 2 *μ*M as the initial Resource concentration, then 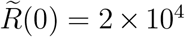 fmole. After scaling, 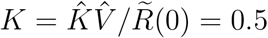 and 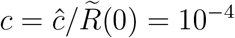, comparable to values in Table 1.

To ensure the coexistence of H and M, M must grow faster than H for part of the maturation cycle. Since we have assumed M and H to have the same affinity for R (Table 1), *g*_*Mmax*_ must exceed *g*_*Hmax*_, and M’s affinity for Byproduct (1/*K*_*MB*_) must be sufficiently large. Moreover, metabolite release and consumption need to be balanced to avoid extreme ratios between metabolite releaser and consumer. Thus for ancestral M, we chose *g*_*Mmax*_ = 0.58 (equivalent to a doubling time of 1.2 hrs). We set 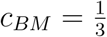 (units of 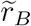), meaning that Byproduct released during one H biomass growth is sufficient to generate 3 potential M biomass, which is biologically achievable ([37, 74]). When we chose 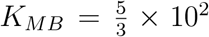 (units of 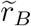), H and M can coexist for a range of *f*_*P*_ (Figure 2). This value is biologically realistic. For example, suppose that H releases hypoxanthine as Byproduct. A hypoxanthine-requiring *S. cerevisiae* M strain evolved under hypoxanthine limitation could achieve a Monod constant for hypoxanthine at 0.1 *μ*M (bioRxiv). If the volume of the community is 10 *μ*L, then 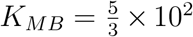 (units of 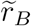) corresponds to an absolute release rate 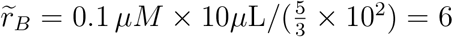 fmole per releaser biomass born. At 8 hour doubling time, this translates to 6 fmole/(1 cell × 8 hr) ≈ 0.75 fmole/cell/hr, within the ballpark of experimental observation (~0.3 fmole/cell/hr, bioRxiv). As a comparison, a lysine-overproducing yeast strain reaches a release rate of 0.8 fmole/cell/hr (bioRxiv) and a leucine-overproducing strain reaches a release rate of 4.2 fmole/cell/hr ([74]). Death rates *δ*_*H*_ and *δ*_*M*_ were chosen to be 0.5% of H and M’s respective upper bound of maximal growth rate, which are within the ballpark of experimental observations (e.g. the death rate of a *lys*- strain in lysine-limited chemostat is 0.4% of maximal growth rate, bioRxiv).

We assume that H and M consume the same amount of R per new cell (*c*_*RH*_ = *c*_*RM*_) since the biomass of various microbes share similar elemental (e.g. carbon or nitrogen) compositions [75]. Specifically, *c*_*RH*_ = *c*_*RM*_ = 10^−4^ (units of 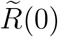), meaning that the Resource supplied to each Newborn community can yield a maximum of 10^4^ total biomass.

In some simulations (e.g. Figures 6, S8, S17), growth parameters (maximal growth rates *g*_*Mmax*_ and *g*_*Hmax*_ and affinities for nutrients 1/*K*_*MR*_, 1/*K*_*MB*_, and 1/*K*_*HR*_) and production cost parameter (0 ≤ *f*_*P*_ ≤ 1) were allowed to change from ancestral values during community maturation, since these phenotypes have been observed to rapidly evolve within tens to hundreds of generations ([29, 30, 31, 32]). For example, several-fold improvement in nutrient affinity and ~20% increase in maximal growth rate have been observed in experimental evolution [32, 30]. We therefore allowed affinities 1/*K*_*MR*_, 1/*K*_*HR*_, and 1/*K*_*MB*_ to increase by up to 3-fold, 5-fold, and 5-fold respectively, and allowed *g*_*Hmax*_ and *g*_*Mmax*_ to increase by up to 20%. These bounds also ensured that evolved H and M could coexist for *f*_*p*_ < 0.5, and that Resource was on average not depleted by *T* to avoid cells entering stationary phase.

We also simulated community selection where improved growth parameters could reduce community function (Figures 6 and S17). In this simulation, *g*_*Hmax*_ was allowed to increase by up to 220% and each Newborn community was supplied with R that can support up to 10^5^ cells (10 units of 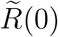).

Although maximal growth rate and nutrient affinity can sometimes show trade-off (e.g. Ref. [30]), for simplicity we assumed here that they are independent of each other. We held metabolite consumption (*c*_*RM*_, *c*_*BM*_, *c*_*RH*_) constant because conversion of essential elements such as carbon and nitrogen into biomass is unlikely to evolve quickly and dramatically, especially when these elements are not in large excess ([75]). Similarly, we held the scaling factors 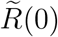 and 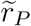 constant, assuming that they do not change rapidly during evolution due to stoichiometric constraints of biochemical reactions. We held death rates (*δ*_*M*_, *δ*_*H*_) constant because they are much smaller than growth rates in general and thus any changes are likely inconsequential.

### 3 Choosing growth parameter ranges so that we can fix growth parameters to upper bounds

Improving individual growth (maximal growth rate and affinity for metabolites) does not always lead to improved community function (Figures 6 and S17). However, we have chosen H and M growth parameters so that improving them from their ancestral values up to upper bounds generally improves community function (see below). When Newborn communities are assembled from “growth-adapted” H and M with growth parameters at upper bounds, two advantages are apparent.

First, after fixing growth parameters of H and M to their upper bounds, we can identify a locally maximal community function. Specifically, for a Newborn with total biomass *BM*(0) = 100 and fixed Resource *R*, we can calculate *P*(*T*) under various *f*_*P*_ and *ϕ*_*M*_(0), assuming that all M cells have the same *f*_*P*_. Since both numbers range between 0 and 1, we calculate *P*(*T*, *f*_*P*_ = 0.01 × *i*, *ϕ*_*M*_(0) = 0.01 × *j*) for integers *i* and *j* between 1 and 99. There is a single maximum for *P*(*T*) when *i* = 41 and *j* = 54. In other words, if M invests 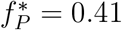 of its potential growth to make Product and if the fraction of M biomass in Newborn 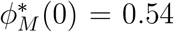, then maximal community function 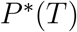 is achieved (Figure 2B; magenta dashed line in Figure 3).

Second, growth-adapted H and M are evolutionarily stable in the sense that deviations (reductions) from upper bounds will reduce both individual fitness and community function, and are therefore disfavored by intra-community selection and inter-community selection.

Below, we present evidence that within our parameter ranges (Table 1), improving growth parameters generally improves community function. When *f*_*P*_ is optimal for community function 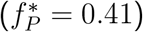, if we fix four of the five growth parameters to their upper bounds, then as the remaining growth parameter improves, community function increases (magenta lines in top panels of Figure S27). Moreover, mutants with a reduced growth parameter are out-competed by their growth-adapted counterparts (magenta lines in bottom panels of Figure S27).

When 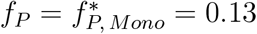 (optimal for M-monoculture function in Figure S25; the starting genotype for most community selection trials in this paper), community function and individual fitness generally increase as growth parameters improve (black dashed lines in Figure S27). However, when M’s affinity for Resource (1/*K*_*MR*_) is reduced from upper bound, fitness improves slightly (black dashed line in Panel J, Figure S27). Mathematically speaking, this is a consequence of the Mankad-Bungay model [27] (Figure S4B). Let *R*_*M*_ = *R/K*_*MR*_ and *B*_*M*_ = *B/K*_*MB*_. Then,

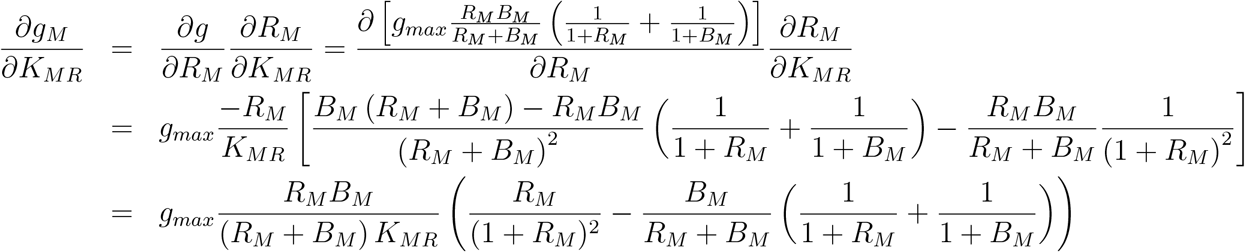

If *R*_*M*_ ≪ 1 ≪ *B*_*M*_ (corresponding to limiting R and abundant B),

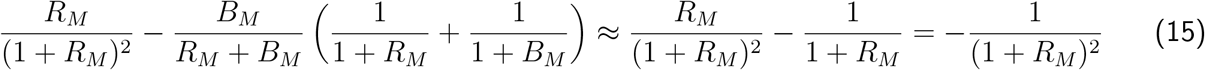

and thus 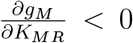. This is the familiar case where growth rate increases as the Monod constant decreases (i.e. affinity increases). However, if *B*_*M*_ ≪ 1 ≪ *R*_*M*_

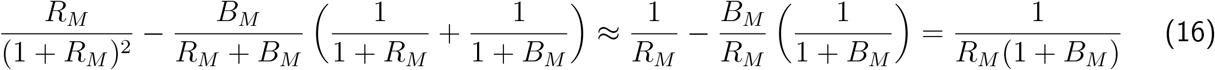

and thus 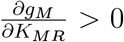. In this case, growth rate decreases as the Monod constant decreases (i.e. affinity increases). In other words, decreased affinity for the abundant nutrient improves growth rate. Transporter competition for membrane space [76] could lead to this result, since reduced affinity for abundant nutrient may increase affinity for rare nutrient. At the beginning of each cycle, R is abundant and B is limiting (Eq. 16). Therefore M cells with lower affinity for R will grow faster than those with higher affinity (Figure S28). At the end of each cycle, the opposite is true (Figure S28). As *f*_*P*_ decreases, M diverts more toward biomass growth and the first stage of B limitation lasts longer. Consequently, M can gain a slightly higher overall fitness by lowering the affinity for R (Figure S28A).

Regardless, decreased M affinity for Resource (1*/K_MR_*) only leads to a very slight increase in M fitness (Figure S27J) and a very slight decrease in *P*(*T*) (Figure S28B). Moreover, this only occurs at low *f*_*P*_ at the beginning of community selection, and thus may be neglected. Indeed, if we start all growth parameters at their upper bounds and *f*_*P*_ = 0.13, and perform community selection while allowing all parameters to vary (Figure S29), then 1*/K*_*MR*_ decreases somewhat, yet the dynamics of *f*_*P*_ is similar to when we only allow *f*_*P*_ to change (compare Figure S29D with Figure 3A).

### 4 Mutation rate and the distribution of mutation effects

Literature values of mutation rate and the distribution of mutation effects are highly variable. Below, we briefly review the literature and discuss rationales of our choices.

Among mutations, a fraction is neutral in that they do not affect the phenotype of interest. For example, the vast majority of synonymous mutations are neutral [77]. Furthermore, mutations with small effects may appear neutral, which can depend on the effective population size and selection condition. For example, at low population size due to genetic drift (i.e. changes in allele frequencies due to chance), a beneficial or deleterious mutation may not be selected for or selected against, and is thus neutral with respect to selection [78, 79]. As another example, the same mutation in an antibiotic-degrading gene can be neutral under low antibiotic concentrations, but deleterious under high antibiotic concentrations [80]. We term all these cases as “neutral” mutations.

Since a larger fraction of neutral mutations is equivalent to a lower rate of phenotype-altering mutations, our simulations define “mutation rate” as the rate of non-neutral mutations that either enhance a phenotype (“enhancing mutations”) or diminish a phenotype (“diminishing mutations”). Enhancing mutations of maximal growth rates (*g*_*Hmax*_ and *g*_*Mmax*_) and of nutrient affinities (1/*K_HR_*, 1/*K_M R_*, 1/*K*_*M B*_) enhance the fitness of an individual (“beneficial mutations”). In contrast, enhancing mutations in *f*_*P*_ diminish the fitness of an individual (“deleterious mutations”).

Depending on the phenotype, the rate of phenotype-altering mutations is highly variable. Although mutations that cause qualitative phenotypic changes (e.g. drug resistance) occur at a rate of 10^−8^~10^−6^ per genome per generation in bacteria and yeast [81, 82], mutations affecting quantitative traits such as growth rate occur much more frequently. For example in yeast, mutations that increase growth rate by ≥ 2% occur at a rate of ~ 10^−4^ per genome per generation (calculated from Figure 3 of Ref. [83]), and mutations that reduce growth rate occur at a rate of 10^−4^ ~ 10^−3^ per genome per generation [35, 84]. Moreover, mutation rate can be elevated by as much as 100-fold in hyper-mutators where DNA repair is dysfunctional [85, 86, 84]. In our simulations, we assume a high, but biologically feasible, rate of 2 × 10^−3^ phenotype-altering mutations per cell per generation per phenotype to speed up computation. At this rate, an average community would sample ~20 new mutations per phenotype during maturation. We have also simulated with a 100-fold lower mutation rate. As expected, evolutionary dynamics slowed down, but all of our conclusions still held (Figure S18).

Among phenotype-altering mutations, tens of percent create null mutants, as illustrated by experimental studies on protein, viruses, and yeast [33, 34, 35]. Thus, we assumed that 50% of phenotype-altering mutations were null (i.e. resulting in zero maximal growth rate, zero affinity for metabolite, or zero *f*_*P*_). Among non-null mutations, the relative abundances of enhancing versus diminishing mutations are highly variable in different experiments. It can be impacted by effective population size. For example, with a large effective population size, the survival rate of beneficial mutations is 1000-fold lower due to clonal interference (competition between beneficial mutations) [87]. The relative abundance of enhancing versus diminishing mutations also strongly depends on the starting phenotype [33, 80, 78]. For example with ampicillin as a substrate, the wild-type TEM-1 *β*-lactamase is a “perfect” enzyme. Consequently, mutations were either neutral or diminishing, and few enhanced enzyme activity [80]. In contrast with a novel substrate such as cefotaxime, the enzyme had undetectable activity, and diminishing mutations were not detected while 2% of tested mutations were enhancing [80]. When modeling H-M communities, we assumed that the ancestral H and M had intermediate phenotypes that can be enhanced or diminished.

We based our distribution of mutation effects on experimental studies where a large number of enhancing and diminishing mutants have been quantified in an unbiased fashion. An example is a study from the Dunham lab where the fitness effects of thousands of *S. cerevisiae* mutations were quantified under various nutrient limitations [36]. Specifically for each nutrient limitation, the authors first measured 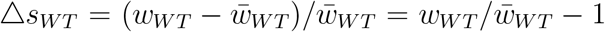, the deviation in relative fitness of thousands of barcoded wild-type control strains from the wild-type mean fitness (i.e. selection coefficients). Due to experimental noise, Δ*s*_*WT*_ is distributed with zero mean and non-zero variance. Then, the authors measured thousands of Δ*s*_*MT*_, each corresponding to the relative fitness change of a bar-coded mutant strain with respect to the mean of wild-type fitness (i.e. 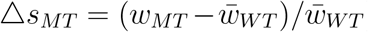). From these two distributions, we derived *μ*_Δ*s*_, the probability density function (PDF) of relative fitness change caused by mutations Δ*s* = Δ*s*_*MT*_ − Δ*s*_*WT*_ (see Figure S6 for interpreting PDF), in the following manner.

First, we calculated *μ_m_*(Δ*s*_*MT*_), the discrete PDF of the relative fitness change of mutant strains, with bin width 0.04. In other words, *μ_m_*(Δ*s*_*MT*_) =counts in the bin of [Δ*s*_*MT*_ − 0.02, Δ*s*_*MT*_ + 0.02] / total counts/0.04 where Δ*s*_*MT*_ ranges from −0.6 and 0.6 which is sufficient to cover the range of experimental outcome. The Poissonian uncertainty of *μ*_*m*_ is 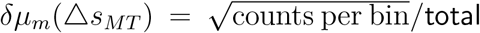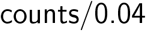. Repeating this process for the wild-type collection, we obtained the PDF of the relative fitness change of wild-type strains *μ_w_*(Δ*s*_*WT*_). Next, from *μ_w_*(Δ*s*_*WT*_) and *μ_m_*(Δ*s*_*MT*_), we derived *μ*_Δ*s*_(Δ*s*), the PDF of Δ*s* with bin width 0.04:

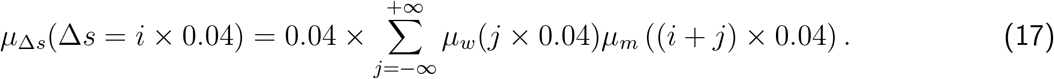

assuming that Δ*s*_*MT*_ and Δ*s*_*WT*_ are independent from each other. Here, *i* is an integer from −15 to 15. The uncertainty for *μ*_Δ*s*_ was calculated by propagation of error. That is, if *f* is a function of *x*_*i*_ (*i* = 1, 2, …, *n*), then *s*_*f*_, the error of *f*, is 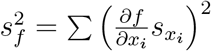, where 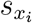 is the error or uncertainty of *x*_*i*_. Thus,

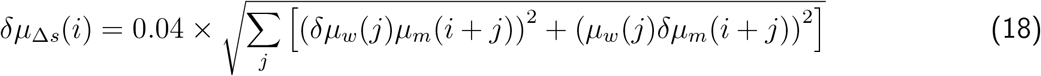

where *μ_w_*(*j*) is short-hand notation for *μ_w_*(Δ*s*_*WT*_ = *j* × 0.04) and so on. Our calculated *μ*_Δ*s*_(Δ*s*) with error bar of *δμ*_Δ*s*_ is shown in Figure S6.

Our reanalysis demonstrated that distributions of mutation fitness effects *μ*_Δ*s*_(Δ*s*) are largely conserved regardless of nutrient conditions and mutation types (Figure S6B). In all cases, the relative fitness changes caused by beneficial (fitness-enhancing) and deleterious (fitness-diminishing) mutations can be approximated by a bilateral exponential distribution with means *s*_+_ and *s*_−_ for the positive and negative halves, respectively. After normalizing the total probability to 1, we have:

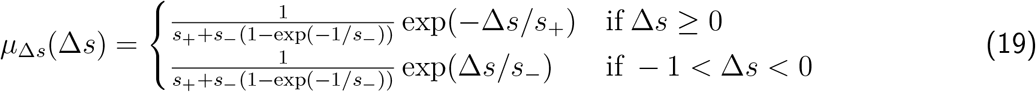

We fitted the Dunham lab haploid data (since microbes are often haploid) to Eq. 19, using *μ*_Δ*s*_(*i*)/*δμ*_Δ*s*_(*i*) as the weight for non-linear least squared regression (green lines in Figure S6B). We obtained *s*_+_ = 0.050 ± 0.002 and *s*_−_ = 0.067 ± 0.003.

Interestingly, exponential distribution described the fitness effects of deleterious mutations in an RNA virus remarkably well [33]. Based on extreme value theory, the fitness effects of beneficial mutations were predicted to follow an exponential distribution [88, 89], which has gained experimental support from bacterium and virus [90, 91, 92] (although see [93, 83] for counter examples). Evolutionary models based on exponential distributions of fitness effects have shown good agreements with experimental data [87, 94].

We have also simulated smaller average mutational effects based on measurements of spontaneous or chemically-induced (instead of deletion) mutations. For example, the fitness effects of nonlethal deleterious mutations in *S. cerevisiae* were mostly 1%~5% [35], and the mean selection coefficient of beneficial mutations in *E. coli* was 1%~2% [90, 87]. As an alternative, we also simulated with *s*_+_ = *s*_−_ = 0.02, and obtained the same conclusions (Figure S19).

### 5 Modeling epistasis on *f*_*P*_

Epistasis, where the effect of a new mutation depends on prior mutations (“genetic background”), is known to affect evolutionary dynamics. Epistatic effects have been quantified in various ways. Experiments on viruses, bacteria, yeast, and proteins have demonstrated that if two mutations were both deleterious or random, viable double mutants experienced epistatic effects that distributed nearly symmetrically around a value close to zero [95, 96, 97, 98, 99]. In other words, a significant fraction of mutation pairs show no epistasis, and a small fraction show positive or negative epistasis (i.e. a double mutant displays a stronger or weaker phenotype than expected from additive effects of the two single mutants). Epistasis between two beneficial mutations can vary from being predominantly negative [96] to being symmetrically distributed around zero [97]. Furthermore, a beneficial mutation tends to confer a lower beneficial effect if the background already has high fitness (“diminishing returns”) [100, 97, 101].

A mathematical model by Wiser et al. incorporates diminishing-returns epistasis [94]. In this model, beneficial mutations of advantage *s* in the ancestral background are exponentially distributed with probability density function (PDF) *α* exp(−*αs*), where 1/*α* > 0 is the mean advantage. After a mutation with advantage *s* has occurred, the mean advantage of the next mutation would be reduced to 1/[*α*(1 + *gs*)], where *g* > 0 is the “diminishing returns parameter”. Wiser et al. estimates *g* ≈ 6. This model quantitatively explains the fitness dynamics of evolving *E. coli* populations.

Based on the above experimental and theoretical literature, we modeled epistasis on *f*_*P*_ in the following manner. Let the relative mutation effect on *f*_*P*_ be Δ*f*_*P*_ = (*f_P,mut_* − *f*_*P*_)/*f*_*P*_ (note Δ*f_P_* ≥ −1). Then, *μ*(Δ*f_P_*, *f*_*P*_), the PDF of Δ*f*_*P*_ at the current *f*_*P*_ value, is described by a form similar to Eq. 19:

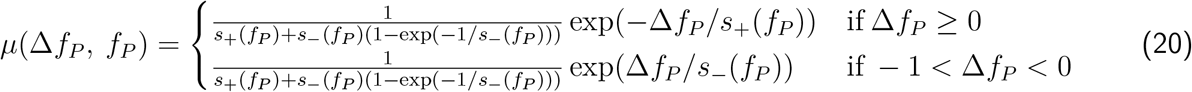

Here, *s*_+_(*f*_*P*_) and *s_−_*(*f*_*P*_) are respectively the mean Δ*f*_*P*_ for enhancing and diminishing mutations at current *f*_*P*_. We assigned *s*_+_(*f*_*P*_) = *s*_+*init*_/(1 + *g* × (*f_P_ /f_P, init_* − 1)), where *f*_*P, init*_ is the *f*_*P*_ of the initial background in a community selection simulation 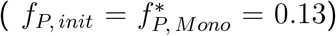, *s*_+*init*_ is the mean enhancing Δ*f*_*P*_ occurring in the initial background, and 0 < *g* < 1 is the epistatic factor. Similarly, *s_−_*(*f*_*P*_) = *s_−init_* ×(1+*g* ×(*f*_*P*_ / *f*_*P*, *init*_ − 1)) is the mean |Δ*f*_*P*_| for diminishing mutations at current *f*_*P*_. In the initial background, since *f*_*P*_ = *f_P, init_*, we have *s*_+_(*f*_*P*_) = *s*_+*init*_ and *s_−_*(*f*_*P*_) = *s*_−*init*_ (*s*_+*init*_ = 0.050 and *s*_*−init*_ = 0.067 in Figure S6). Consistent with the diminishing returns principle, for subsequent mutations that alter *f*_*P*_, if current *f*_*P*_ > *f*_*P,init*_, then a new enhancing mutation became less likely and its mean effect smaller, while a new diminishing mutation became more likely and its mean effect bigger (ensured by *g* > 0; Figure S20 right panel). Similarly, if current *f*_*P*_ < *f*_*P,init*_, then a new enhancing mutation became more likely and its mean effect bigger, while a diminishing mutation became less likely and its mean effect smaller (ensured by 0 < *g* < 1; Figure S20 left panel). In summary, our model captured not only diminishing-returns epistasis, but also our understanding of mutational effects on protein stability [78].

### 6 Simulation code of community selection

As described in the main text, our simulations tracked the biomass and phenotypes of individual cells as well as the amounts of Resource, Byproduct, and Product in each community throughout community selection. Cell biomass growth, cell division, and changes in chemical concentrations were calculated deterministically. Stochastic processes including cell death, mutation, and the partitioning of cells of a chosen Adult community into Newborn communities were simulated using the Monte Carlo method.

Specifically, each simulation was initialized with a total of *n*_*tot*_ = 100 Newborn communities with identical configuration:

- each community had 100 total cells of biomass 1. Thus, total biomass *BM* (0) = 100.
- 40 cells were H. 60 cells were M with identical *f*_*P*_. Thus, M biomass *M*(0) = 60 and fraction of M biomass *ϕ*_*M*_(0) = 0.6.

Our community selection simulations did not consider mutations arising during pre-growth prior to inoculating Newborns of the first cycle, because incorporating pre-growth had little impact on evolution dynamics (Figure S30). Thus, all M cells have the same phenotype, and all H cells have the same phenotype.

At the beginning of each selection cycle, a random number was used to seed the random number generator for each Newborn community. This number was saved so that the maturation of each Newborn community can be replayed. In most simulations, the initial amount of Resource was 1 unit of 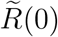 unless otherwise specified, the initial Byproduct was *B*(0) = 0 and the initial Product *P* (0) = 0.

The maturation time *T* was divided into time steps of Δ*τ* = 0.05. Resource *R*(*t*) and Byproduct *B*(*t*) during each time interval [*τ*, *τ* + Δ*τ*] were calculated by solving the following equations (similar to Eqs. 9-10) using the initial condition *R*(*τ*) and *B*(*τ*) via the ode23s solver in Matlab:

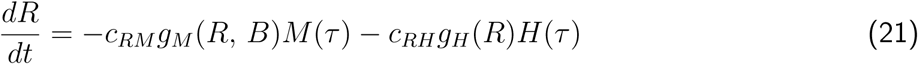

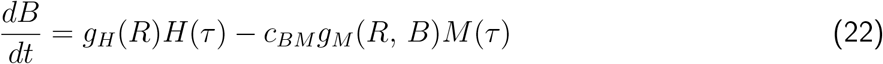

where *M*(*τ*) and *H*(*τ*) were the biomass of M and H at time *τ* (treated as constants during time interval [*τ*, *τ* + Δ*τ*]), respectively. The solutions from Eq. 21 and 22 were used in the integrals below to calculate the biomass growth of H and M cells.

Suppose that H and M were rod-shaped organisms with a fixed diameter. Thus, the biomass of an H cell at time *τ* could be written as the length variable *L*_*H*_(*τ*). The continuous growth of *L*_*H*_ during *τ* and *τ* + Δ*τ* could be described as

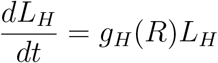

or

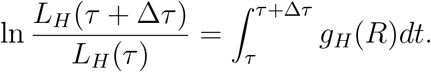

Thus,

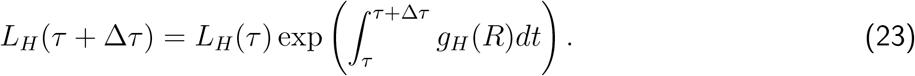

Similarly, let the length of an M cell be *L*_*M*_(*τ*). The continuous growth of M could be described as

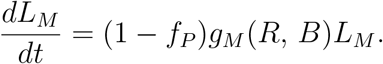

Thus for an M cell, its length *L*_*M*_(*τ* + Δ*τ*) could be described as

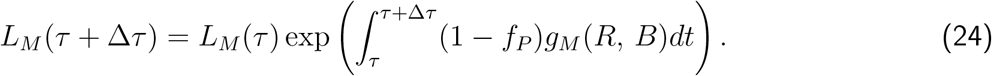

From Eq. 7 and 8, within Δ*τ*,

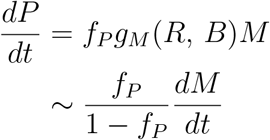

and therefore

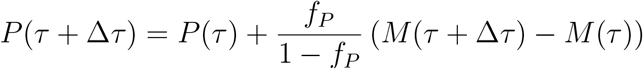

where *M*(*τ* + Δ*τ*) = Σ*L*_*M*_ (*τ* + Δ*τ*) represented the sum of the biomass (or lengths) of all M cells at *τ* + Δ*τ*.

At the end of each Δ*τ*, each H and M cell had a probability of *δ*_*H*_ Δ*τ* and *δ*_*M*_ Δ*τ* to die, respectively. This was simulated by assigning a random number between [0, 1] for each cell. Cells assigned with a random number less than *δ*_*H*_ Δ*τ* or *δ*_*M*_ Δ*τ* then got eliminated. For surviving cells, if a cell’s length ≥ 2, this cell would divide into two cells with half the original length.

After division, each mutable phenotype of each cell had a probability of *P_mut_* to be modified by a mutation (Methods, Section 4). As an example, let’s consider mutations in *f*_*P*_. If a mutation occurred, then *f*_*P*_ would be multiplied by (1 + Δ*f*_*P*_), where Δ*f*_*P*_ was determined as below.

First, a uniform random number *u*_1_ between 0 and 1 was generated. If *u*_1_ ≤ 0.5, Δ*f*_*P*_ = −1, which represented 50% chance of a null mutation (*f*_*P*_ = 0). If 0.5 < *u*_1_ ≤ 1, Δ*f*_*P*_ followed the distribution defined by Eq. 20 with *s*_+_(*f*_*P*_) = 0.05 for *f*_*P*_-enhancing mutations and *s_−_*(*f*_*P*_) = 0.067 for *f*_*P*_-diminishing mutations when epistasis was not considered (Methods, Section 4). In the simulation, Δ*f*_*P*_ was generated via inverse transform sampling. Specifically, *C*(Δ*f*_*P*_), the cumulative distribution function (CDF) of Δ*f*_*P*_, could be found by integrating Eq. 19 from −1 to Δ*f*_*P*_:

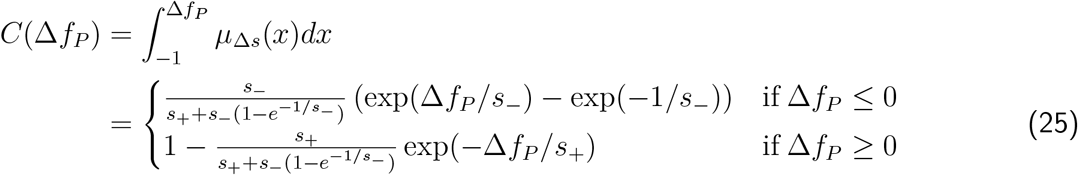

The two parts of Eq. 25 overlap at *C*(Δ*f*_*P*_ = 0) = *s_−_* (1−exp(−1/*s*_−_))/ [*s*_+_ + *s*_−_ (1−exp(−1/*s*_−_))]. In order to generate Δ*f*_*P*_ satisfying the distribution in Eq. 19, a uniform random number *u*_2_ between 0 and 1 was generated and we set *C*(Δ*f*_*P*_) = *u*_2_. Inverting Eq. 25 yielded

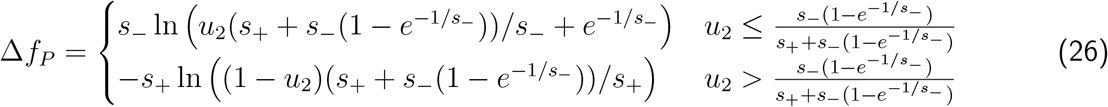

When epistasis was considered, *s*_+_(*f*_*P*_) = *s*_+*init*_/(1 + *g* × (*f_P_ /f_P, init_* − 1)) and *s*_−_ (*f*_*P*_) = *s*_−*init*_×(1 + *g* × (*f_P_ /f_P, init_ − 1*)) were used in Eq. 26 to calculated Δ*f*_*P*_ for each cell. (Methods Section 5).

If a mutation increased or decreased the phenotypic parameter beyond its bound (Table 1), the phenotypic parameter was set to the bound value.

The above growth/death/division/mutation cycle was repeated from time 0 to *T*. Note that since the size of each M and H cell can be larger than 1, the integer numbers of M and H cells, *I_M_* and *I_H_*, are generally smaller than the numerical values of biomass *M* and *H*, respectively. At the end of *T*, Adult communities were sorted according to their *P*(*T*) values. The Adult community with the highest *P*(*T*) (or a randomly-chosen Adult in control simulations) was chosen for reproduction.

Before community reproduction, the current random number generator state was saved so that the random partitioning of Adult communities could be replayed. To mimic partitioning Adult communities via pipetting into Newborn communities with an average total biomass of *BM_target_*, we first calculated the fold by which this Adult would be diluted as *n*_*D*_ = ⌊(*M*(*T*) + *H*(*T*)) /*BM_target_*⌋. Here *BM_target_* = 100 was the pre-set target for Newborn total biomass, and ⌊*x*⌋ is the floor (round down) function that generates the largest integer that is smaller than *x*. If the Adult community had *I_H_* (*T*) H cells and *I_M_* (*T*) cells, *I_H_* (*T*) + *I_M_* (*T*) random integers between 1 and *n_D_* were uniformly generated so that each M and H cell was assigned a random integer between 1 and *n_D_*. All cells assigned with the same random integer were then assigned to the same Newborn, generating *n_D_* newborn communities. This partition regimen can be experimentally implemented by pipetting 1/*n_D_* volume of an Adult community into a new well. If *n_D_* was less than *n_tot_* (the total number of communities under selection), all *n_D_* newborn communities were kept and the Adult with the next highest function was partitioned to obtain an additional batch of Newborns until we obtain *n_tot_* Newborns. The next cycle then began.

To fix *BM*(0) to *BM_target_* and *ϕ_M_*(0) to *ϕ_M_*(*T*) of the parent Adult, the code randomly assigned M cells from the chosen Adult until the total biomass of M came closest to *BM_target_ϕ_M_* (*T*) without exceeding it. H cells were assigned similarly. Because each M and H cells had a length between 1 and 2, the biomass of M could vary between *BM_target_ϕ_M_*(*T*) − 2 and *BM_target_ϕ_M_*(*T*) and the biomass of H could vary between *BM_target_*(1 − *ϕ_M_*(*T*)) − 2 and *BM_target_*(1 − *ϕ_M_*(*T*)). Variations in *BM*(0) and *ϕ_M_*(0) were sufficiently small so that community selection improved *f_P_*(*T*) (Figure 3D and E). We also simulated sorting cells where H and M cell numbers (instead of biomass) were fixed in Newborns. Specifically, ⌊*BM_target_φ*_*M*_(*T*)/1.5⌋ M cells and ⌊*BM_target_*(1 − *φ_M_*(*T*))/1.5⌋ H cells were sorted into each Newborn community, where we assumed that the average biomass of a cell was 1.5, and *υ_M_*(*T*) = *I_M_*(*T*)/(*I_M_*(*T*) + *I_H_*(*T*)) was calculated from cell numbers in the parent Adult community. We obtained the same conclusion (Figure S13, right panels).

To fix Newborn total biomass *BM*(0) to the target total biomass *BM_target_* while allowing *ϕ_M_*(0) to fluctuate (Figure S12 left panels), H and M cells were randomly assigned to a Newborn community until *BM*(0) came closest to *BM_target_* without exceeding it (otherwise, *P*(*T*) might exceed the theoretical maximum). For example, suppose that a certain number of M and H cells had been sorted into a Newborn so that the total biomass was 98.6. If the next cell, either M or H, had a biomass of 1.3, this cell would go into the community so that the total biomass would be 98.6 + 1.3 = 99.9. However, if a cell of mass 1.6 happened to be picked, this cell would not go into this community so that this Newborn had a total biomass of 98.6 and the cell of mass 1.6 would go to the next Newborn. Thus, each Newborn might not have exactly the biomass of *BM_target_*, but rather between *BM_target_* − 2 and *BM_target_*. Experimentally, total biomass can be determined from the optical density, or from the total fluorescence if cells are fluorescently labeled ([54]). To fix the total cell number (instead of total biomass) in a Newborn, the code randomly assigned a total of ⌊*BM_target_/*1.5⌋ cells into each Newborn, assuming an average cell biomass of 1.5. We obtained the same conclusion, as shown in Figure S13 left panel.

To fix *ϕ_M_*(0) to *ϕ_M_*(*T*) of the chosen Adult community from the previous cycle while allowing *BM*(0) to fluctuate (Figure S12 right panels), the code first calculated dilution fold *n_D_* in the same fashion as mentioned above. If the Adult community had *I_H_*(*T*) H cells and *I_M_*(*T*) cells, *I_M_*(*T*) random integers between [1, *n_D_*] were then generated for each M cell. All M cells assigned the same random integer joined the same Newborn community. The code then randomly dispensed H cells into each Newborn until the total biomass of H came closest to *M*(0)(1 − *ϕ_M_*(*T*))/*ϕ_M_*(*T*) without exceeding it, where *M*(0) was the biomass of all M cells in this Newborn community. Again, because each M and H had a biomass (or length) between 1 and 2, *ϕ_M_*(0) of each Newborn community might not be exactly *ϕ_M_*(*T*) of the chosen Adult community. We also performed simulations where the ratio between M and H cell numbers in the Newborn community, *I_M_*(0)/*I_H_*(0), was set to *I_M_*(*T*)/*I_H_*(*T*) of the Adult community, and obtained the same conclusion (Figure S13 center panels).

### 7 Problems associated with alternative definitions of community function and alternative means of reproducing an Adult

Here we describe problems associated with two alternative definitions of community function and one alternative method of community reproduction.

One alternative definition of community function is Product per M biomass in an Adult community: *P*(*T*)/*M*(*T*). To illustrate problems with this definition, let’s calculate *P*(*T*)/*M*(*T*) assuming that cell death is negligible. From Eq. 7 and 8,

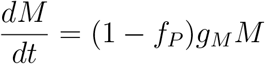

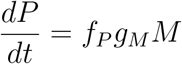

where biomass growth rate *g*_*M*_ is a function of *B* and *R*. Thus,

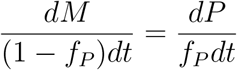

and we have

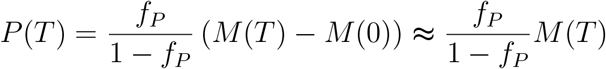

if *M*(*T*) ≫ *M*(0) (true if *T* is long enough for cells to double at least three or four times).

If we define community function as 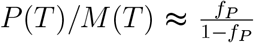, then higher community function requires higher 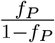 or higher *f*_*P*_. However, if we select for very high *f*_*P*_, then M can go extinct (Figure 2).

In our community selection scheme, the average total biomass of Newborn communities was set to a constant *BM*_*target*_. Alternatively, each Adult community can be partitioned into a constant number of Newborn communities. If Resource is not limiting, there is no competition between H and M, and *P*(*T*) increases as *M*(0) and *H*(0) increase. Therefore, selection for higher *P*(*T*) results in selection for higher Newborn total biomass (instead of higher *f*_*p*_). This will continue until Resource becomes limiting, and then communities will get into the stationary phase.

### 8 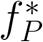 is smaller for M group than for H-M community

For groups or communities with a certain *∫*_*T*_*g*_*M*_*dt*, we can calculate *f*_*P*_ optimal for community function from Eq. **??** by setting

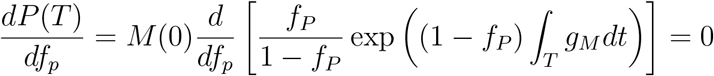

We have

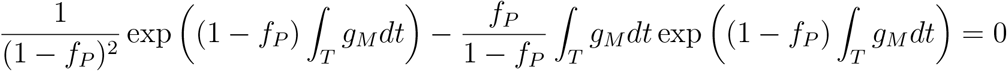

or

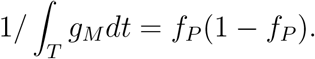

If *∫*_*T*_*g*_*M*_*dt* ≫ 1, *f*_*P*_ is very small, then the optimal *f*_*P*_ for *P*(*T*) is

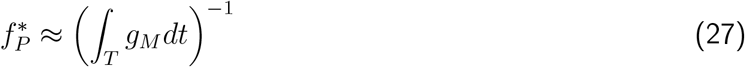

M grows faster in monoculture than in community because B is supplied in excess in monoculture while in community, H-supplied Byproduct is initially limiting. Thus, *g dt* is larger in monoculture than in community. According to Eq. 27, 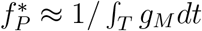 is smaller for monoculture than for community.

### 9 Stochastic fluctuations during community reproduction

The number of cells in a Newborn community is approximately 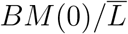, where 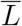 is the average biomass (or length) of M and H cells. This number fluctuates in a Poissonian fashion with a standard deviation of 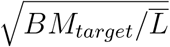. As a result, the biomass of a Newborn communities fluctuates around *BM*_*target*_ with a standard deviation of 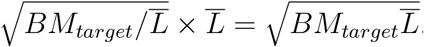.

Similarly, *M*(0) and *H*(0) fluctuate independently with a standard deviation of 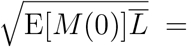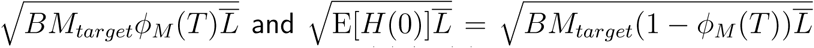, respectively, where “E” means the expected value. Therefore, *M*(0)*/H*(0) fluctuates with a variance of

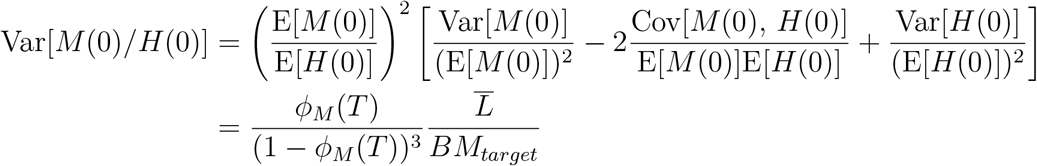

where “Cov” means covariance and “Var” means variance, and *ϕ*_*M*_(*T*) is the fraction of M biomass in the Adult community from which Newborns are generated.

### 10 Mutualistic H-M community

In the mutualistic H-M community, Byproduct inhibits the growth of H. According to [102], the growth rate of *E. coli* decreases exponentially as the exogenously added acetate concentration increases. Thus, we only need to modify the growth of H by a factor of exp(−*B/B*_0_) where *B* is the concentration of Byproduct and *B*_0_ is the concentration of Byproduct at which H’s growth rate is reduced by *e*^−^~0.37:

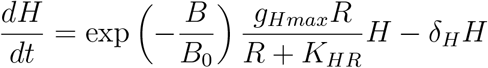

The larger *B*_0_, the less inhibitory effect Byproduct has on H and when *B*_0_ → +∞ Byproduct does not inhibit the growth of H. For simulations in Figure S22, we set *B*_0_ = 2*K*_*MB*_.

## Acknowledgment

We thank the following for discussions: Lin Chao (UCSD), Maitreya Dunham (UW Seattle), Corina Tarnita (Princeton), Harmit Malik (Fred Hutch), Jeff Gore (MIT), Daniel Weissman (Emory), Tony Long (UC Irvine), and Alvaro Sanchez (Yale). Some of these discussions took place at the 2017 “Eco-Evolutionary Dynamics in Nature and the Lab" workshop at Kavli Institute of Theoretical Physics, UC Santa Barbara, the 2017 “Systems Biology and Molecular Economy of Microbial Communities” workshop at the International Centre for Theoretical Physics, Trieste, Italy, and the 2018 “Physical Principles Governing the Organization of Microbial Communities” workshop at the Aspen Center for Physics, Colorado, USA. We thank Chichun Chen, Bill Hazelton, Samuel Hart, David Skelding, Doug Jackson, Maxine Linial, Delia Pinto-Santini, Kirill Korolev (Boston University), and Alex Sigal (K-RITH) for feedback on the manuscript. We are particularly indebted to Jim Bull (UT Austin) and reviewers (Sara Mitri, James Boedicker, and two anonymous reviewers) who generously provided detailed critiques. This research was facilitated by the High Performance Computing Shared Resource of the Fred Hutch (P30 CA015704).

## Supplementary Figures

**Figure S1:**
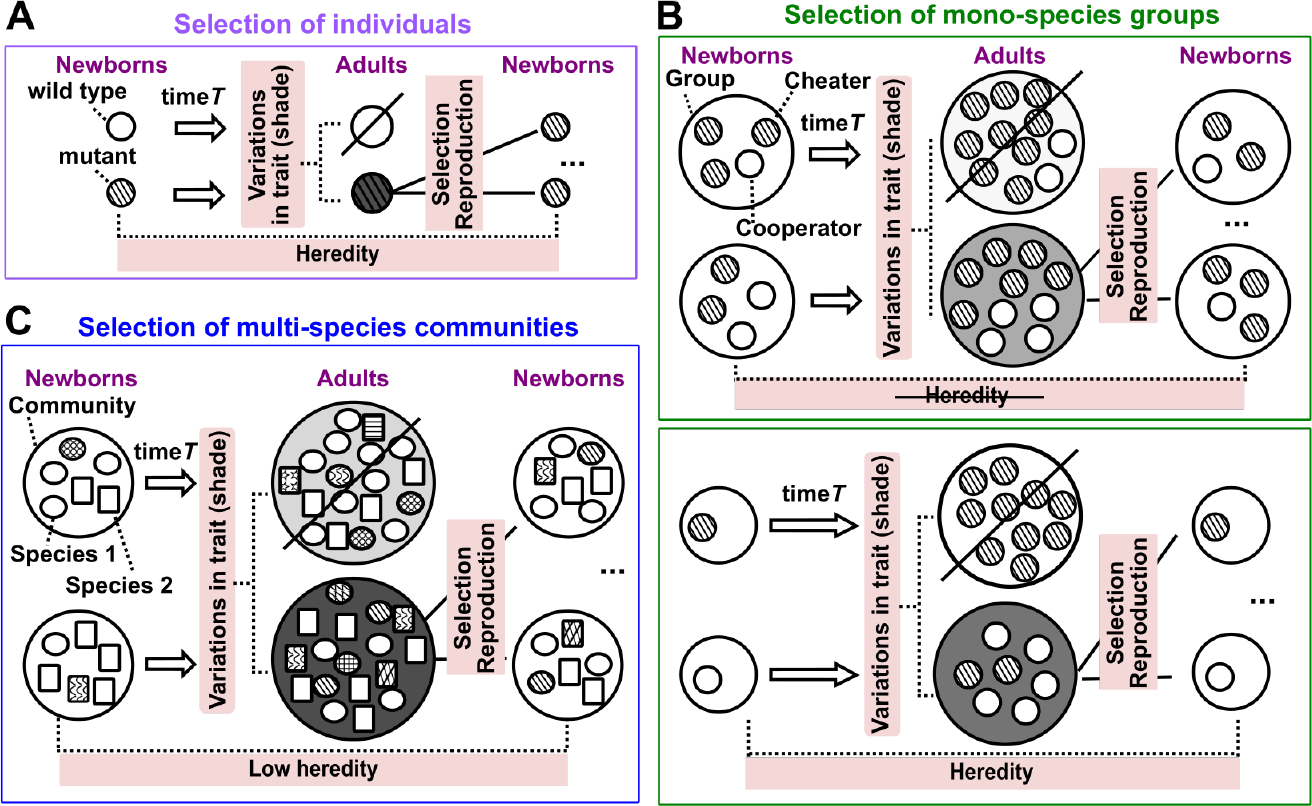
Artificial selection is more challenging for multi-species communities than for individuals or mono-species groups. Artificial selection can be applied to any population of entities [103]. An entity can be an individual (**A**), a mono-species group (**B**), or a multi-species community (**C**). Unlike natural selection which selects for fastest-growing cells, artificial selection generally selects for traits that are costly to individuals. In each selection cycle, a population of “Newborn” entities grow for maturation time *T* to become “Adults”. Adults expressing a higher level of the trait of interest (darker shade) are chosen to reproduce. An individual reproduces by making copies of itself, while an Adult group or community can reproduce by randomly splitting into multiple Newborns of the next selection cycle. Successful artificial selection requires that i) entities display trait variations; ii) trait variations can be selected to result in differential entity survival and reproduction; and iii) entity trait is sufficiently heritable from one selection cycle to the next [104]. In all three types of selection, entity variations can be introduced by mutations and recombinations in individuals. However, heredity can be low in community selection. (**A**) Artificial selection of individuals has been successful [18, 19, 20, 105, 106], since a trait is largely heritable so long as mutation and recombination are sufficiently rare. (**B, C**) In group and community selection, if *T* is small so that newly-arising genotypes cannot rise to high frequencies within a selection cycle, then Adult trait is mostly determined by Newborn *composition* (the biomass of each genotype in each member species). Then, *variation* can be defined as the dissimilarity in Newborn composition within a selection cycle, and *heredity* as the similarity of Newborn composition from one cycle to the next for Newborns connected through lineage (tubes with same-colored outlines in Figure 4A). (**B)** Artificial selection of mono-species groups has been successful [43, 45, 15]. Suppose cooperators but not cheaters pay a fitness cost to generate a product (shade). Artificial selection for groups producing high total product favors cooperator-dominated groups, although within a group, cheaters grow faster than cooperators. At a large Newborn population size (**top**), all Newborns will harbor similar fractions of cheaters, and thus inter-group variation will be small [62]. During maturation, cheater frequency will increase, thereby diminishing heredity. In contrast, when Newborn groups are initiated at a small size such as one individual (**bottom**), a Newborn group will comprise either a cooperator or a cheater, thereby ensuring variation. Furthermore, even if cheaters were to arise during maturation, a fraction of Newborns of the next cycle will by chance inherit a cooperator, thereby ensuring some level of heredity. Thus, group selection can work when Newborn size is small. (**C**) Artificial selection of multi-species communities may be hindered by insufficient heredity. During maturation, the relative abundance of genotypes and species can rapidly change due to ecological interactions and evolution, which compromises heredity. During community reproduction, stochastic fluctuations in Newborn composition further reduce heredity.

**Figure S2:**
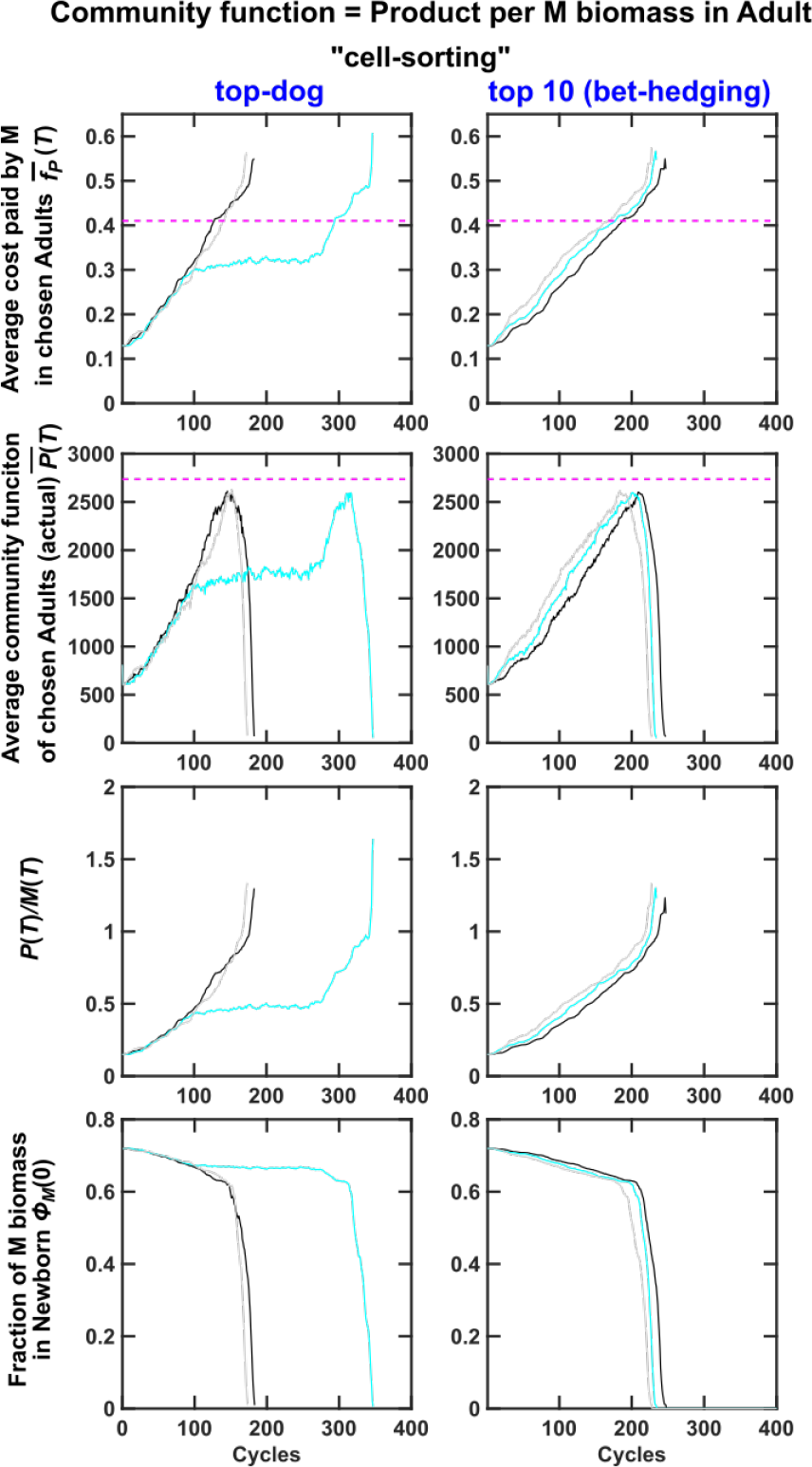
Problems of defining community function as *P*(*T*)*/M*(*T*). When the community function was defined by *P*(*T*)*/M*(*T*), average *f*_*P*_ of the chosen communities rapidly increased to such a high level that M was out-competed by H, as demonstrated by Figure 2A top panel. Consequently, selection abruptly came to a stop. Black, cyan and gray curves are three independent simulation trials. 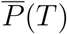 was averaged across chosen Adults. 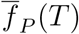 was obtained by first averaging among M within each chosen Adult, and then averaging across all chosen Adults.

**Figure S3:**
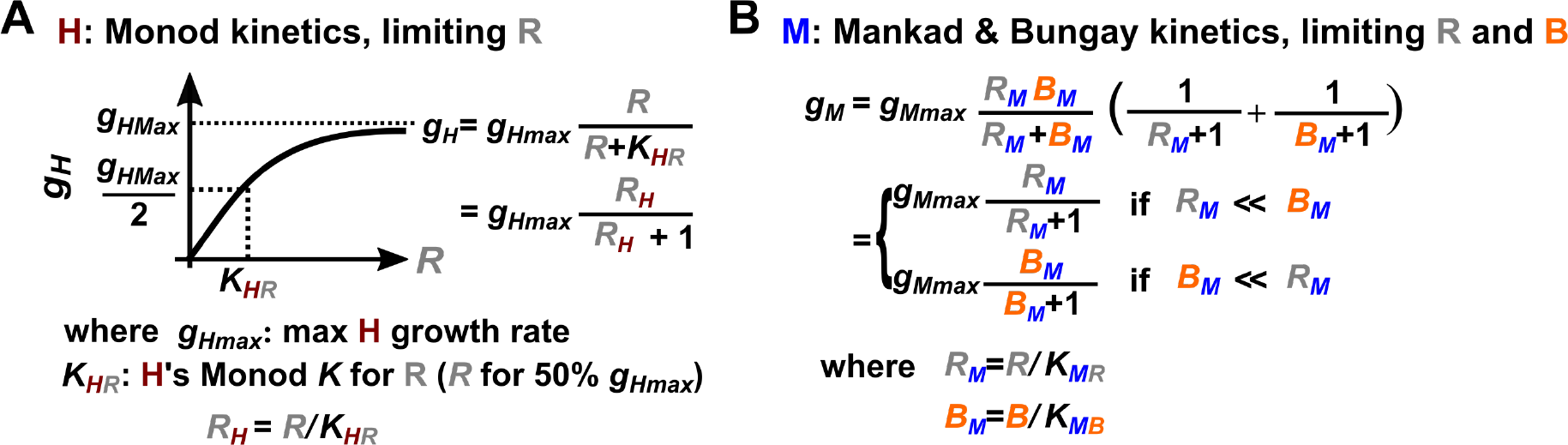
Growth models of H and M. (**A**) H growth follows Monod kinetics, reaching half maximal growth rate when *R* = *K*_*HR*_. (**B**) M growth follows dual-substrate Mankad-Bungay kinetics. When Resource *R* is in great excess (*R*_*M*_ ≫ *B*_*M*_) or Byproduct *B* is in great excess (*B*_*M*_ ≫ *R*_*M*_), we recover mono-substrate Monod kinetics (**A**).

**Figure S4:**
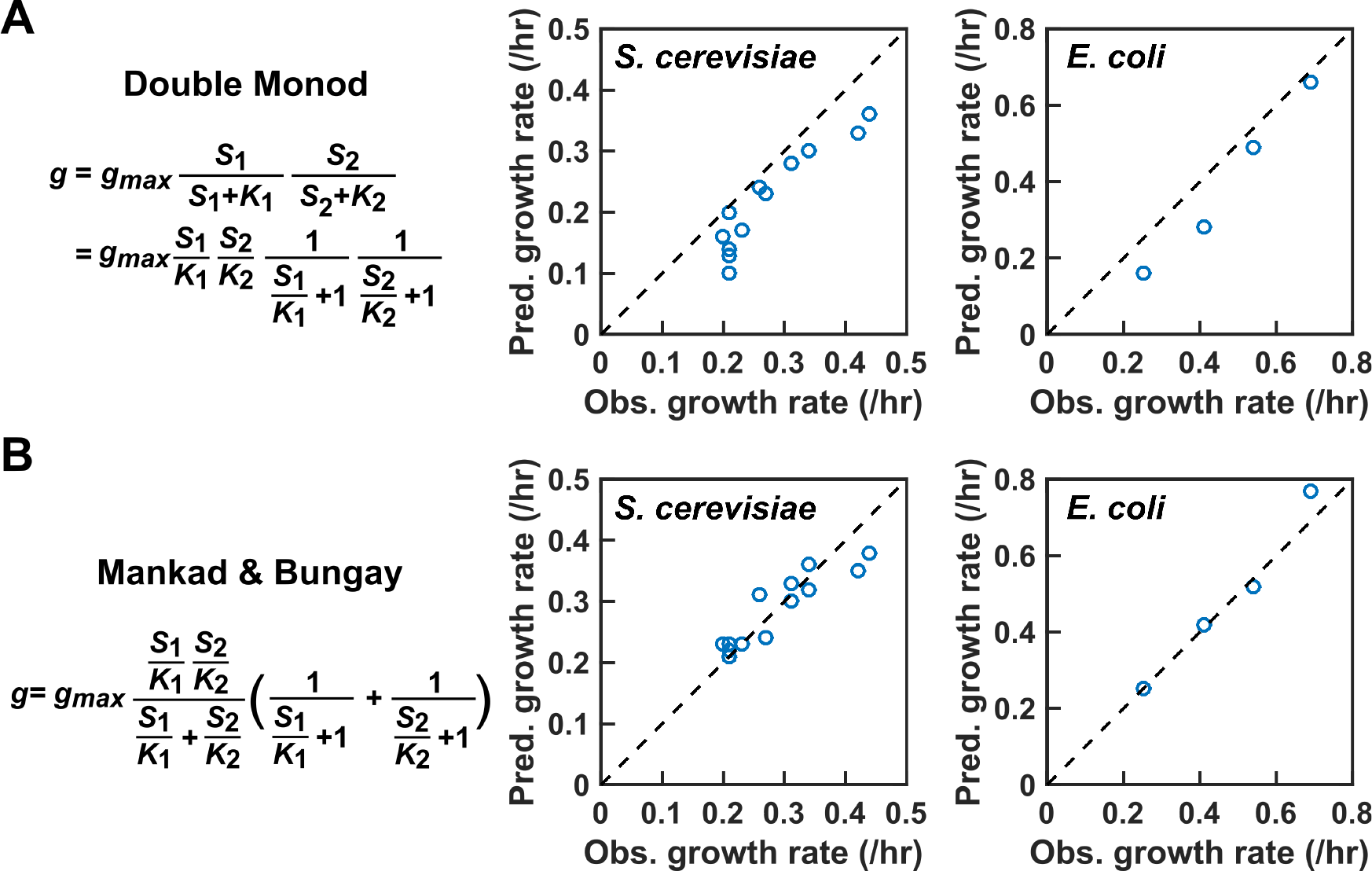
A comparison of dual-substrate models. Suppose that cell growth rate depends on each of the two substrates S_1_ and S_2_ in a Monod-like, saturable fashion. When S_2_ is in excess, the *S*_1_ at which half maximal growth rate is achieved is *K*_1_. When S_1_ is in excess, the *S*_2_ at which half maximal growth rate is achieved is *K*_2_. (**A**) In the “Double Monod” model, growth rate depends on the two limiting substrates in a multiplicative fashion. In the model proposed by Mankad and Bungay (**B**), growth rate takes a different form. In both models, when one substrate is in excess, growth rate depends on the other substrate in a Monod-fashion. However, when 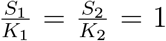, the growth rate is predicted to be *g*_*max*_/2 by Mankad & Bunday model, and *g*_*max*_/4 by the Double Monod model. Mankad and Bungay model outperforms the Double Monod model in describing experimental data of *S. cerevisiae* and *E. coli* growing on low glucose and low nitrogen. The figures are plotted using data from Ref. [27].

**Figure S5:**
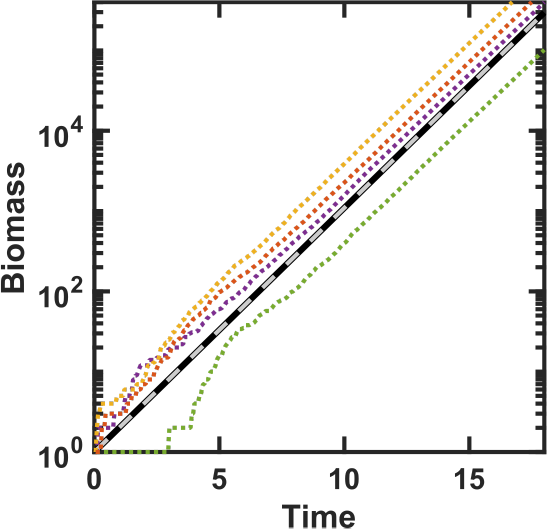
A comparison of different simulations of exponential cell growth in excess metabolites. Thick black line: analytical solution with biomass growth rate (0.7/time unit). Grey dashed line: simulation assuming that biomass increases exponentially at 0.7/time unit and that cell division occurs upon reaching a biomass threshold, an assumption used in our model. Color dotted lines: simulations assuming that cell birth is discrete and occurs at a probability equal to the birth rate multiplied with the length of simulation time step (Δ*τ* = 0.05 time unit). When a cell birth occurs, biomass increases discretely by 1, resulting in step-wise increase in color dotted lines at early time.

**Figure S6:**
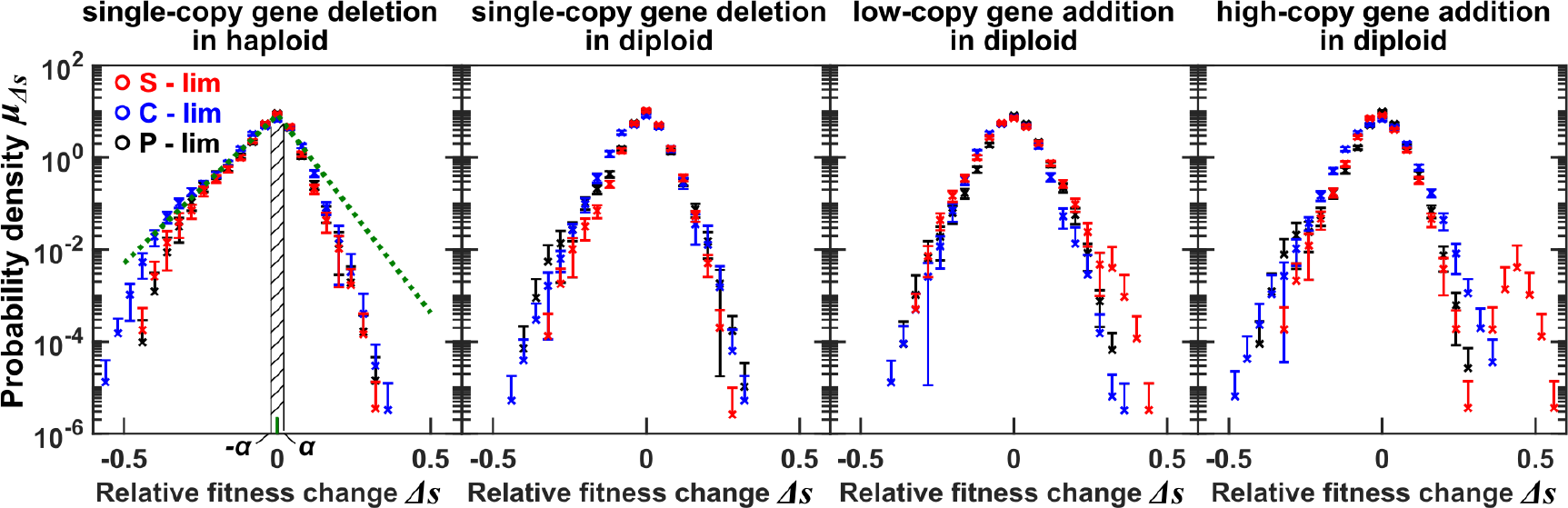
Probability density functions of changes in relative fitness due to mutations (*μ*_Δ*s*_(Δ*s*)). We derived *μ*_Δ*s*_(Δ*s*) from the Dunham lab data [36] where bar-coded mutant strains were competed under sulfate-limitation (red), carbon-limitation (blue), or phosphate-limitation (black). Error bars represent uncertainty *δμ*_Δ*s*_ (the lower error bar is omitted if the lower estimate is negative). In the leftmost panel, green lines show non-linear least squared fitting of data to Eq. 19 using all three sets of data. Note that data with larger uncertainty are given less weight, and thus deviate more from the fitting. For an exponentially-distributed probability density function *p*(*x*) = exp(−*x/r*)*/r* where *x, r* > 0, the average of *x* is *r*. When plotted on a semi-log scale, we get a straight line with slope 1*/r*, and inverting this gets us the average effect *r*. From the green line on the right side, we obtain the average effect of enhancing mutations *s*_+_ = 0.050 ± 0.002, and from the green line on the left side, we obtain the average effect of diminishing mutations *s*_−_ = 0.067 ± 0.003. The probability of a mutation altering a phenotype by ±*α* is the area of the hatched region drawn in the leftmost panel.

**Figure S7:**
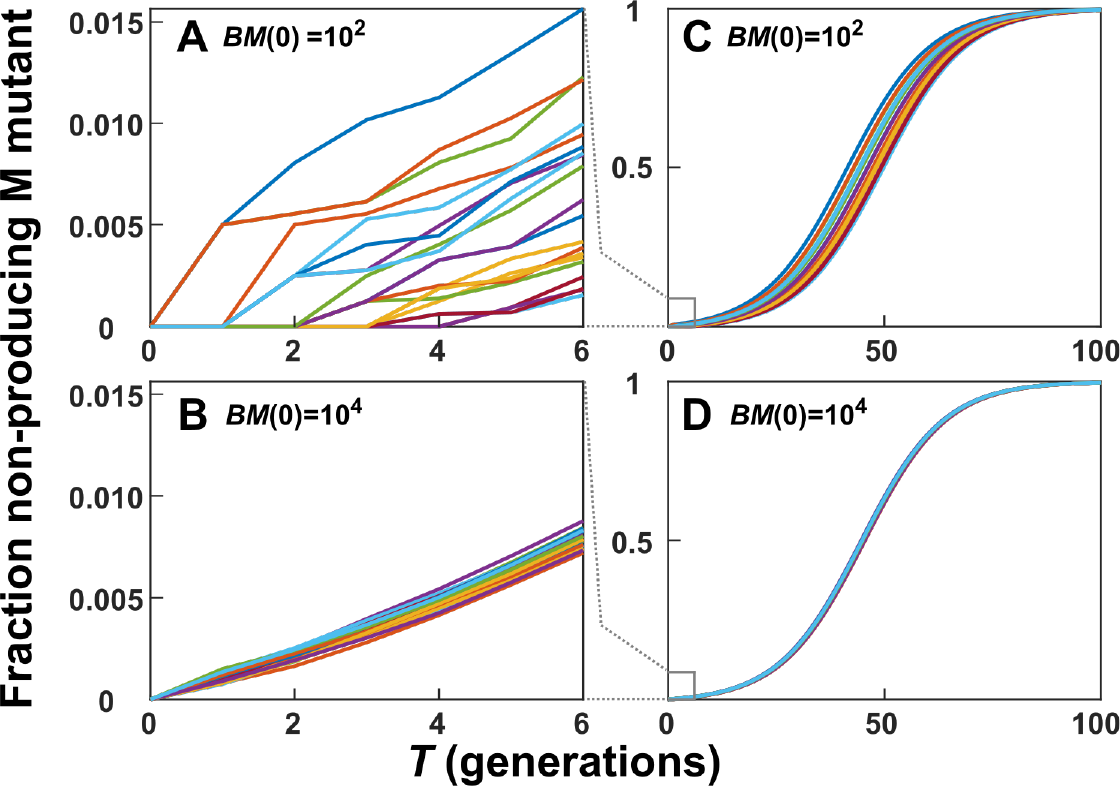
Large Newborn group size or long maturation time allows non-contributors to accumulate and reduces inter-group variation. For simplicity, we modeled the growth of Newborn groups of M cells. From a Newborn biomass of 10^2^ or 10^4^ wild-type M cells, M population multiplied for 6 or 100 generations. Immediately following cell division, wild-type daughter cells mutated to non-contributors with a probability of 10^−3^. Wild-type and mutant cells followed exponential growth. The growth rate of wild-type cells was 0.87 times that of mutants. The fraction of biomass made up by mutants at each wild-type doubling is shown. Note the different scales.

**Figure S8:**
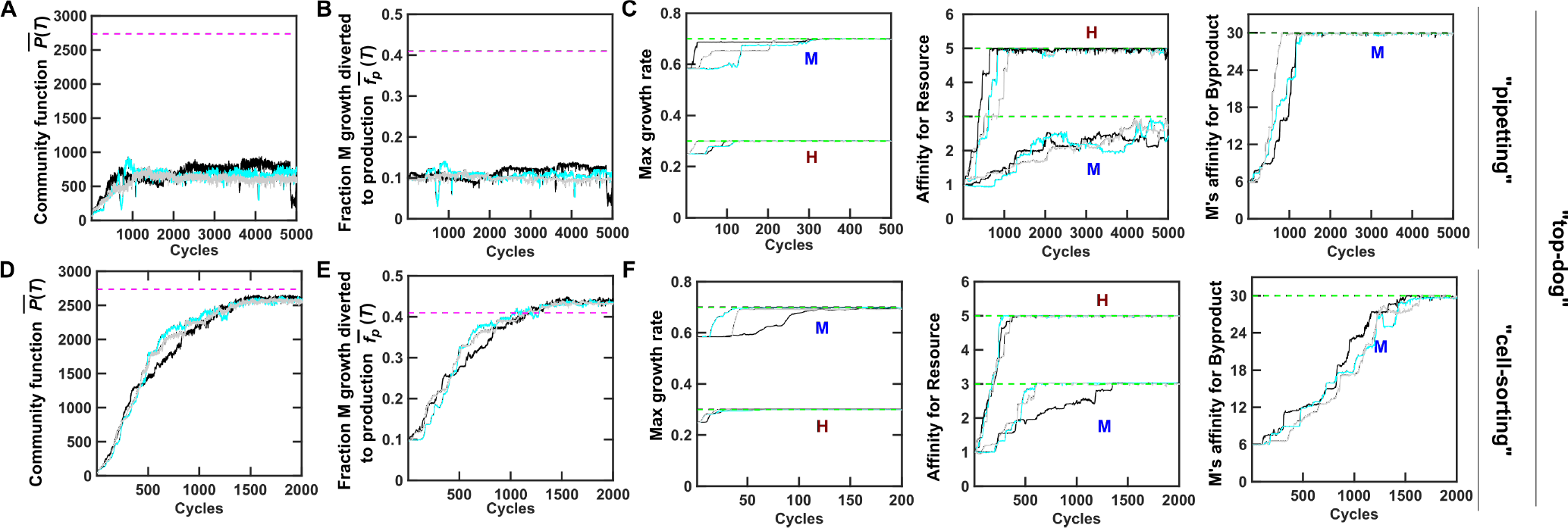
Improved individual growth can promote community function. Here, we allowed mutations to alter M’s *f*_*P*_ and H and M’s growth parameters. Communities are chosen using the “top-dog” strategy. (**A-C**) Community reproduction via pipetting (i.e. Newborn biomass and species composition can fluctuate). Community function *P*(*T*) increased upon community selection (**A**). Since *f*_*P*_ remained unchanged (**B**), this increase in *P*(*T*) must be due to improved growth parameters (**C**). (**D-F**) Community reproduction via biomass sorting (i.e. fixed Newborn total biomass and species composition). In both cases, the five growth parameters increased to their respective evolutionary upper bounds (green dashed lines). Magenta dashed lines: *f*_*P*_ optimal for community function and maximal community function *P*(*T*) when all five growth parameters are fixed at their evolutionary upper bounds and *ϕ*_*M*_(0) is also optimal for *P*(*T*). Black, cyan, and gray curves show three independent simulations. 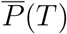 is averaged across chosen Adults. 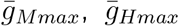, and 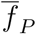 are obtained by averaging within each chosen Adult and then averaging across chosen Adults. *K*_*SpeciesMetabolite*_ are averaged within each chosen Adult, then averaged across chosen Adults, and finally inverted to represent average affinity. Note different *x* axis scales. The maximal growth rates (*g*_*Mmax*_ and *g*_*Hmax*_) have the unit of 1/time. Affinity for Resource (1*/K*_*MR*_, 1*/K*_*HR*_) has the unit of 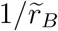, where 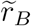 is the initial amount of Resource in Newborn. Affinity for Byproduct (1/*K*_*MB*_) has the unit of 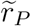, the amount of Product released at the cost of one M biomass. More details on parameters can be found in Table 1.

**Figure S9:**
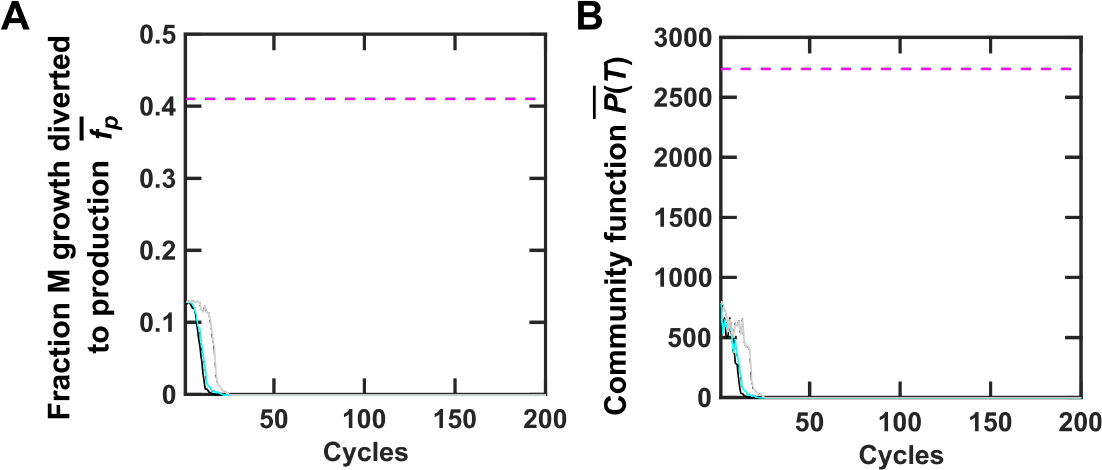
Community function declines to zero in the absence of inter-community selection for higher community function. When random Adults were chosen to reproduce (“pipetting”), natural selection favored zero *f*_*P*_ (**A**). Consequently, *P*(*T*) decreased to zero (**B**). Black, cyan and gray curves are three independent simulation trials. 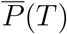 was averaged across the two randomly chosen Adults. 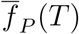 was obtained by first averaging among M within each randomly chosen Adult and then averaging across the two randomly chosen Adults.

**Figure S10:**
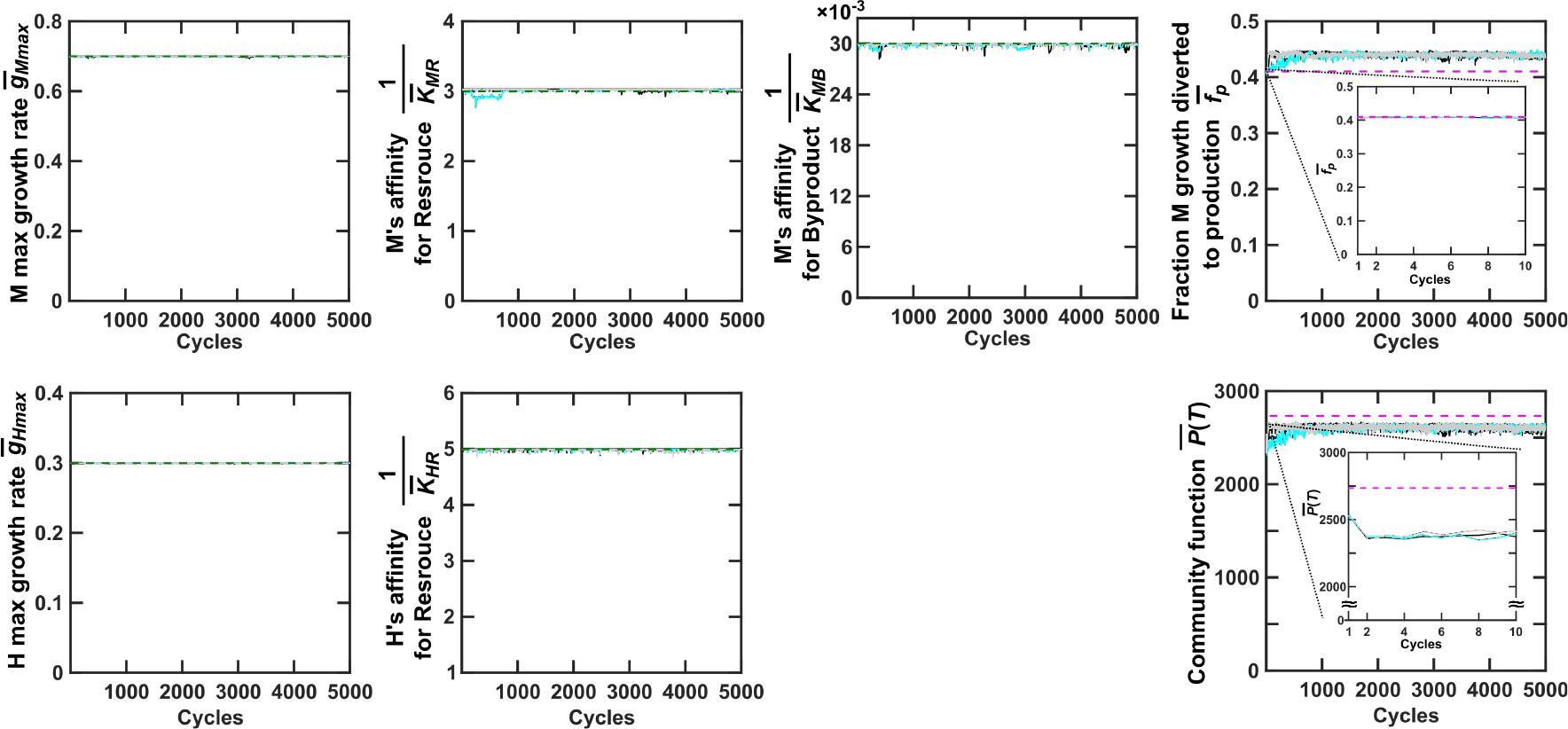
*P*\*(*T*) is a local optimum because it cannot be further improved. We started each Newborn community with total biomass *BM*(0) = 100, all five growth parameters at their evolutionary upper bounds, and 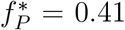 and 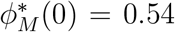 to achieve *P*\*(*T*). We then allowed all five growth parameters and *f*_*P*_ to mutate while applying community selection. To ensure effective community selection (Figure 3D-F), *BM*(0) was fixed to 100, and *ϕ*_*M*_(0) was fixed to *ϕ*_*M*_ (*T*) of the chosen Adult community from the previous cycle during community reproduction. We found that all five growth parameters remained at their respective evolutionary upper bounds. At the end of the first cycle (Cycle = 1 in insets), even though 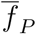 did not change, 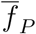 had already declined from the original magenta dashed line. This is because species interactions have driven *ϕ*_*M*_(0) from the optimal 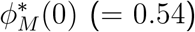 to near the steady state value (*ϕ*_*M*_ = 0.72, compare with *ϕ*_*M,SS*_ represented by the green dashed line in Figure 2A bottom panel). Later, over hundreds of cycles, 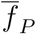 gradually increased while 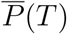 was still below maximal. This is because species composition gravitated toward steady state *ϕ*_*M,SS*_ which deviated from the optimal 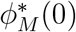 ([64]). Other legends are the same as Figure S8.

**Figure S11:**
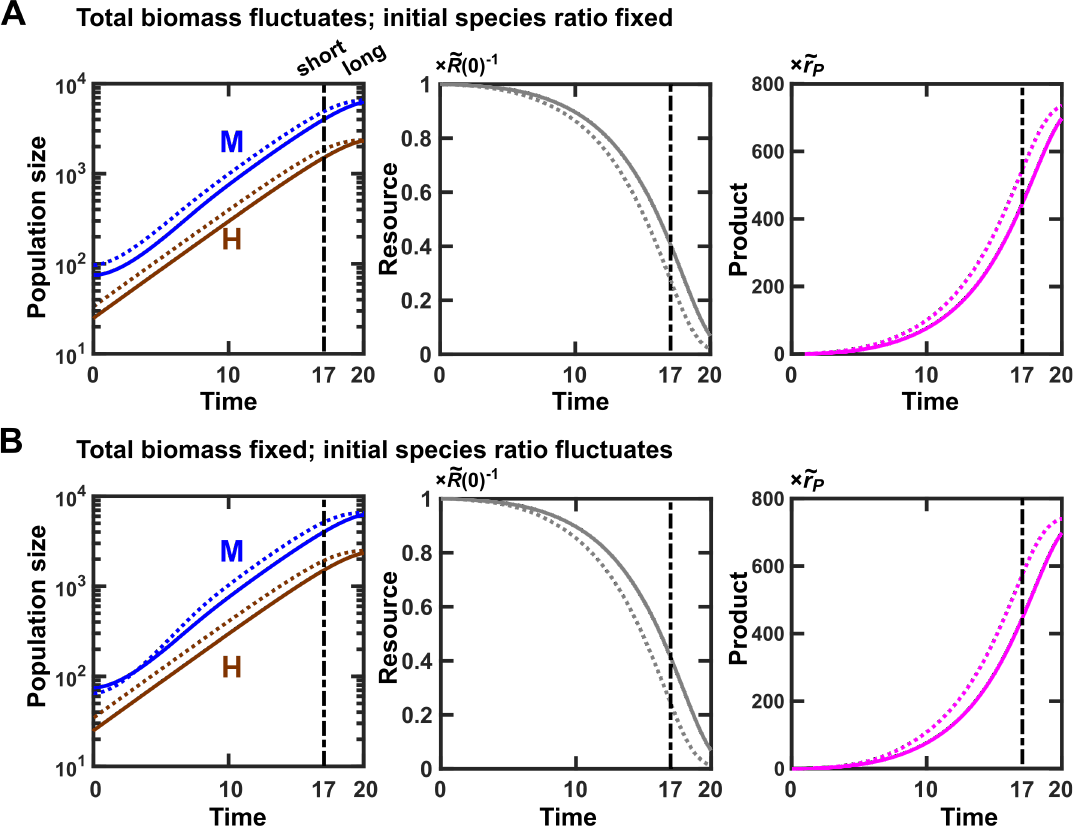
Variations in community function can arise from non-heritable variations in New-born compositions. An average Newborn community (solid lines) has a total biomass of 100 with 75% M. (**A**) A “lucky” Newborn community (dotted lines), by stochastic fluctuations, has a total biomass of 130 with 75% M. Even though the two communities share identical *f*_*P*_ = 0.1, biomass of M in the Newborn starting with a total biomass of 130 can grow to a higher value (left), deplete more Resource (middle), and make more Product (right) by the end of short *T* (*T* = 17). (**B**) A “lucky” Newborn community (dotted lines), by stochastic fluctuations, has a total biomass of 100 with 65% (instead of 75%) M. Even though the two communities share identical *f*_*P*_ = 0.1, higher fraction of Helper H biomass results in faster accumulation of Byproduct. Consequently, M (dotted) can enjoy a shorter growth lag, grow to a larger size (left), deplete more Resource (middle), and make more Product (right) by the end of short *T* (*T* = 17). In both cases, the difference between lucky (dotted) and average (solid) communities is diminished at longer *T* (*T* = 20) compared to shorter *T* (*T* = 17, dash dot line).

**Figure S12:**
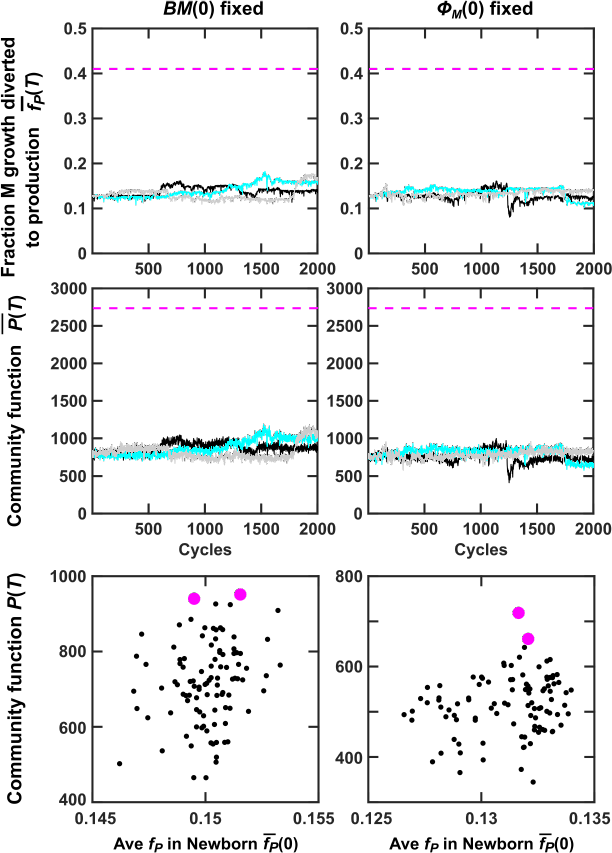
Fixing either total biomass *BM*(0) or fraction of M biomass *ϕ*_*M*_(0) in Newborns did not significantly improve community selection when using the “top-dog” strategy. Black, cyan and gray curves are three independent simulation trials. 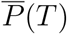 was averaged across the two chosen Adults. 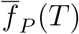 was obtained by first averaging among M within each chosen Adult and then averaging across the two chosen Adults.

**Figure S13:**
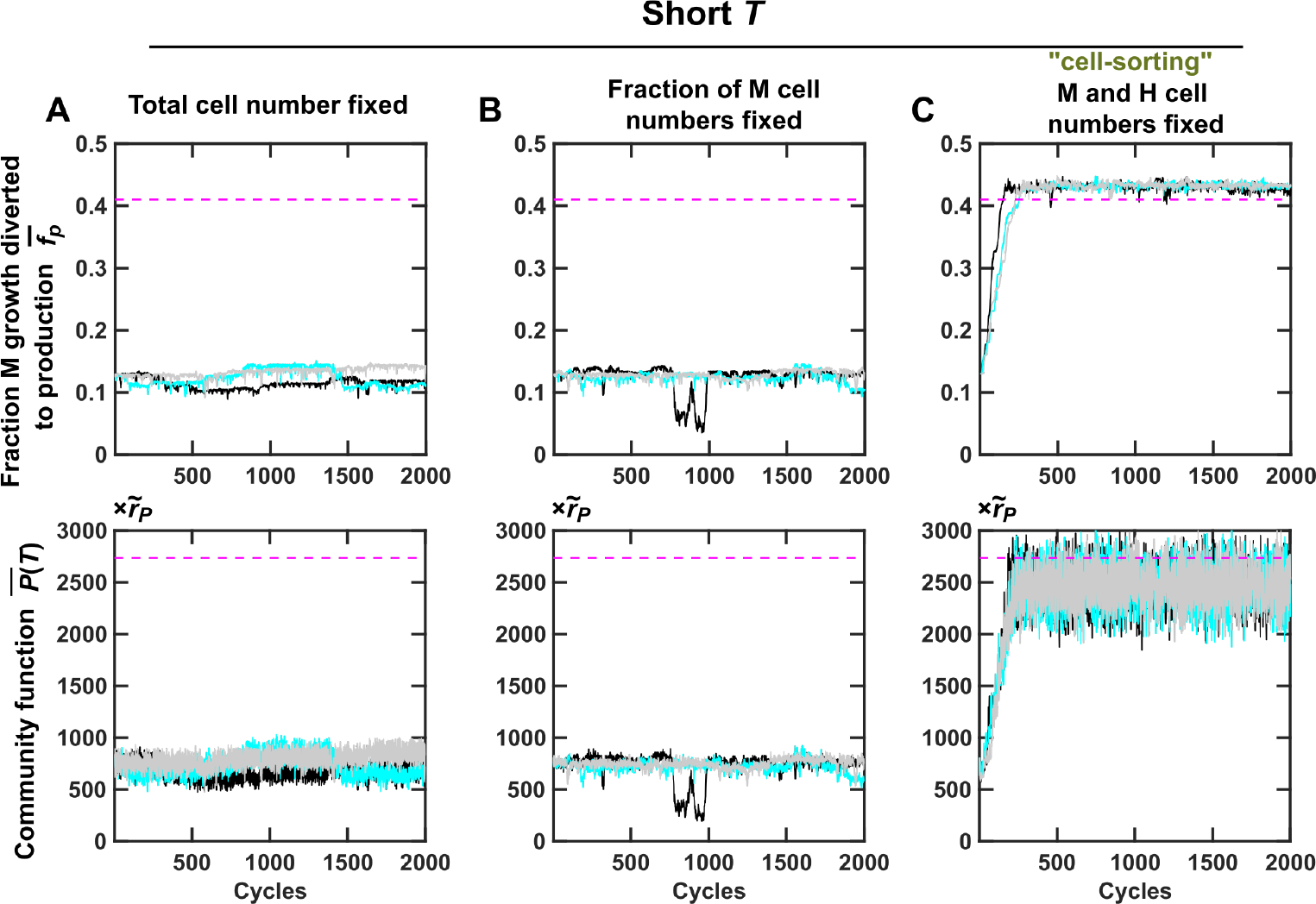
Fixing H and M cell numbers (instead of biomass) during community reproduction allows short-*T* selection regimen to improve community function under the “top-dog” strategy. (**A**) the total cell number in Newborn communities was fixed to ⌊*BM*_*target*_/1.5⌋ where ⌊*x*⌋ means rounding down *x* to the nearest integer. (**B**) The ratio between M and H cell numbers in Newborn communities were fixed to *I*_*M*_ (*T*)*/I*_*H*_ (*T*), where *I*_*M*_ (*T*) and *I*_*H*_ (*T*) were the number of M and H cells in the chosen Adult community from the previous cycle, respectively. (**C**) The total cell numbers of Newborn communities were fixed to ⌊*BM*_*target*_/1.5⌋ and the ratio between M and H cell numbers were fixed to *I*_*M*_ (*T*)*/I*_*H*_ (*T*). See Methods Section 6 for details of simulating community reproduction. Black, cyan and gray curves are three independent simulation trials. 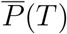 was averaged across the two chosen Adults. 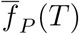 was obtained by first averaging among M within each chosen Adult and then averaging across the two chosen Adults.

**Figure S14:**
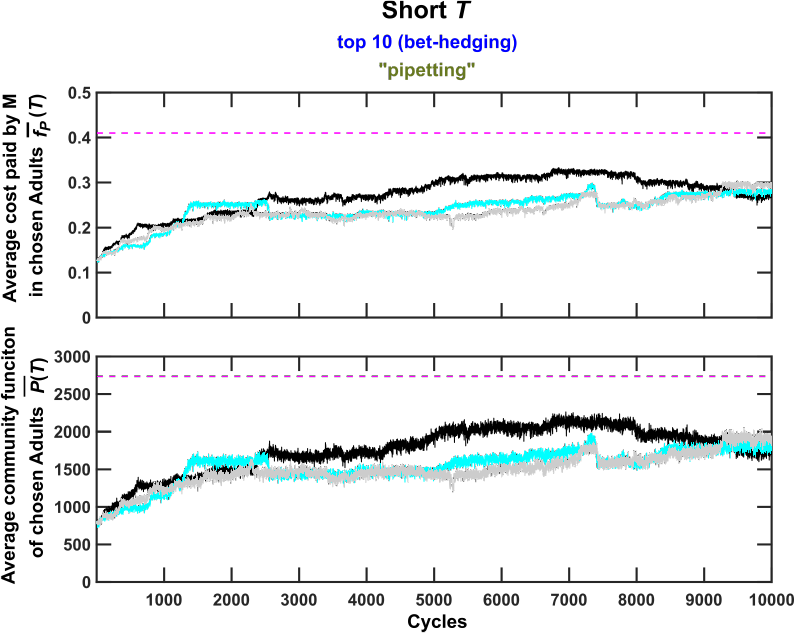
Bet-hedging can improve selection efficacy when community function experiences non-heriable variations, but not as effectively as reducing non-heritable variations in community function. Simulation setup is identical to that in Figure 3G-H, except that selection here lasts more cycles. Compared to Figure 3A-C, bet-hedging was more effective. However, compared to Figure 3D-F, bet-hedging did not improve community function to the same extent, even over 10^4^ cycles. Black, cyan and gray curves are three independent simulation trials. 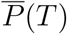 was averaged across the chosen Adults. 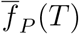 was obtained by first averaging among M within each chosen Adult and then averaging across all chosen Adults.

**Figure S15:**
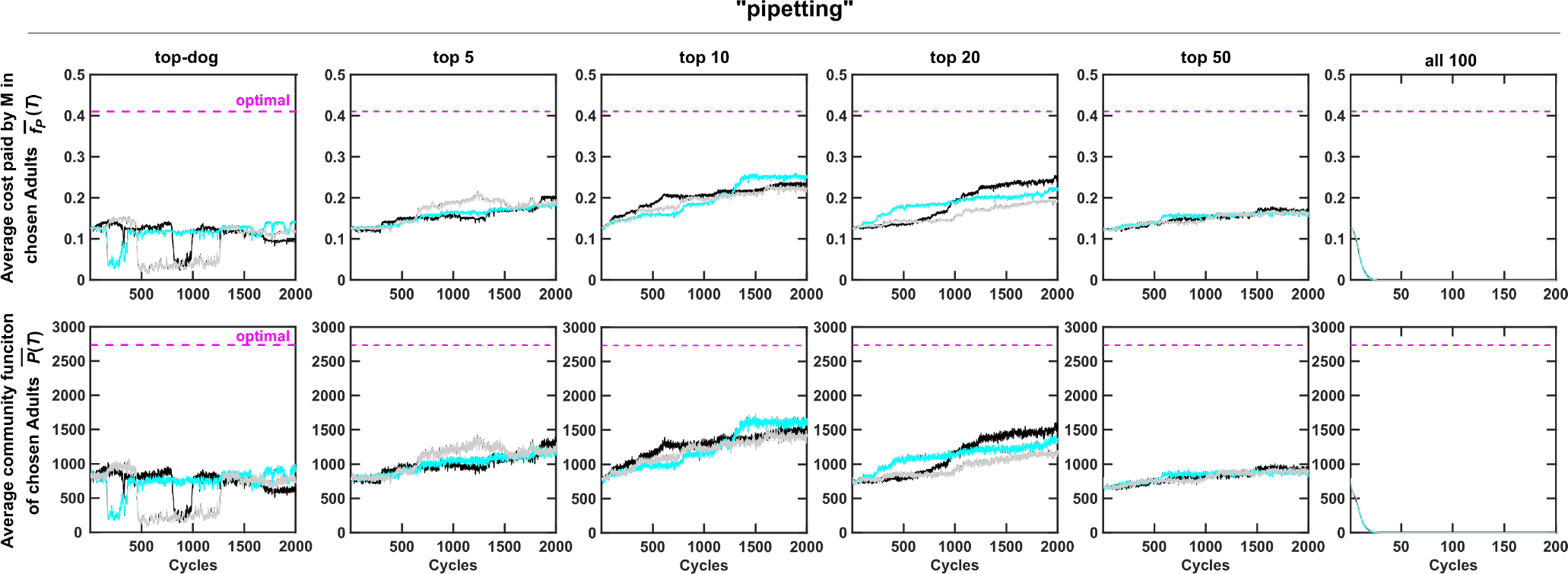
Bet-hedging strategies promoted community selection under a wide range of selection strengths. In a bet-hedging strategy, top *k* Adults each contributed 100*/k* Newborns into the next cycle. Here, Adults were reproduced (split) into Newborns via pipetting. When all Adults contributed one Newborn each, selection strength was zero and thus natural selection quickly reduced average *f*_*P*_ and community function to zero (rightmost column). Black, cyan and gray curves are three independent simulation trials. 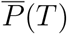 was averaged across the two chosen Adults. 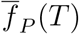 was obtained by first averaging among M within each chosen Adult and then averaging across all chosen Adults.

**Figure S16:**
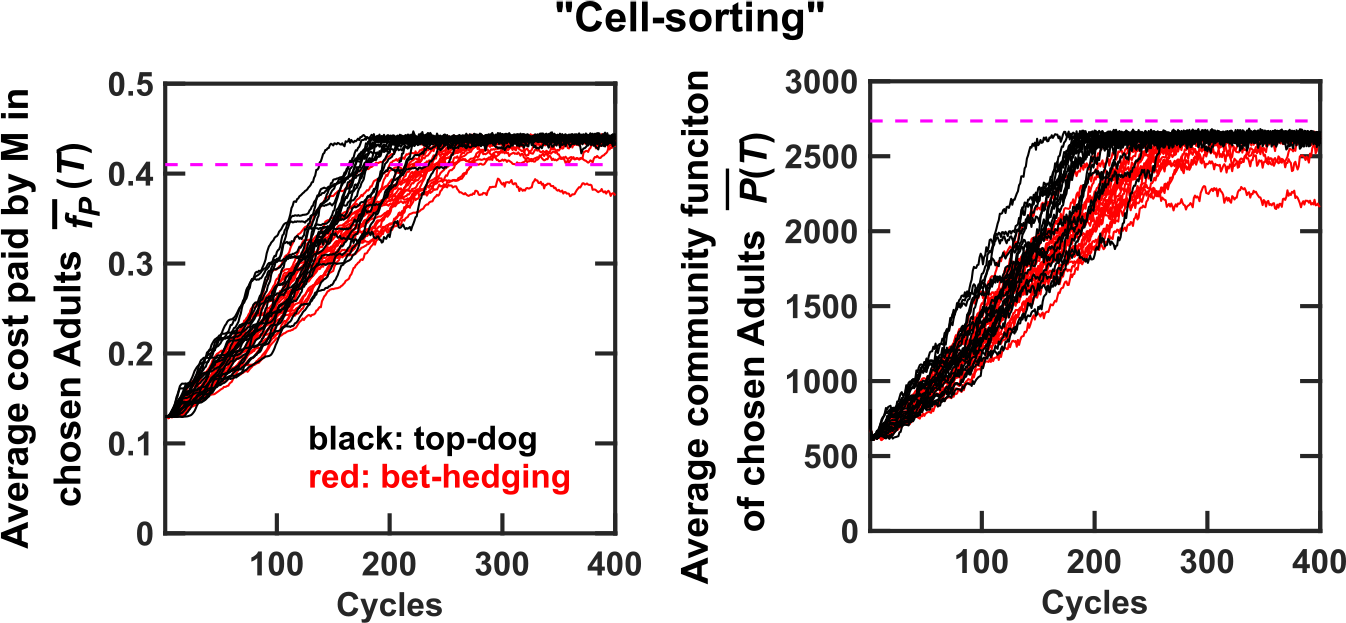
The “top-dog” strategy is superior to the “bet-hedging” strategy when non-heritable variation in community function is low.. 20 replicas of selection simulations were performed using either the “top-dog” strategy (black curves) or the “bet-hedging” strategy (top ten Adults chosen to reproduce; red curves). Community reproduction was through cell-sorting. Community functions improved slightly faster and to a slightly higher level using the “top-dog” strategy. Thus, when non-heritable variations in community function were suppressed, the “top-dog” strategy was superior to the “bet-hedging” strategy.

**Figure S17:**
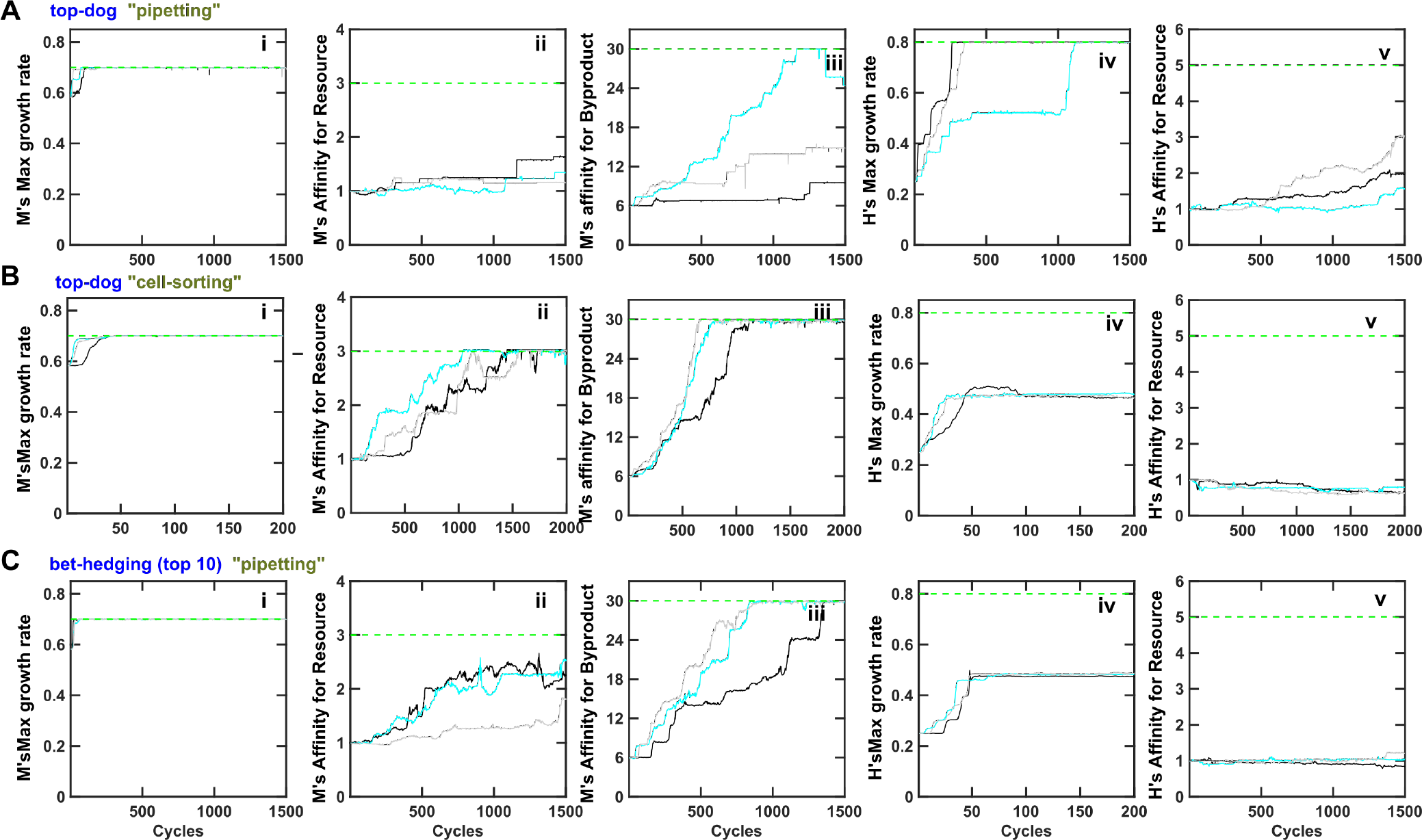
Community function can be improved even if it is costly to both species. Identical to Figure 6, the evolutionary upper bound for 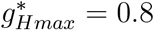 was larger than that of *g*_*Mmax*_ 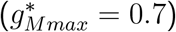, opposite to that in Figure 3. (**A**) Chosen Adult communities were reproduced through pipetting such that both *BM*(0) and *ϕ*_*M*_(0) could stochastically fluctuate. Eventually, *g*_*Hmax*_ and *g*_*Mmax*_ evolved to their respective upper bounds, and thus *g*_*Hmax*_ > *g*_*Mmax*_ (compare **i** and **iv**). This would ordinarily lead to extinction of M. However, community selection managed to maintain M at a very low level (Figure 6A bottom panel). (**B**, **C**) Adult communities were chosen using the top-dog strategy and reproduced through cell biomass sorting (**B**) or chosen using the bet-hedging strategy and reproduced through pipetting (**C**). Community selection worked in the sense that both 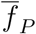 and *P*(*T*) improved over cycles (Figure 6). Strikingly, the maximal growth rate of H *g*_*Hmax*_ did not increase to its upper bound 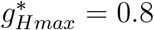, and H’s affinity for Resource even decreased from the ancestral level in some cases. Here, Resource supplied to Newborn communities could support 10^5^ total biomass to accommodate faster growth rate. Other legends are the same as Figure S8.

**Figure S18:**
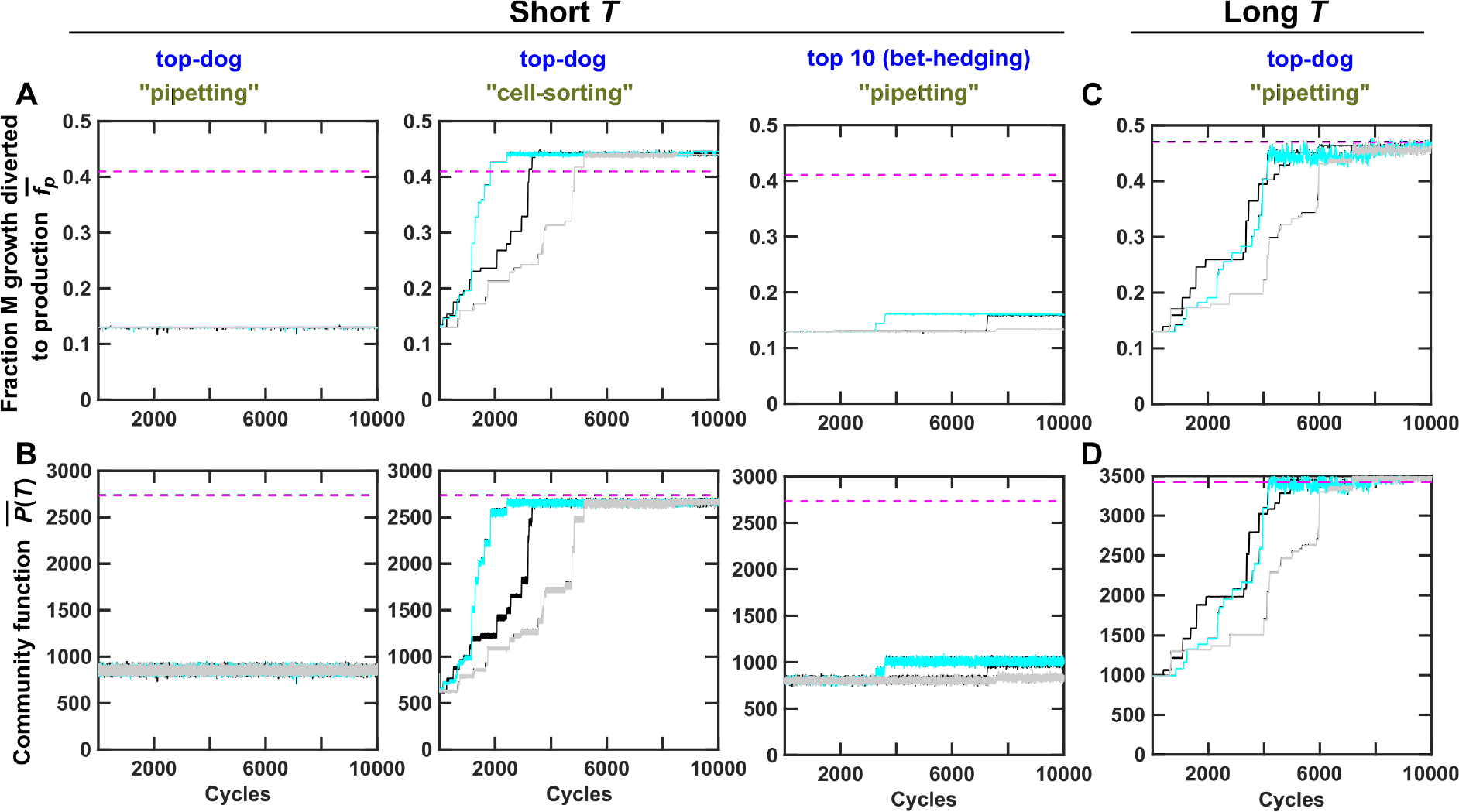
Evolution dynamics of chosen Adult communities at a mutation rate of 2 × 10^−5^ per cell per generation. (**A, B**) At short maturation time (*T* = 17, Resource was not exhausted in an average community), fixing both *BM*(0) and *ϕ*_*M*_(0) (“cell-sorting”) improved community function. The bet-hedging strategy slightly improved community function. (**C, D**) At long maturation time (*T* = 20, Resource was nearly exhausted in an average community), community function improved without fixing *BM*(0) or *ϕ*_*M*_(0). At this mutation rate, because the population size of a community never exceeds 10^4^, a mutation occurs on average every 5 cycles, resulting in step-wise improvement in both 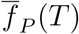 and 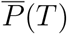. Black, cyan and gray curves are three independent simulation trials. 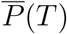 was averaged across all chosen Adults. 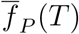 was obtained by first averaging among M within each chosen Adult and then averaging across all chosen Adults.

**Figure S19:**
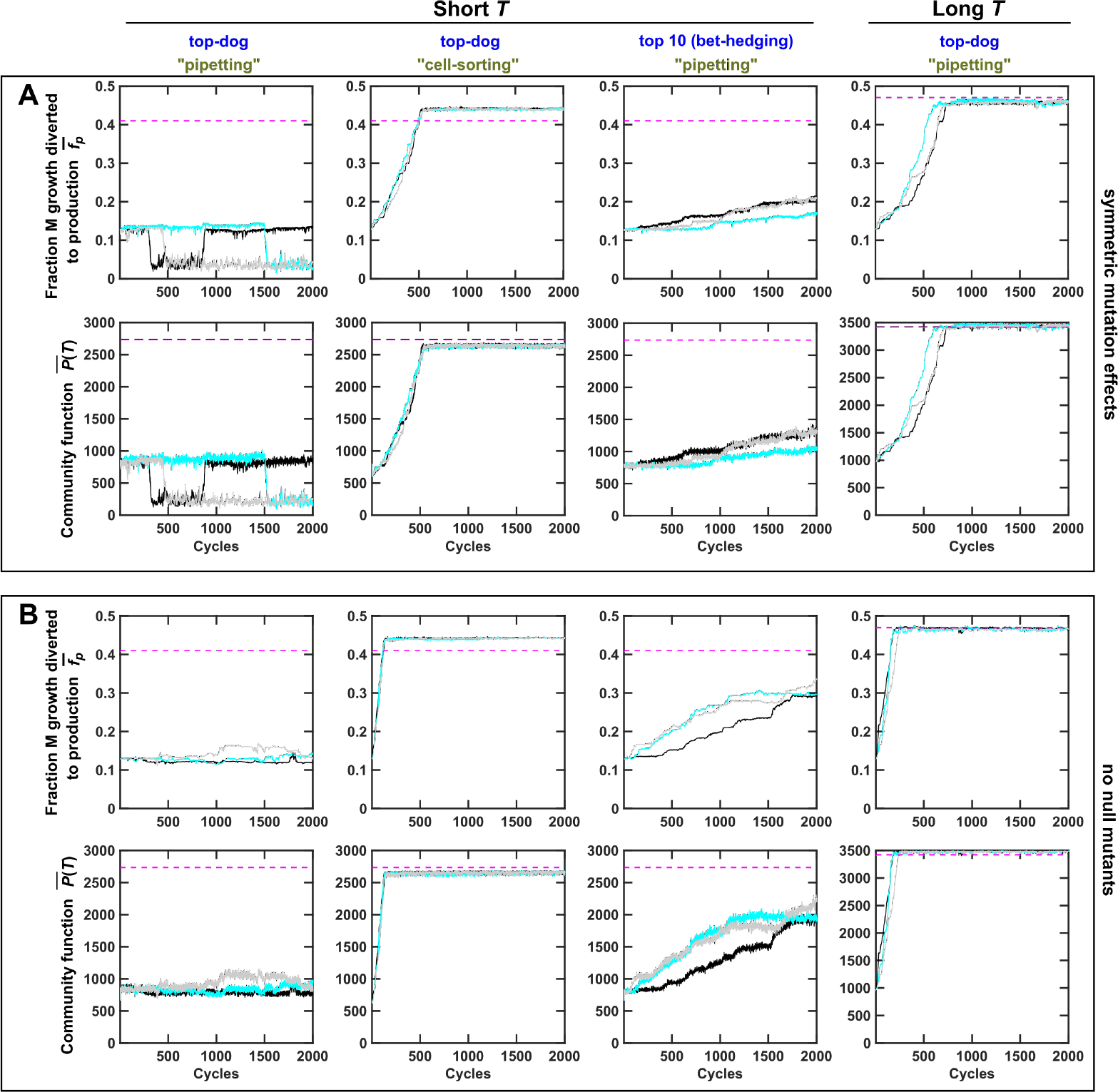
Evolutionary dynamics of chosen Adult communities under different distributions of mutation effects. (A) Evolutionary dynamics where half of the mutations reduced *f*_*P*_ to zero and the distribution of mutation effects of the other half is specified by Eq. 19 where *s*_+_ = *s*_−_ = 0.02 are constants. (B) Evolutionary dynamics when null mutations in *f*_*P*_ did not occur. The distribution of mutation effects is thus similar to that used in Figure 3, except that the null mutations were eliminated. Specifically, the distribution of mutation effects is specified by Eq. 19 where *s*_+_ = 0.05 and *s*_−_ = 0.067. 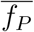 as well as *P*(*T*) were more stable compared to those in Figure 3. Black, cyan and gray curves are three independent simulation trials. 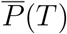 was averaged across the chosen Adults. 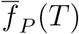 was obtained by first averaging among M within each chosen Adult and then averaging across all chosen Adults.

**Figure S20:**
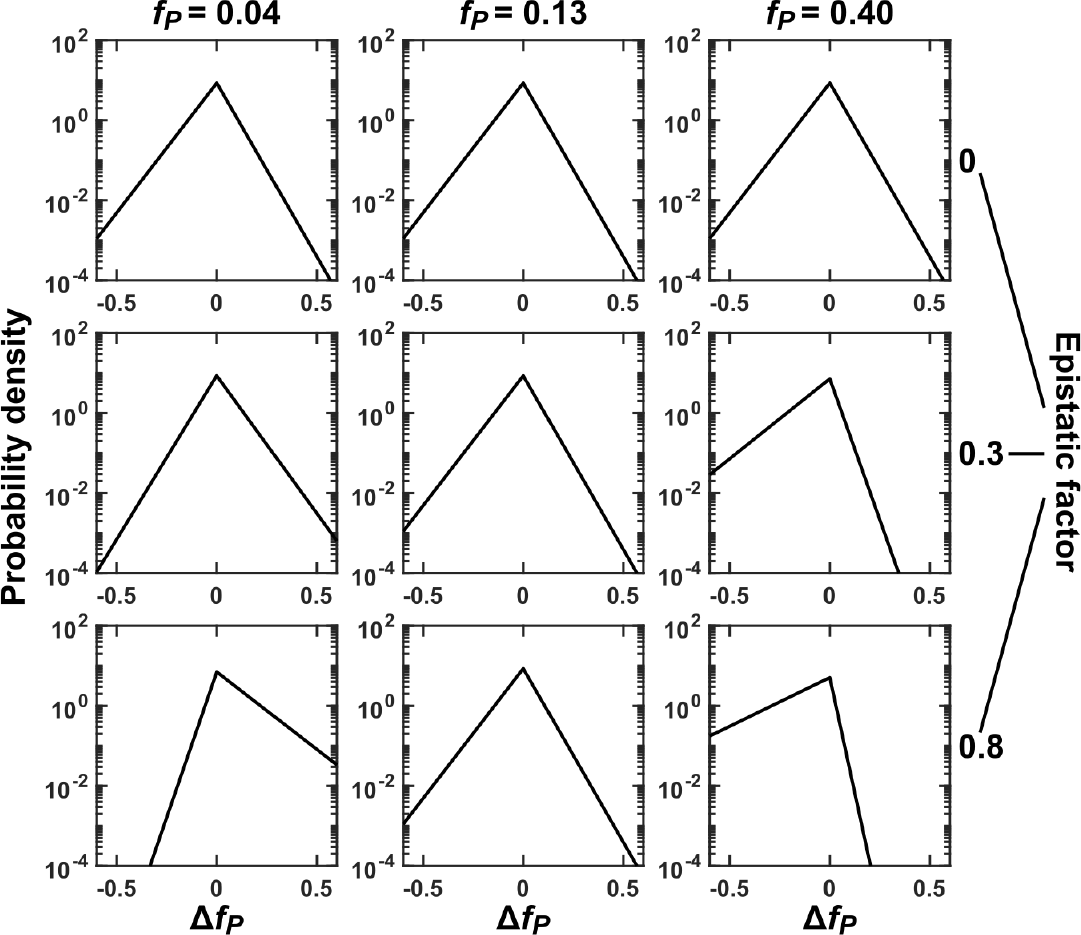
Mutation effects under epistasis. Distribution of mutation effects at different current *f*_*P*_ values (marked on top) are plotted. (Top) When there is no epistasis, distribution of mutational effects on *f*_*P*_ (Δ*f*_*P*_) are identical regardless of current *f*_*P*_. (Middle and Bottom) With epistasis (see Methods Section 5 for definition of epistasis factor), mutational effects on *f*_*P*_ depend on the current value of *f*_*P*_. If current *f*_*P*_ is low (left), enhancing mutations are more likely to occur (the area to the right of Δ*f*_*P*_ = 0 becomes bigger) and their mean mutational effect becomes larger (mean=1/slope becomes larger due to smaller slope), while diminishing mutations are less likely to occur and their mean mutational effect is smaller. If current *f*_*P*_ is high (right), the opposite is true.

**Figure S21:**
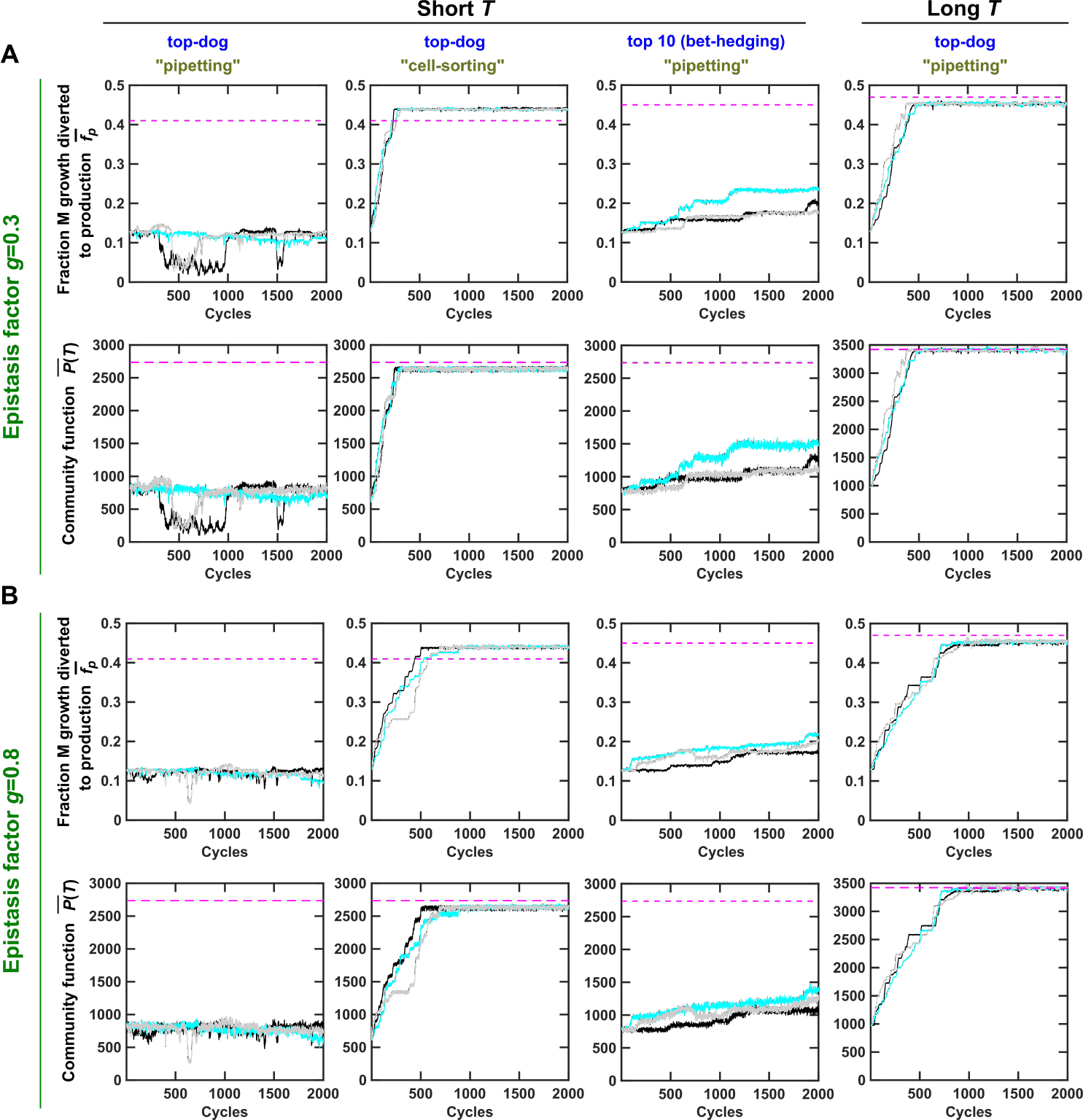
Evolutionary dynamics of chosen Adults when epistasis is considered. When we incorporated different epistasis strengths (epistasis factor of 0.3 and 0.8), we obtained essentially the same conclusions as when epistasis was not considered (Figure 3). Other legend details can be found in Figure 3.

**Figure S22:**
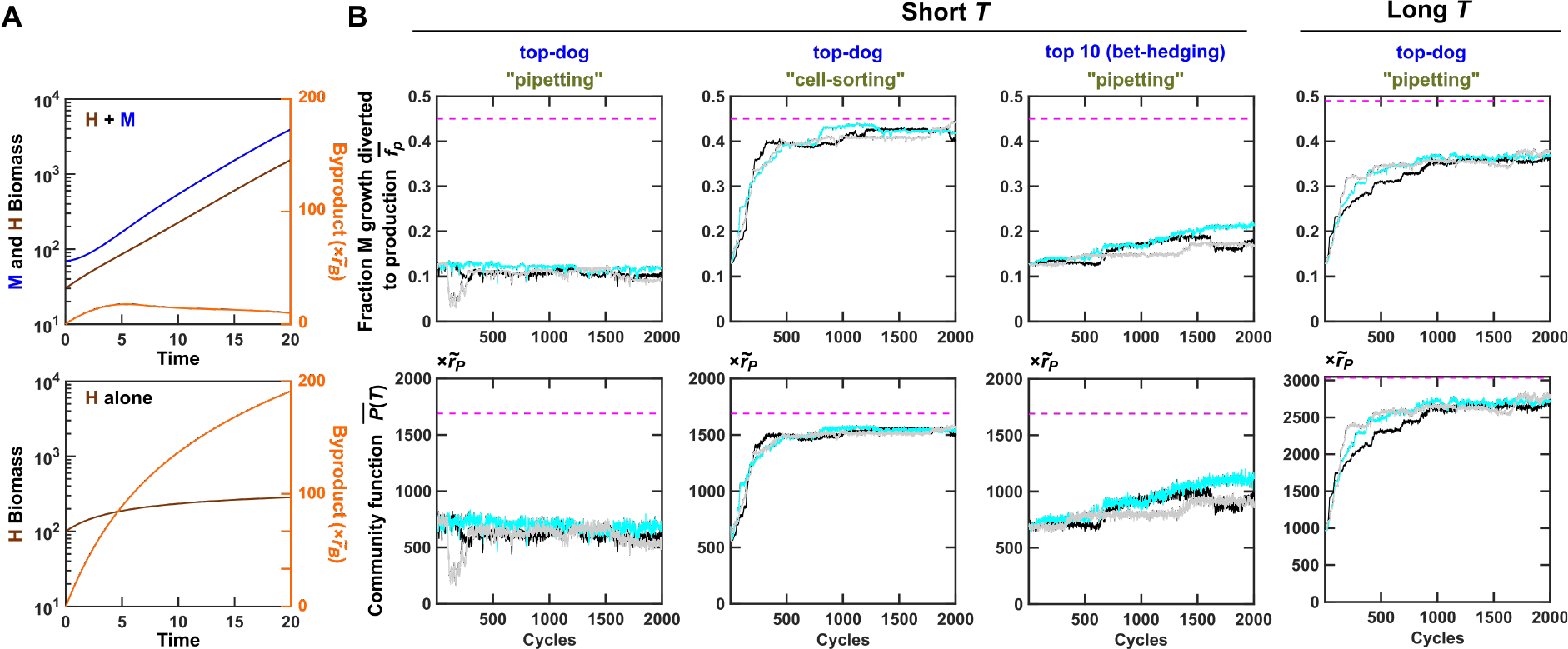
Selection dynamics of mutualistic H-M communities. In the mutualistic H-M community, H generates Byproduct which is essential for M but inhibitory to H. (**A**) H can grow to a high density in the presence of M (top) but not in the absence of M (bottom). (**B**) Similar to community selection on commensal H-M communities, selection was promoted by bet-hedging or cell-sorting at short *T* (*T* = 17), or via extending *T* (*T* = 20). Black, cyan and gray curves are three independent simulation trials. 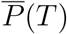 was averaged across the chosen Adults. 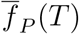 was obtained by first averaging among M within each chosen Adult and then averaging across all chosen Adults.

**Figure S23:**
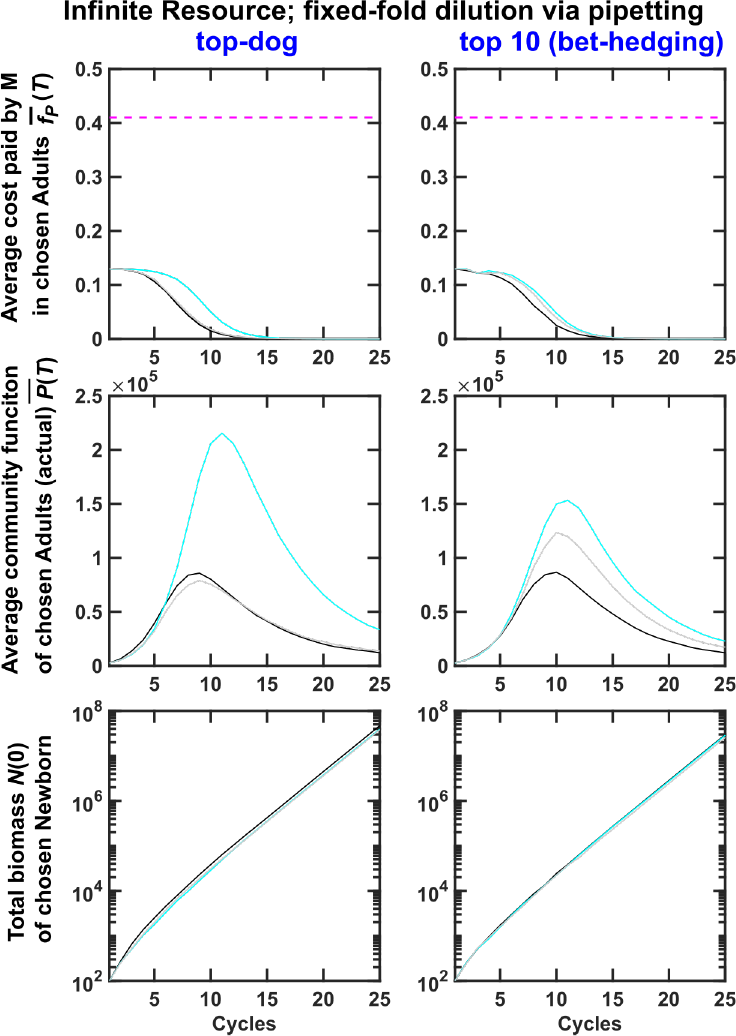
Artificial selection on community function in excess Resource failed under fixed-fold pipetting dilution scheme. Excess Resource was supplied to each Newborn (*R*(0)*/K*_*MR*_ = 10^6^), and chosen Adults were reproduced via a fixed-fold (100-fold) pipetting dilution into Newborns. Because of pipetting, Newborns with larger total biomass will tend to be selected (Figure 4F). Community selection quickly failed as Newborn total biomass increased exponentially (bottom) while non-producing M cells with *f*_*P*_ = 0 quickly took over (top; Figure S7B). Black, cyan and gray curves are three independent simulation trials. 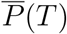 was averaged across chosen Adults. 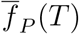 was obtained by first averaging among M within each chosen Adult and then averaging across all chosen Adults.

**Figure S24:**
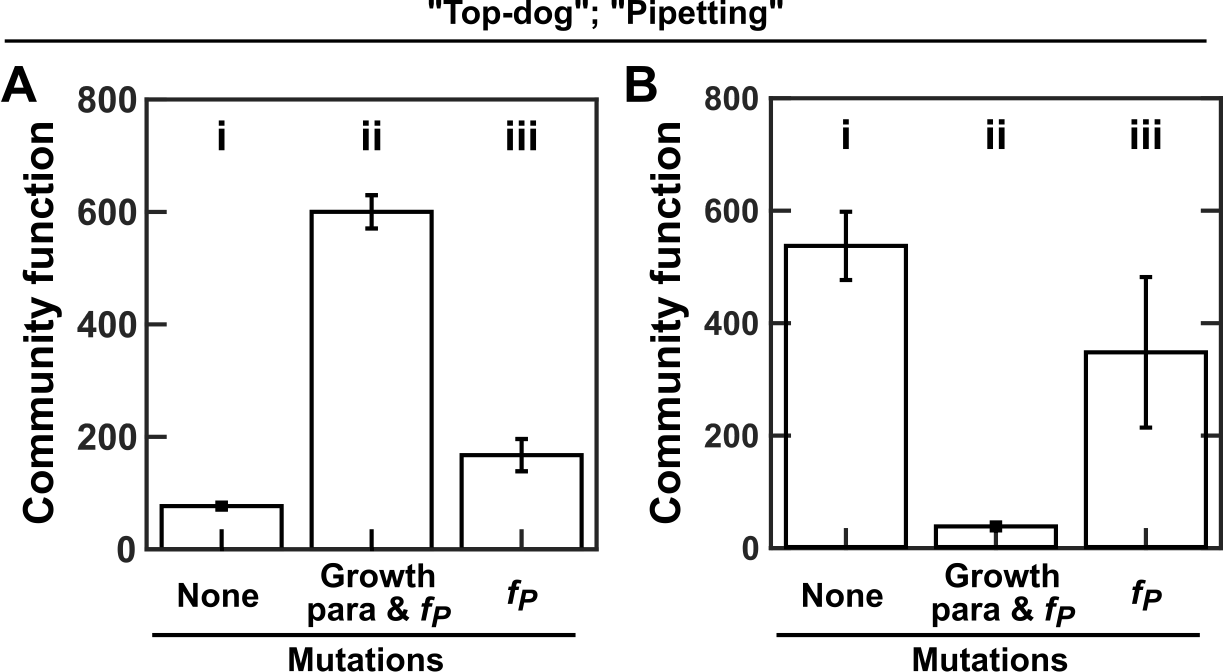
Improving growth can improve or impair community function, depending on evolutionary upper bounds of growth parameters. Plotted here are plateaued community function after 1500 cycles when simulation did or did not allow mutations in growth parameters or *f*_*P*_. (**A**) When evolutionary upper bound for *g_Hmax_* 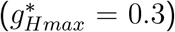 was lower than that of *g_Hmax_* 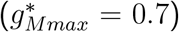, improving growth parameters improved community function. Compared to community function where no mutations were allowed (i), community function improved when growth parameters and *f*_*P*_ were allowed to mutate (ii). Preventing mutations in growth parameters diminished community function improvement (iii). In this case, improved growth of M and H resulted in higher community function. Evolutionary dynamics are shown in Figure S8C. (**B**) When evolutionary upper bound for *g_Hmax_* 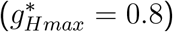 was larger than that of *gHmax* 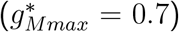, improving growth parameters could decrease community function. Compared to community function where no mutations were allowed (i), community function decreased when growth parameters and *f*_*P*_ were allowed to mutate (ii). Preventing mutations in growth parameters diminished reduction in community function (iii). In this case, improved growth of M and H resulted in lower community function. Evolutionary dynamics are shown in Figure S17A. In **B**, Resource supplied to Newborn communities could support 10^5^ total biomass to accommodate faster growth rate. In both **A** and **B**, community reproduction occurred through volumetric dilution via pipetting, and the top-dog strategy was used. Error bars are calculated form three independent selections.

**Figure S25:**
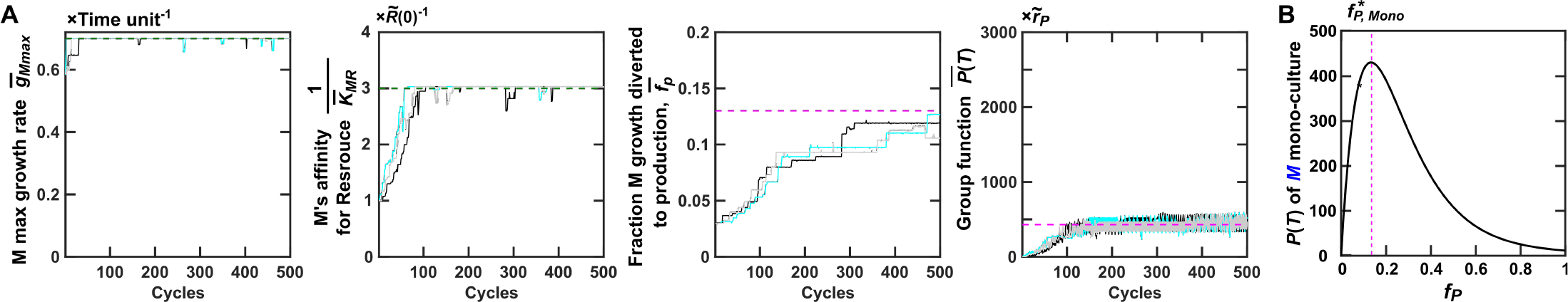
Selection dynamics of M mono-species groups. (**A**) Phenotypes averaged over chosen groups are plotted for 500 selection cycles. Because Byproduct is in excess, *K*_*MB*_ terms are no longer relevant in equations (Figure S4, *R*_*M*_ ≪ *B*_*M*_). Upper bounds of *g*_*Mmax*_ and 1*/K*_*MR*_ are marked with green dashed lines. Magenta lines mark *f*_*P*_ optimal for group function and maximal *P*(*T*) when *g*_*Mmax*_ and 1*/K*_*MR*_ are fixed at their upper bounds and when Byproduct is in excess. (**B**) Suppose that a Newborn M group starts with a single Manufacturer (biomass 1) supplied with excess Byproduct and the same amount of Resource as in a Newborn H-M community (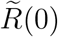 could support 10^4^ M biomass). Then, maximal group function is achieved at 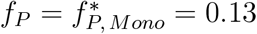 (dashed line), lower than the optimal *f*_*P*_ for the community function 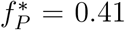 (Figure 2B). Here, the growth parameters of M are all fixed at their evolutionary upper bounds and *P*(*T*) has the unit of 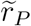.

**Figure S26:**
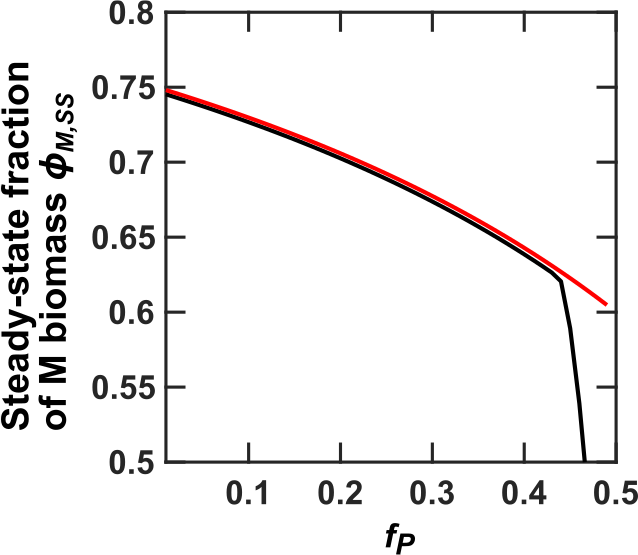
Comparison between the steady-state *ϕ*_*M,SS*_ calculated from Eqs. 6-10 (black curve) and from Eq. 14 (red line).

**Figure S27:**
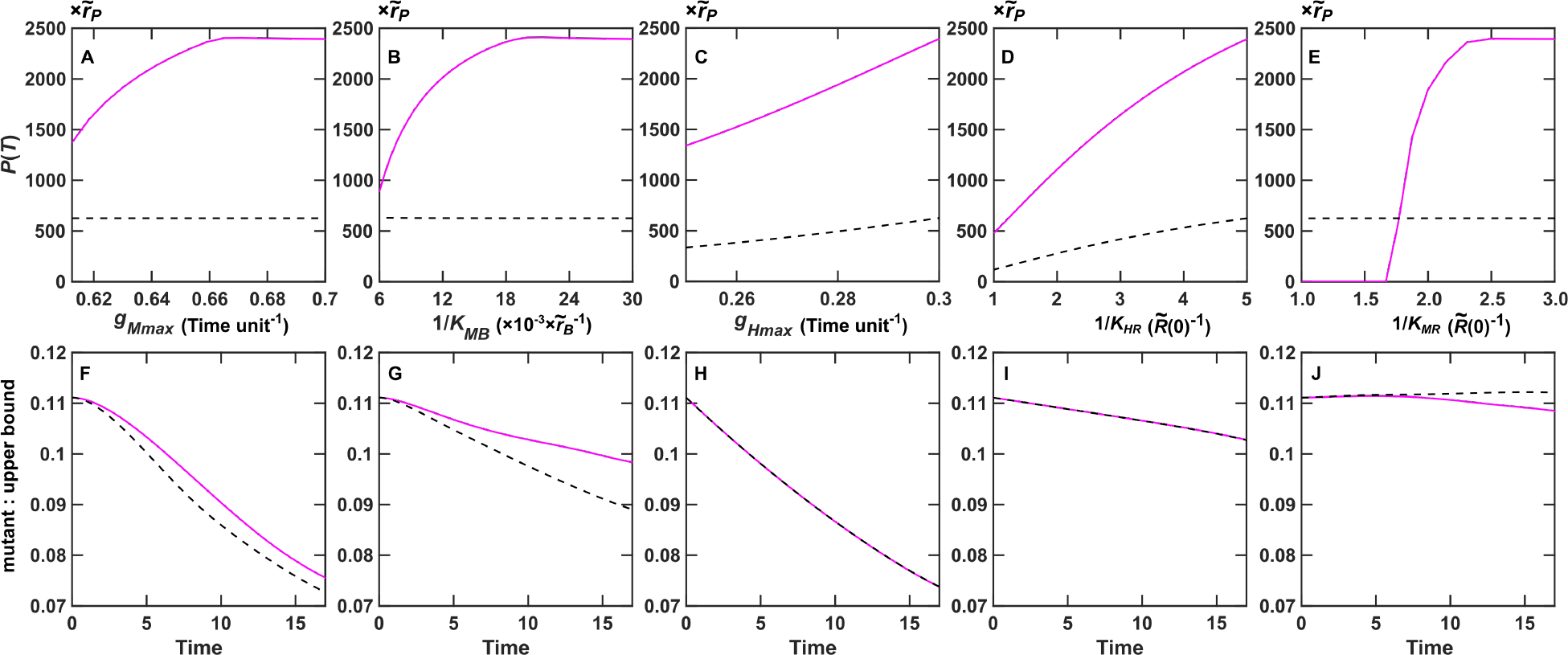
Improved maximal growth rates and nutrient affinities generally, but do not always, improve individual fitness and community function. In all figures, solid and dashed lines respectively represent calculations with 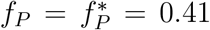 (optimal for community function; Figure 2B) and 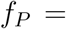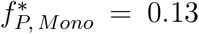 (optimal for M monoculture production when Byproduct is in excess; Figure S25). Except for the indicated growth parameter, all other growth parameters were set to their respective upper bounds. Dynamics within one selection cycle is plotted. (**A**-**D**) Community function increases as the indicated growth parameter increases. For example in (**A**), all growth parameters except for *g*_*Mmax*_ were set to their upper bounds. For each *g*_*Mmax*_, the steady-state *ϕ*_*M,SS*_ was calculated using equations in Methods Section 1. This steady-state *ϕ*_*M,SS*_ was then used to calculate *P*(*T*). (**F**-**I**) The ratio between mutant population (whose indicated growth parameter was 10% lower than the upper bound) and growth-adapted population over maturation time *T* = 17. The decreasing ratio indicates that the mutant has a lower fitness compared to the growth-adapted cells. For example in (**F**), a Newborn community had 70 M and 30 H. 90% of M were growth-adapted and had upper bound *g*_*Mmax*_ = 0.7 (“upper bound”). 10% of M had *g*_*Mmax*_ = 0.63, 10% less than the upper bound (“mutant”). The ratio between “mutant” and “upper bound” cells declined over maturation time, indicating that mutant M cells had a lower fitness. (**E**, **J**) When *f*_*P*_ = 0.13 (black dashed line) but not when *f*_*P*_ = 0.41 (magenta line), increasing M’s affinity for Resource (1*/K*_*MR*_) slightly decreases individual fitness, and barely affects community function.

**Figure S28:**
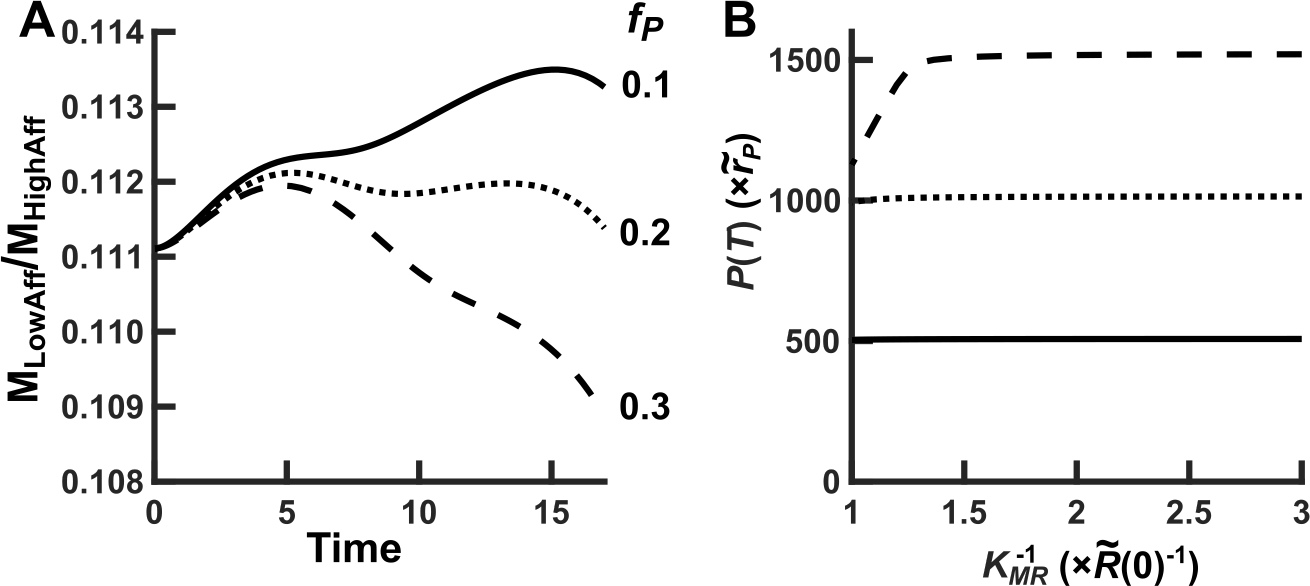
At low *f*_*P*_, M’s lower affinity for Resource can increase its growth rate. **(A)** The ratio between M_LowAff_ (the population size of M with low affinity for Resource 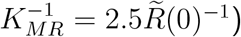 and M_HighAff_ (the population size of M with high affinity for Resource 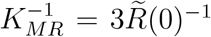 when their *f*_*P*_ is equal to 0.1 (solid line), 0.2 (dotted line) and 0.3 (dashed line) are plotted over one maturation cycle when grown together in the H-M community. **(B)** *P*(*T*) improves over increasing affinity 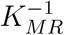 when *f*_*P*_ is 0.1 (solid line), 0.2 (dotted line) and 0.3 (dashed line). The dependence of *P*(*T*) on 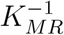 is rather weak for low *f*_*P*_. For example, when 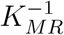 increases from 1 to 3, *P*(*T*) increases by only 2% and 0.6% for *f*_*P*_ = 0.2 and *f*_*P*_ = 0.1, respectively.

**Figure S29:**
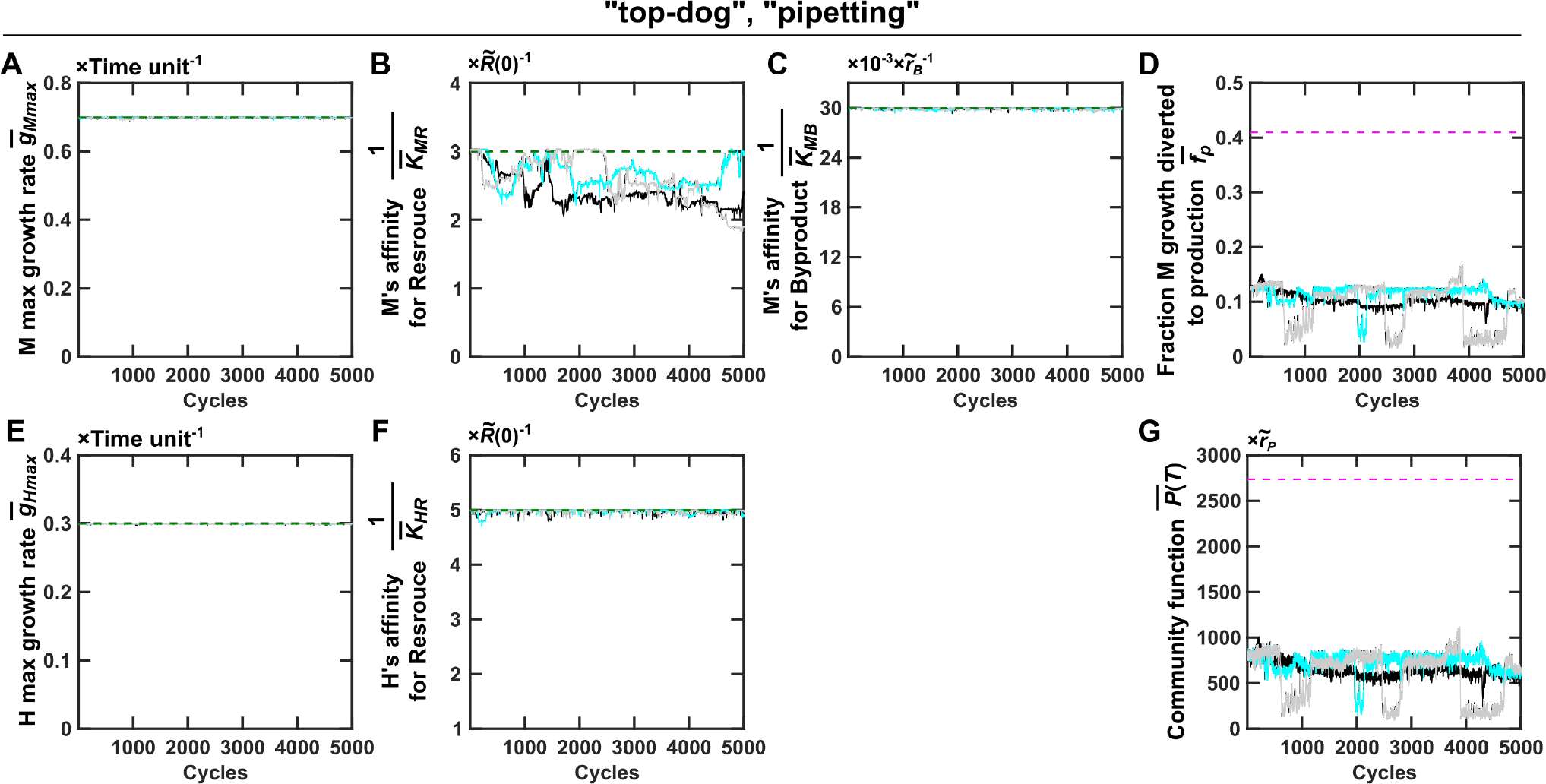
Selection dynamics of communities of mono-adapted H and M when allowing all parameters to vary. In the Newborn communities of the first cycle of community selection, all growth parameters of H and M were at their upper bounds and 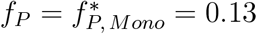 (Figure S25). When we simulated community selection while allowing all growth parameters and *f*_*P*_ to vary, M’s affinity for R 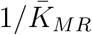 decreased slightly because at low *f*_*P*_ = 0.13, M with a lower affinity for R (lower 1*/K*_*MR*_) has a slightly improved individual fitness (Figure S28). Other growth parameters (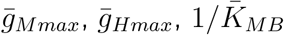 and 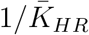) remain mostly constant during community selection because mutants with lower-than-maximal values were selected against by intra-community selection and by inter-community selection (Figure S27). Other legends are the same as Figure S8.

**Figure S30:**
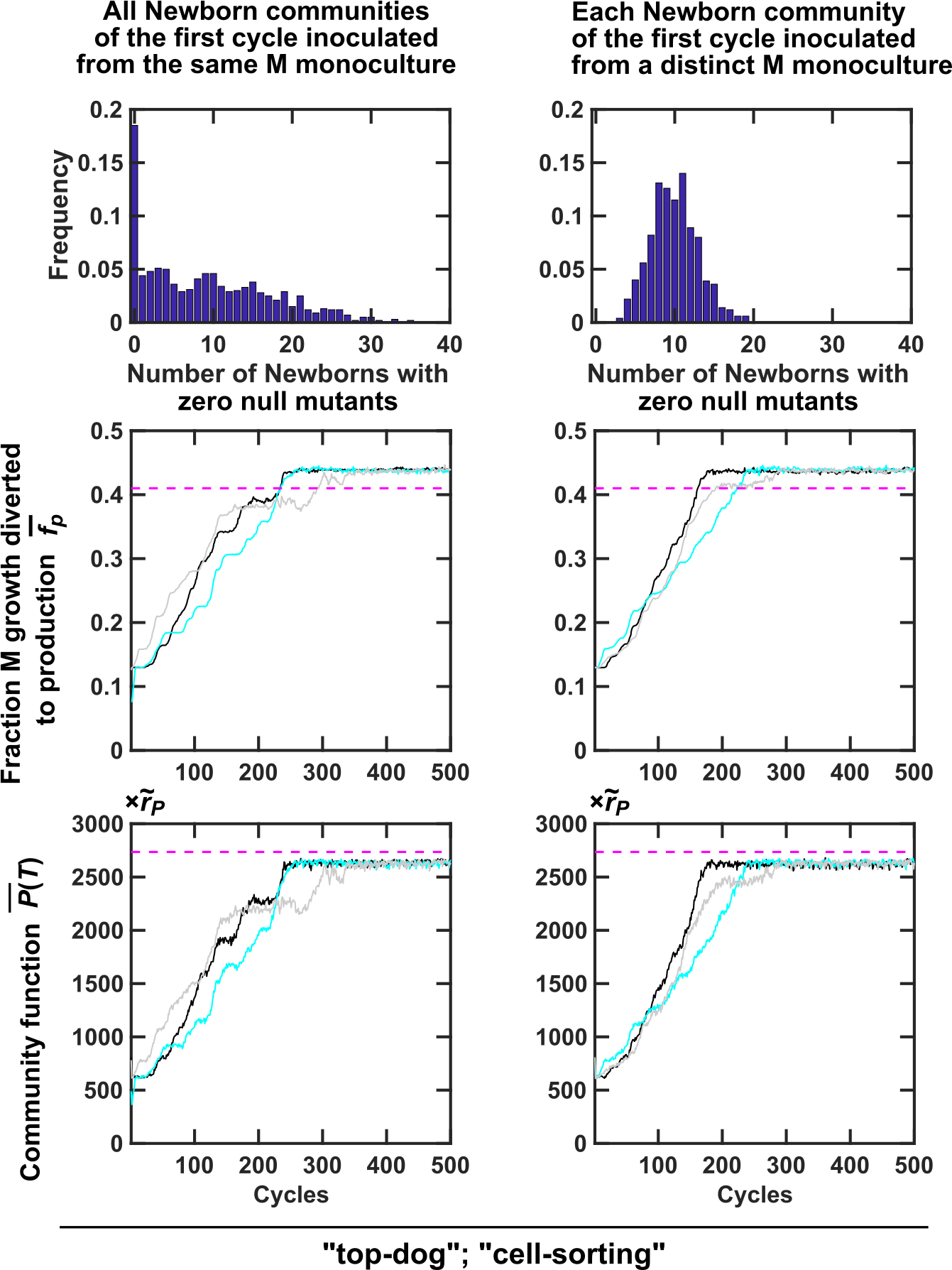
Different methods of pre-growth had limited impact on selection dynamics. (Top Panels) Histograms of the number of Newborn communities free of non-contributor M mutants when Newborn communities of the first cycle were inoculated from a single M monoculture (**Left panel**) or from independently-grown M monocultures (**Right panel**). To generate the histograms, the pre-growth and inoculation process was repeated 100 times. (**Middle and Bottom panel**s) Improvement in 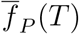 and 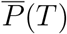 was only slightly slower when Newborn communities from the first cycle were inoculated by the same M monoculture (**Left panel**) than by distinct monocultures (**Right panel**). Here we assumed that each M monoculture grew from a single non-null M cell. This M cell went through ~23 doublings and therefore multiplied into ~10^7^ cells. Every time a non-null M cell divides, the mother and daughter cells can independently mutate and become a null M cell (*f*_*P*_ = 0) at a fixed probability of 10^−3^. If a non-null M cell has *f*_*P*_ = 0.13, then it will grow at a rate 87% of that of a null cell. After ~23 doublings, the M monocultures have on average ~3% null mutants. 60 randomly-chosen M cells from the same monoculture or from distinct monocultures, together with 40 H cells, were used to inoculate each of the 100 Newborns for the first selection cycle. The top-dog strategy was used to choose Adults which were then reproduced via cell-sorting. Black, cyan and gray curves are three independent simulation trials. 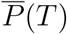 was averaged across the two chosen Adults. 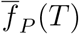 was obtained by first averaging among M within each chosen Adult and then averaging across the two chosen Adults.

